# Growth and adaptation mechanisms of tumour spheroids with time-dependent oxygen availability

**DOI:** 10.1101/2022.04.24.489294

**Authors:** Ryan J. Murphy, Gency Gunasingh, Nikolas K. Haass, Matthew J. Simpson

**Affiliations:** Mathematical Sciences, Queensland University of Technology, Brisbane, Australia; The University of Queensland Diamantina Institute, The University of Queensland, Brisbane, Australia

## Abstract

Tumours are subject to external environmental variability. However, *in vitro* tumour spheroid experiments, used to understand cancer progression and develop cancer therapies, have been routinely performed for the past fifty years in constant external environments. Furthermore, spheroids are typically grown in ambient atmospheric oxygen (normoxia), whereas most *in vivo* tumours exist in hypoxic environments. Therefore, there are clear discrepancies between *in vitro* and *in vivo* conditions. We explore these discrepancies by combining tools from experimental biology, mathematical modelling, and statistical uncertainty quantification. Focusing on oxygen variability to develop our framework, we reveal key biological mechanisms governing tumour spheroid growth. Growing spheroids in time-dependent conditions, we identify and quantify novel biological adaptation mechanisms, including unexpected necrotic core removal, and transient reversal of the tumour spheroid growth phases.

## 1 Introduction

*In vivo* tumours are subject to various types of environmental variability, for example due to fluctuating oxygen and nutrient availability [1–4]. To study cancer progression and develop cancer therapies, tumour spheroid experiments have been successfully and routinely performed for the past fifty years [2, 5–12]. However, tumour spheroid experiments are typically performed in constant environments and focus on the overall size of spheroids [2,13–18]. By experimentally controlling oxygen availability and using mathematical modelling and statistical uncertainty quantification, we develop a new framework to study the impact of external environmental variability on the growth of tumour spheroids and their internal structure. Using our framework we identify and quantify novel biological adaptation mechanisms driven by environmental variability. This work begins to bridge the gap between *in vitro* and in vivo conditions, and lays the foundation for future experimental, mathematical, and statistical spheroid studies.

Oxygen availability is of particular importance since it is vital to the effectiveness of cancer therapies, such as chemotherapy and radiotherapy [1, 19, 20], and can be controlled in spheroid experiments. However, spheroid experiments are typically performed in ambient atmospheric conditions (21% oxygen), sometimes referred to as normoxia [13,16]. In contrast, untreated tumours typically grow in variable hypoxic conditions (0.3-4.2% oxygen) [1,20–22]. While many single-cell studies, and some spheroid studies, explore the role of environmental variability [2, 22–27], oxygen parameters critical to reproduce results are commonly not reported [28].

To visualise spheroid growth in normoxia, hypoxia, and time-dependent oxygen conditions we use fluorescent ubiquitation cell cycle indicator (FUCCI) transduced cell lines and hypoxia markers (Figure 1a-e) [13, 14, 29, 30]: nuclei of cells in gap 1 (G1) phase fluoresce red, shown in magenta for clarity (Figure 1d); nuclei of cells in synthesis, gap 2, and mitotic (S/G2/M) phases fluoresce green (Figure 1d); and, regions of hypoxia are indicated by cyan (Figure 1b,c,e). Spheroids grown in constant normoxia experience three phases of growth (Figure 1a-c,f). In phase (i) spheroids grow exponentially as all cells are able to proliferate, indicated by the presence of cells in the S/G2/M phases throughout the tumour spheroid shown by green (Figure 1a). In phase (ii) cells in the central region of the spheroid arrest in G1 phase while cells at the periphery continue to proliferate resulting in inhibited growth (Figure 1b). This arrested region is thought to arise due spatial differences in nutrient availability, possibly oxygen, and/or a build up of metabolic waste from cells. In phase (iii) the spheroid is characterised by three regions: a central region composed of a necrotic core, 0 < *r* < *R*_n_(*t*); an intermediate region of living but proliferation-inhibited cells, *R*_n_(*t*) < *r* < *R*_i_(*t*); and, a region at the periphery composed of living and proliferating cells, *R*_i_(*t*) < *r* < *R*_o_(*t*) (Figure 1c). In comparison to spheroids grown in normoxia, spheroids grown in hypoxia form their necrotic core earlier, the distance from the edge of the spheroid to the hypoxic region and overall size are smaller (Figure 2).

**Figure 1:**
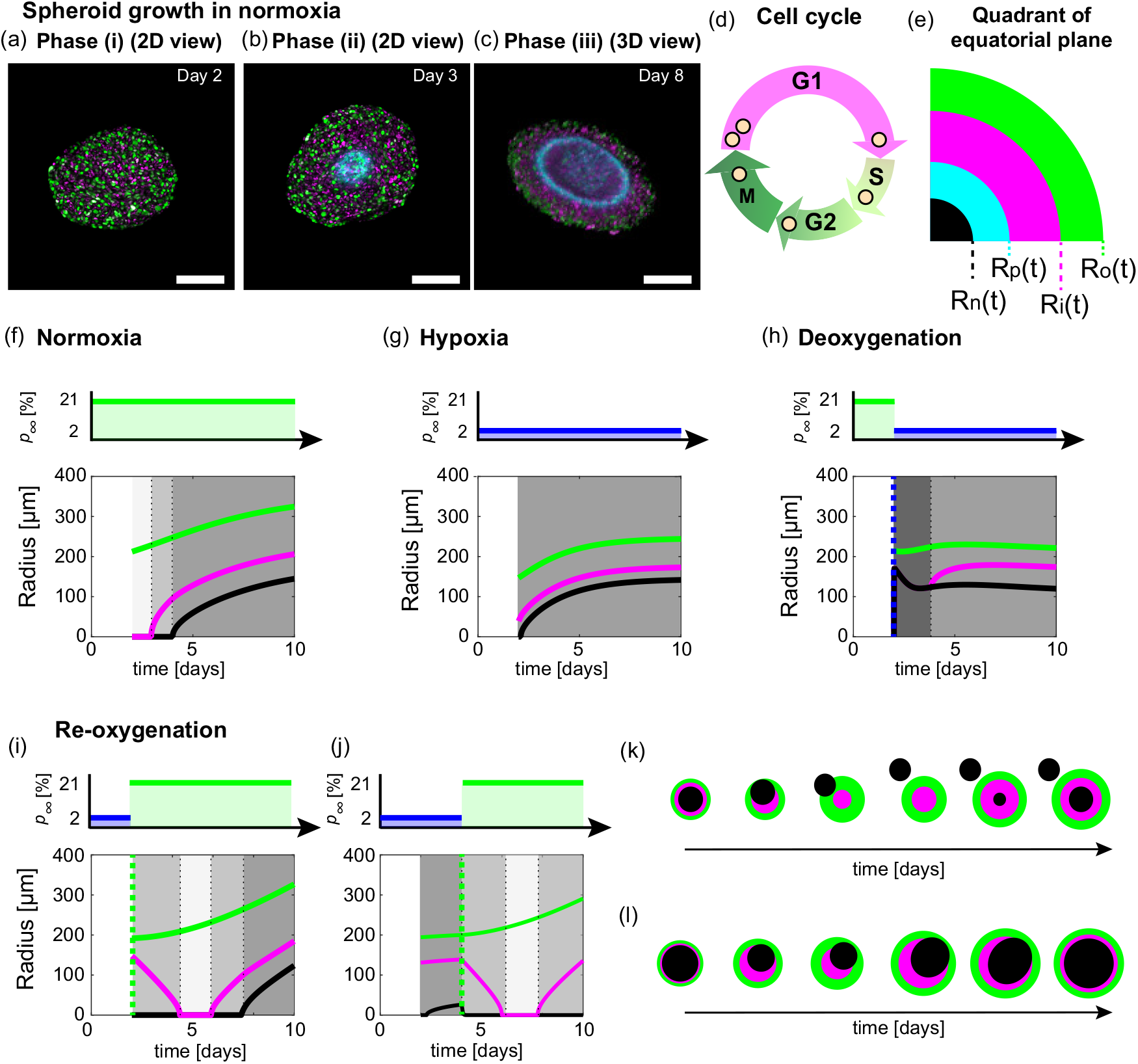
Impact of external environment on the structure of growing tumour spheroids: a focus on oxygen availability. (a-c) Tumour spheroid growth in standard experimental protocols occurs in three phases. Experimental images shown for FUCCI-transduced human melanoma WM983b spheroids grown in normoxia. (a-b) Experimental images of the equatorial plane of spheroids on Day 2 and 3 after seeding. (c) 3D z-stack representation of half of a spheroid on Day 8 after seeding. Scale bars are 200μm. Colours in (a-c) correspond to cell cycle schematic shown in (d): cells in G1 phase (magenta) and cells in S/G2/M phase (green). Pimonidazole staining reveals the hypoxic regions of spheroid (cyan). (e) Schematic for spherically symmetric spheroid structure representing a quadrant of the equatorial plane of a spheroid. Spheroids in normoxia experience three phases of growth, resulting in a spheroid with three regions at later times: a central region composed of a necrotic core, 0 < *r* < *R*_n_(*t*) (black); an intermediate region of living but proliferation-inhibited cells, *R*_n_(*t*) < *r* < *R*_i_(*t*) (magenta); and, a region at the periphery composed of living and proliferating cells, *R*_i_(*t*) < *r* < *R*_o_(*t*) (green). The hypoxic radius, *R*_p_(*t*) (cyan) satisfies *R*_n_(*t*) ≤ *R*_p_(*t*) ≤ *R*_o_(*t*). (f-j) Schematics for oxygen conditions and time evolution of spheroid structure and overall size in (f) normoxia, (g) hypoxia, (h) deoxygenation experiments, and (i-j) re-oxygenation experiments. Note in (i-j) spheroids transiently undergo the growth phases in reverse. Greyscale shading in (f-j) represent growth phases. (k-l) Spheroid schematics showing (k) necrotic core removal and (l) movement of necrotic core without removal.

**Figure 2:**
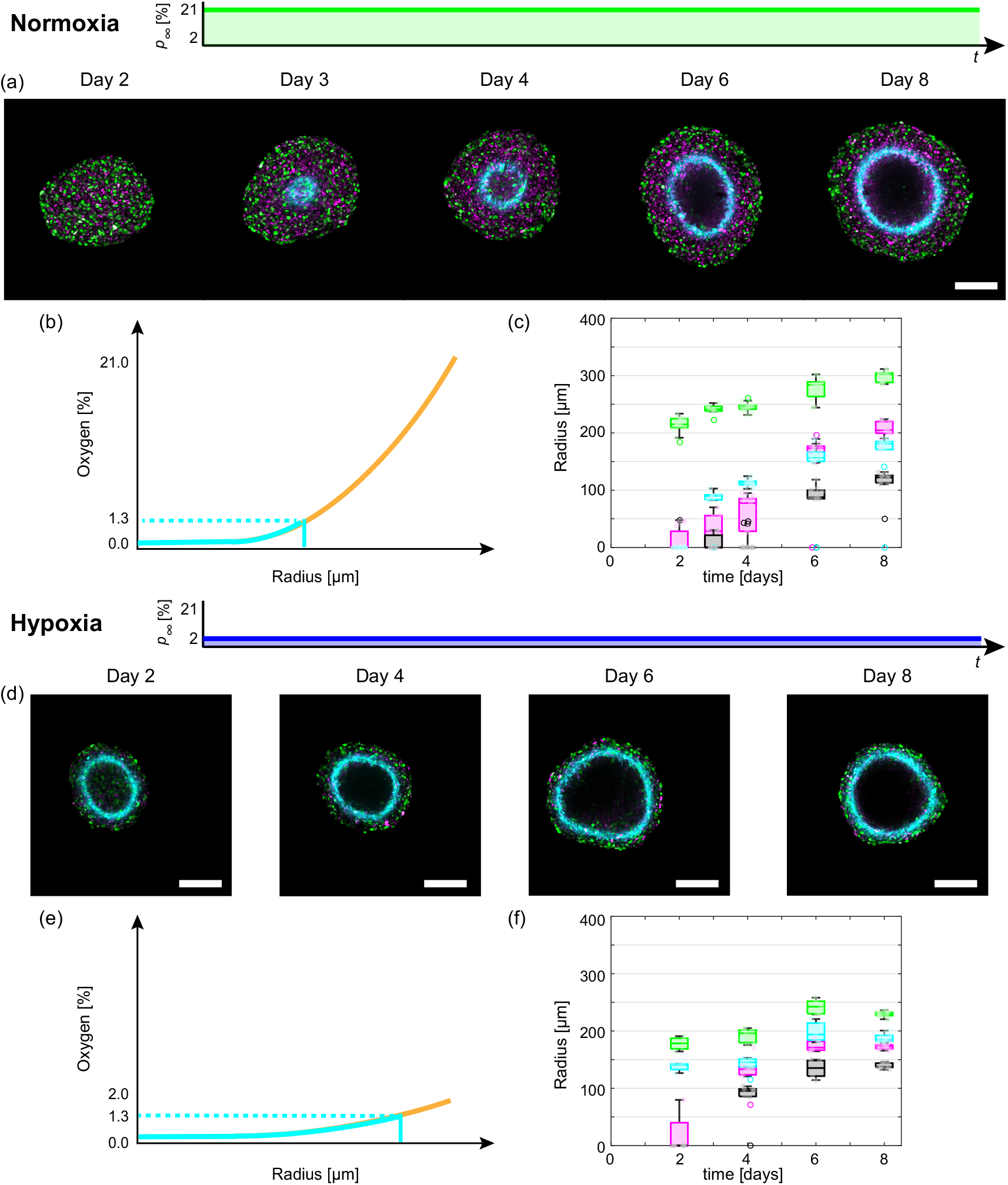
Tumour spheroid growth in normoxia and hypoxia. (a,d) Experimental images of the equatorial plane of FUCCI-transduced WM983b spheroids grown in (a) normoxia (21% oxygen) and (b) hypoxia (2% oxygen). (a) Images shown on Day 2, 3, 4, 6, and 8 after seeding. (d) Images shown in Day 2, 4, 6, and 8 after seeding. Scale bars are 200μm. (b,e) Schematics for oxygen partial pressure within spheroids for (b) normoxia and (e) hypoxia. (c,f) Time-evolution of *R*_o_(*t*) (green), *R*_i_(*t*) (magenta), *R*_n_(*t*) (black), and *R*_p_(*t*) (cyan) for spheroids grown in (c) normoxia, and (f) hypoxia. Note that each spheroid measurement is an end-point measurement.

To investigate environmental variability we perform additional experiments in time-dependent oxygen conditions. In these experiments we observe various tumour spheroid adaptation mechanisms (Figure 1h-l). For instance, in re-oxygenation experiments we discover a novel adaptation process where the necrotic core of the spheroid that has formed prior to re-oxygenation moves within the spheroid and in certain situations exits the spheroid as a single object (Figure 1k, Movie S1). Further, for fifty years tumour spheroid growth has been described by three sequential growth phases but reoxygenation experiments show that spheroids can transiently experience these phases in the reverse order (Figure 1i,j). Other observations from these experiments agree with intuitive expectations, but have not previously been explored nor quantified.

Throughout this study we quantitatively analyse experimental data using mathematical modelling and statistical uncertainty quantification. We start with the seminal Greenspan mathematical model [10, 15, 16, 31]. Greenspan’s model describes the three phases of growth and is relatively simple in comparison to other models [31–34]. This simplicity is a great advantage. We are able to extend the model to analyse environmental variability while retaining physical and biologically insightful interpretations of results. Further, by using parameter identifiability analysis, with both profile likelihood and Bayesian inference approaches, we estimate key biological parameters and reveal biological adaptation mechanisms.

In the following we first analyse spheroid experiments in normoxia and hypoxia. Such experiments demonstrate that Greenspan’s model describes the experimental data remarkably well. Further, that oxygen mechanisms accurately describe the growth and formation of the necrotic core, whereas other mechanisms, possibly waste mechanisms, likely result in the growth and formation of the inhibited region. We then extend the mathematical model to interpret deoxygenation and re-oxygenation experiments, providing quantitative insights to biological adaption mechanisms throughout. We conclude by describing the unexpected behaviours observed in re-oxygenation experiments.

## 2 Results

Here we focus on WM983b spheroids in normoxia, hypoxia and deoxygenation experiments. Similar results for WM793b and WM164 spheroids are discussed in Supplementary Discussion E - F. For re-oxygenation experiments, we compare results from WM983b and WM793b spheroids as we observe a range of behaviours. In Supplementary Discussion F we discuss additional WM164 re-oxygenation experiments.

### 2.1 Oxygen diffusion alone is insufficient to describe spheroid growth

We capture end-point equatorial plane images for spheroids grown in normoxia and hypoxia measuring *R*_o_(*t*), *R*_i_(*t*), *R*_n_(*t*), and *R*_p_(*t*) (Figure 2a,c,d,f) (Methods: Image processing). These measurements are remarkably consistent within each condition and time point (Figures 2c,f). Comparing spheroids grown in normoxia and hypoxia, we observe vastly different tumour growth dynamics and internal structure 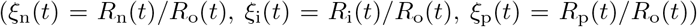, even when comparing spheroids of similar size (Figures 2c,f, S21).

For deeper mechanistic insight we use mathematical modelling and statistical uncertainty quantification to interpret our observations. Specifically, we show that Greenspan’s mathematical model [10] accurately describes spheroid growth in both normoxia and hypoxia. Then using parameter estimation we identify biological mechanisms that differ between normoxia and hypoxia. Key model details are now discussed, for further details see Methods 4.1.1 and Supplementary Discussion C.1. The model assumes each spheroid is spherically symmetric and maintained by cell-cell adhesion or surface tension. The independent variables are time t [days], and radial position, *r* [μm]. Conservation of volume gives an equation describing the time evolution of the outer radius, *R*_o_(*t*) [μm],

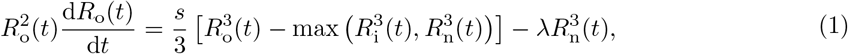

where *s* [day^-1^] is the rate at which cell volume is produced by mitosis per unit volume of living cells, and λ [day^-1^] is the proportionality constant describing the rate at which cell volume is lost from the necrotic core. In these experiments, Equation (1) simplifies as *R*_i_(*t*) ≥ *R*_n_(*t*). This restricts our attention to two interpretations of the model that differ with respect to how *R*_i_(*t*) is defined. In the following discussion, we refer to these interpretations as *hypotheses* and show that hypothesis 2, where oxygen mechanisms drive *R*_n_(*t*) and waste mechanisms drive *R*_i_ (*t*), is more consistent with the spheroids considered in this study.

In hypothesis 1 (Figure 3a), oxygen diffuses with diffusivity, *k* [m^2^ s^-1^], and is consumed by living cells at a constant rate, *α* [m^3^ kg^-1^ s^-1^]. The external oxygen partial pressure is *p*_∞_ [%]. Oxygen diffusion is fast relative to the growth of the spheroid, so that the oxygen partial pressure within the spheroid, *p*(*r*(*t*)) for 0 ≤ r ≤ *R*_o_(*t*), is governed by

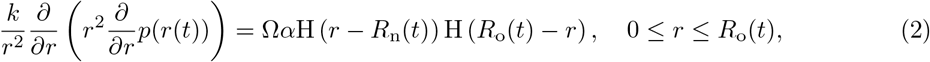

where H(·) is the Heaviside function and Ω [mmHg kg m^-3^] is a conversion constant from volume of oxygen per unit tumour mass to oxygen partial pressure [35]. The inhibited radius, *R*_i_(*t*), is implicitly defined by *p*(*r*(*t*)) = *p*_i_ [%] provided the oxygen partial pressure is sufficiently large (Figure 3a), and *R*_i_(*t*) = 0 otherwise.

**Figure 3:**
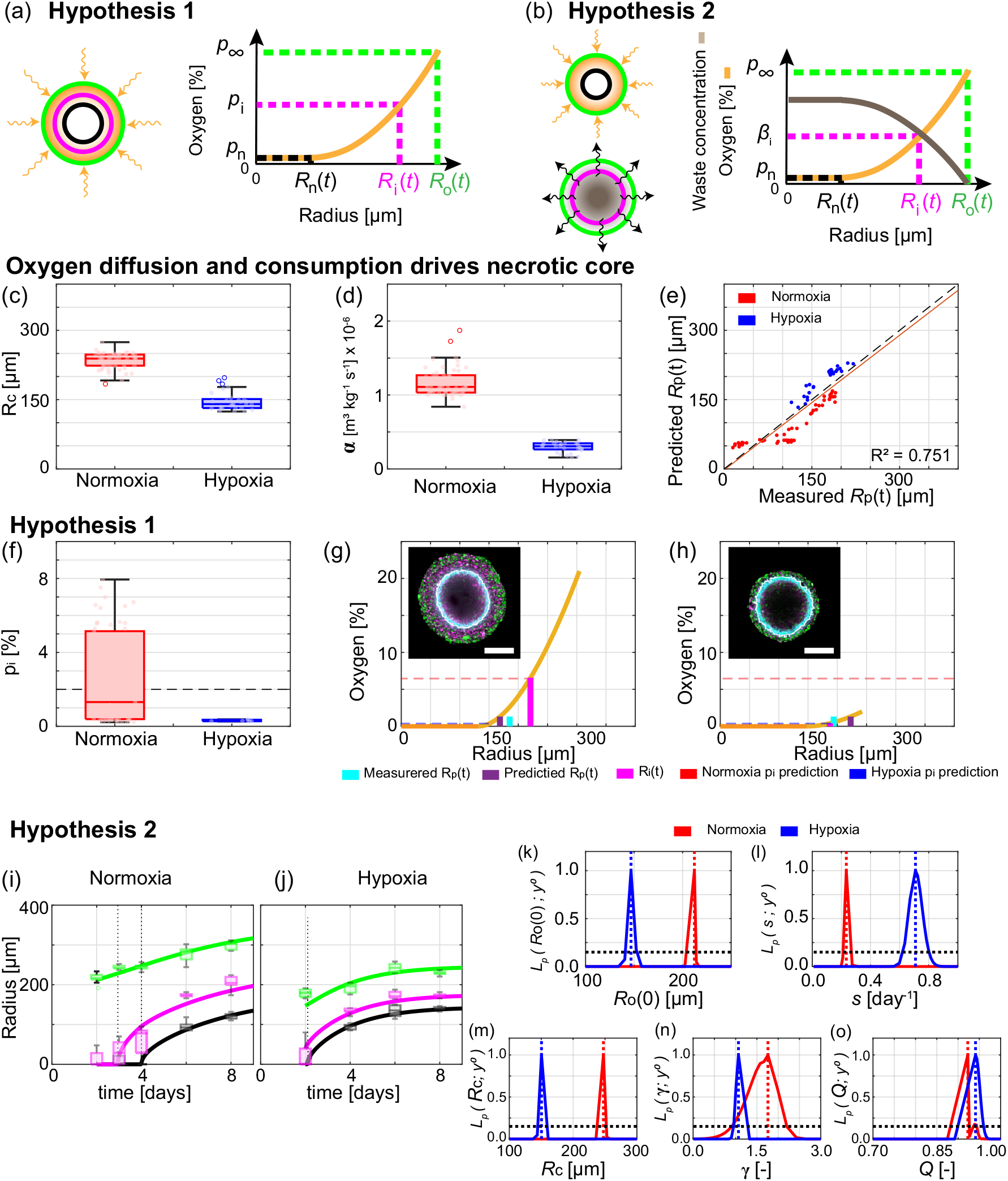
Mechanisms governing tumour spheroid growth in normoxia and hypoxia. (a,b) Schematics for two hypotheses. (a) Oxygen mechanisms describe *R*_i_(*t*) and *R*_n_(*t*). (b) Oxygen mechanisms describe *R*_n_(*t*) and waste mechanisms describe *R*_i_(*t*). (c-e) Oxygen diffusion and consumption describes *R*_n_(*t*). (c) Box chart for estimated outer radius when necrotic region forms, *R*_c_ μm]. (d) Box chart for estimated oxygen consumption rate, α [m^3^ kg^-1^ s^-1^]. (e) Comparison of measured and predicted *R*_p_(*t*) when pimonidazole staining is present. (f) Box chart for estimated oxygen partial pressure defining inhibited region from hypothesis 1, p_i_ [%]. (g-h) Estimated oxygen partial pressure within (g) a spheroid grown in normoxia and (h) a spheroid grown in hypoxia. Insets in (f,g) show FUCCI signal and pimonidazole staining from Day 8 and 6, respectively, with scale bar is 200 μm. In (c-f) data points for normoxia and hypoxia shown in red and blue, respectively. (i,j) Comparison of experimental data with Greenspan’s mathematical model simulated with the maximum likelihood estimates of parameters for (i) normoxia and (j) hypoxia. Time-evolution of outer radius, *R*_o_ (*t*) (green), inhibited radius, *R*_i_(*t*) (magenta), and necrotic radius, *R*_n_(*t*) (black). Data represent an average of twelve spheroids on days 2, 3, 4, 6, and 8 for spheroids grown in normoxia and an average of seven spheroids on days 2, 4, 6, and 8 for spheroids grown in hypoxia (Methods: Experimental methods 4.3). (k-o) Profile likelihoods for (k) initial outer radius, *R*_o_ μm], (l) proliferation rate, *s* [day^-1^], (m) outer radius when necrotic region first forms, *R*_c_ μm], (n) dimensionless parameter relating proliferation rate and mass loss from necrotic core, *γ* [-], and (o) dimensionless parameter relating oxygen and waste mechanisms, *Q* [-] (Methods 4.1.1).

In hypothesis 2 (Figure 3b), diffusible metabolic waste is produced by living cells at a constant rate per unit volume, *P* [mol μm^-3^ day^-1^], and diffuses with diffusivity *κ* |μm^2^ day^-1^]. Waste diffusion is fast relative to the growth of the spheroid, so the waste concentration within the spheroid, *β*(*r*(*t*)) [mol μm^-3^], is governed by

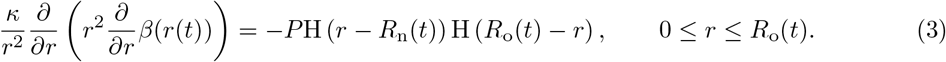

The inhibited radius, *R*_i_(*t*), is implictly defined through *β*(*r*(*t*)) = *β_i_* [mol μm^-3^| provided the waste concentration is sufficiently large (Figure 3b), and *R*_i_(*t*) = 0 otherwise.

Both hypothesis 1 and 2 assume that *R*_n_(*t*) is implicitly defined by *p*(*R*_n_(*t*)) = *p*_n_ provided the oxygen partial pressure is sufficiently small, and *R*_n_(*t*) = 0 otherwise. Informed by experimental results [35], we set *p*_n_ = 0 [%].

Analysis of the model provides an analytical expression for the time when the inhibited region forms (Equation 7.1). For hypothesis 2, the inhibited region is predicted to form at the same time independent of oxygen dynamics, provided the spheroids grown in normoxia and hypoxia are initially the same size. This appears consistent with results in Figure 2a-d where the inhibited region has formed on day 2 for both conditions. In contrast, with hypothesis 1 the time to form the inhibited region depends on oxygen mechanisms and specifically *p*_∞_, but without knowledge of additional parameters further insights are unclear.

By incorporating statistical uncertainty quantification to estimate parameters of the model we gain further mechanistic insights. A key assumption common to hypothesis 1 and 2 is that oxygen diffusion and consumption drives the time evolution of *R*_n_(*t*). We test this assumption directly by analysing radius measurements of each spheroid at each time point independently [35]. Using measurements of *R*_o_(*t*) and *R*_n_(*t*) we estimate: the outer radius when the necrotic region first forms, *R*_c_; α; and *R*_p_(*t*) (Figure 3c-d, Supplementary Discussion D.1.1). Image processing to measure *R*_p_(*t*) is more challenging than for *R*_o_(*t*) due to gradients in the pimonidazole signal (Supplementary Discussion B). However, after careful image processing we find good agreement between experimentally measured and predicted values of *R*_p_(*t*) (*R*^2^ = 0.751, Figure 3e). This approach allows us to estimate the oxygen partial pressure within each spheroid at each time point (Figure 3g-h). From the results in Figures 3e,g,h, S22 we conclude that oxygen of diffusion alone is a reasonable and sufficient mechanism to describe the formation of the necrotic core in WM983b spheroids [35]. Furthermore, these results suggest that the following assumptions are reasonable: *p*_n_ = 0; a constant oxygen consumption rate within the spheroid; spherical symmetry; oxygen at the edge of the spheroid can be approximated with the oxygen settings on the incubator.

Given that *R*_n_(*t*) is reasonably described by oxygen mechanisms we now examine hypothesis 1. Hypothesis 1 assumes that oxygen mechanisms alone drive the time evolution of *R*_i_(*t*). We then estimate *p*_i_ using the estimated oxygen partial pressure within each spheroid, *p*(*r*(*t*)) for 0 < *r* < *R*_o_(*t*), measurements of *R*_i_(*t*) and the definition *p*(*R*_i_(*t*)) = *p*_i_ (Figure 3a,g,h). Estimates of *p*_i_ are consistently larger for spheroids grown in normoxia compared to spheroids grown in hypoxia (Figure 3f). This is inconsistent with hypothesis 1. Specifically, results from normoxia suggest that spheroids grown in hypoxia should have larger inhibited regions than experimentally measured (Figure 3g,h). Similarly, results from spheroids grown in hypoxia suggest that spheroids grown in normoxia should have smaller inhibited regions than experimentally measured. These results provide strong evidence to suggest that oxygen alone is insufficient to describe the formation of the inhibited region across multiple oxygen conditions for this cell line, consistent with results for other cell lines [17].

To test hypothesis 2, which assumes that waste mechanisms drive the time evolution of *R*_i_(*t*), we first analyse measurements of each spheroid at each time point independently. Experimentally measuring waste within spheroids is challenging, so we estimate the waste concentration within each spheroid and use measurements of *R*_i_(*t*) to estimate the outer radius when the inhibited region first forms, 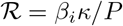 (Figure S23). We observe that 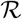 is larger for spheroids grown in normoxia than hypoxia, which may be due to changes in *β_i_* or *P* or *κ* (Figure S23). These results do not provide sufficient evidence to reject hypothesis 2.

To test whether hypothesis 2 is reasonable we estimate model parameters for spheroids grown in normoxia and hypoxia. Specifically, we estimate the five key parameters: Θ_*g*_ = (*R*_o_(0), *s, R*_c_, *γ, Q*), where *γ* = λ/*s* [-] and *Q* [-] are dimensionless quantities (Methods 4.2). Simulating the model with the maximum likelihood estimate (MLE), 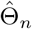, for normoxia shows good agreement with the experimental data from spheroids grown in normoxia (Figure 3i). Similarly, simulating the model at 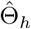 for hypoxia shows good agreement with the experimental data from spheroids grown in hypoxia (Figure 3j). These results suggest the model accurately captures the dynamics of tumour spheroid growth.

Alongside the point estimates 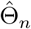 and 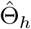, we are interested in forming approximate 95% confidence intervals for each of the parameters. To perform this analysis we employ profile likelihood analysis (Methods 4.2). All profile likelihoods computed here are narrow and well-formed around a single central peak, the MLE, indicating that parameters are identifiable and that a relatively narrow range of parameters give a similar match to the data as the MLE (Figure 3k-o) [16]. Approximate 95% confidence intervals are obtained from these profile likelihoods for each parameter. The profile likelihoods for the initial outer radius, *R*_o_(0), do not overlap and agree with observations that *R*_o_(0) is smaller for spheroids grown in hypoxia than normoxia (Figure 3k). The profile likelihood for *s* interestingly estimates that the rate of cell proliferation per unit volume is faster in hypoxia than normoxia (Figure 3l). This result may seem surprising as a simplistic assumption would be that less oxygen results in less proliferation. However, our result is consistent with observations from other cell lines where an intermediate level of hypoxia encourages more proliferation than normoxia [36–38]. Profile likelihoods for *R*_c_ are consistent with estimates of *R*_c_ obtained by analysing spheroid measurements independently, a good consistency check for the two methods (Figure 3c,m). The other profile likelihoods for *γ* and *Q* overlap suggesting that these parameters are consistent across normoxic and hypoxic conditions (Figure 3n,o). Posterior densities and prediction intervals, estimated using Bayesian inference, also show good agreement with results here and the experimental data (Figure S24). Similar results hold for the other two cell lines (Supplementary Discussion E-F).

In the remainder of this study, we take the most fundamental approach and proceed with Greenspan’s mathematical model and interpret the governing mechanisms with hypothesis 2. We note that other biological mechanisms may also be relevant. However, as the model already appears to capture the key dynamics (Figure 3i,j) we wil avoid overcomplicating the model. Furthermore, in the following analysis of deoxygenation and re-oxygenation experiments we necessarily extend the model.

### 2.2 Adaptation to deoxygenation

The mechanisms underlying how tumour spheroids adapt to time-dependent external environments is unclear. Here, we perform a series of deoxygenation experiments, where spheroids grown in normoxia are transferred to hypoxic conditions at *t* = *t_s_* [days]. Analysing spheroid snapshots reveals how spheroids, and in particular their internal structure, adapt. Extending Greenspan’s mathematical model and using parameter estimation, we identify and quantify key biological adaptation mechanisms.

In the deoxygenation experiments we set *p*_∞_ = 21 [%] for 0 < *t* ≤ *t*_s_ [days], *p*_∞_ = 2 [%] for *t*_s_ < *t* < 8 [days], and *t*_s_ = 2 [days] (Figure 4a). At *t*_s_ = 2 [days] all spheroids are in phase (i) of growth with proliferating cells throughout and no inhibited or necrotic region (Day 2 of Figure 4b). At *t*_s_ + 1 [days] the FUCCI signal in the central region of the spheroid is blurred, relative to the signal at the periphery, indicating dying and dead cells (Day 3 of Figure 4b, Figure S20). Therefore, we identify this central region as the necrotic core (Supplementary Discussion B). Experimental images at later times show that *R*_n_(*t*), *R*_i_(*t*) and *R*_p_(*t*) continue to increase but at a much slower rate (Days 4-8 of Figure 4b,c). Throughout the experiment *R*_o_(*t*) remains approximately constant (Days 2-8 of Figure 4b,c) confirming that the most important changes involve the internal structure and not the overall spheroid size. Further, *ξ_n_*(*t*), *ξ_i_*(*t*), and *ξ*_p_(*t*) approach values observed at late times for spheroids grown in hypoxia (Figure 4c, S25).

**Figure 4:**
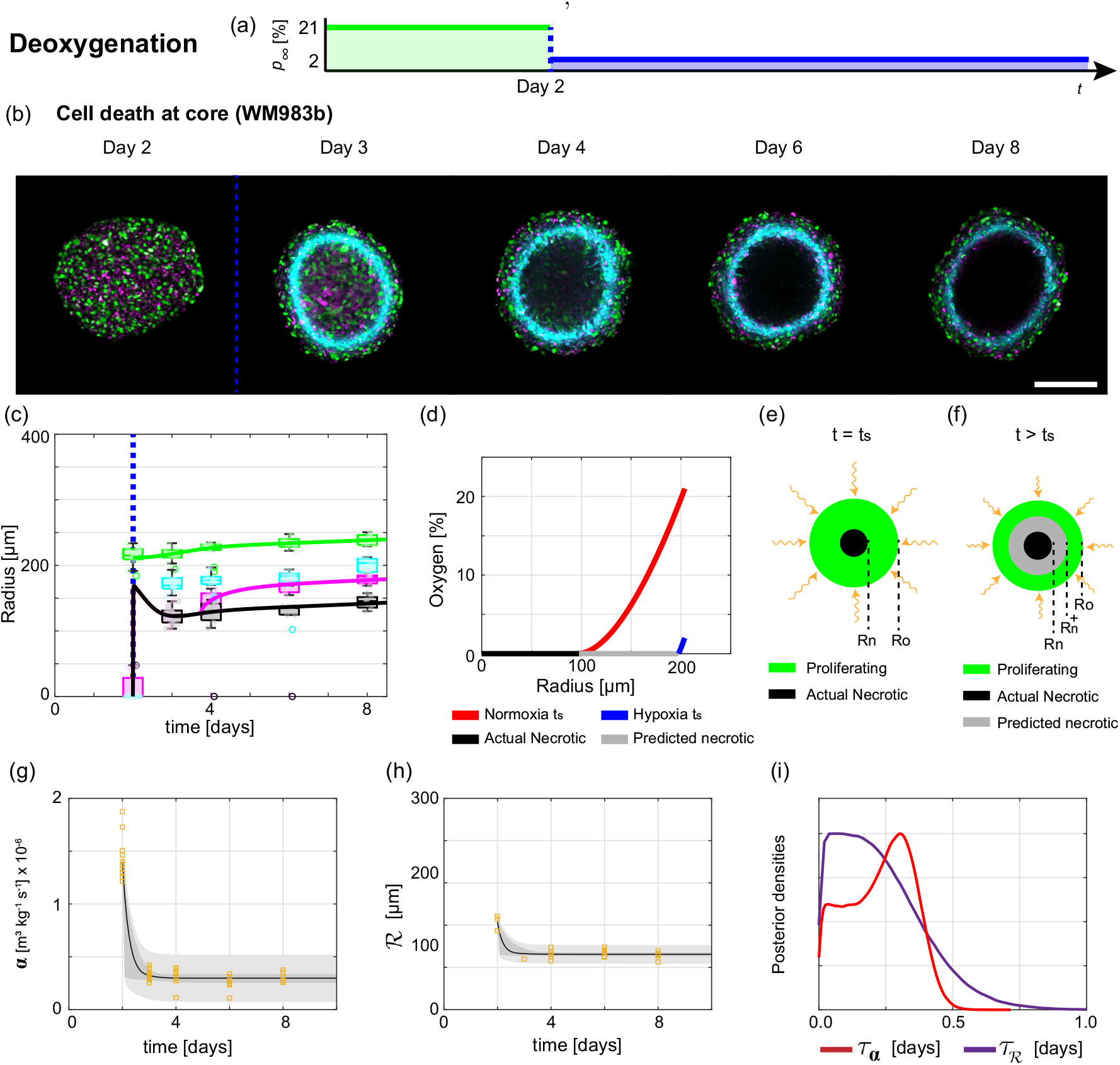
Analysis of deoxygenation experiments reveals tumour spheroid adaptation mechanisms. (a) Schematic for deoxygenation experiment, where the external oxygen environment switches from normoxia to hypoxia at *t*_s_ = 2 [days]. (b) Experimental images of the equatorial plane of WM983b spheroids on Days 2, 3, 4, 6, and 8 after seeding. Scale bar is 200 μm. Colours in (b) correspond to cell cycle schematic shown in Figure 2(c) and hypoxic regions are shown by pimonidazole staining (cyan). (c) Time evolution of outer radius *R*_o_(*t*) (green), inhibited radius *R*_i_(*t*) (magenta), hypoxic radius *R*_p_(*t*) (cyan), and necrotic radius *R*_n_(*t*) (black). Blue dashed lines in (b-c) indicate *t*_s_. (d) Oxygen partial pressure within a spheroid estimated at *t*_s_ under normoxia (red) and hypoxia (blue). (e-f) Immediately after *t*_s_ the predicted necrotic radius, 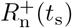, is greater than the actual necrotic radius, *R*_n_(*t*_s_). Cells in region 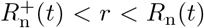 die and increase the volume of the necrotic core. (g) Estimates of *α*(*t*) from experimental data compared to prediction intervals generated from the mathematical model. (h) Estimates of the outer radius when the inhibited region forms, 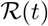, from experimental data compared to prediction intervals generated from the mathematical model. In (g-h) experimental data shown as orange squares, mathematical model simulated with the posterior means of the parameters (black), 50% posterior region of prediction interval (dark grey), and 97.5% posterior region of prediction interval (light grey). (i) Posterior density estimates for adaptation timescales *τ_α_* (dashed) and 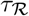 (solid).

To interpret these deoxygenation experiments we extend Greenspan’s mathematical model [10]. We assume that the change in *p*_∞_ at *t*_s_ is instantaneous, which is reasonable since the switch from normoxia to hypoxia requires only 1-2 minutes when the spheroids are transferred between incubators. This time is very short in comparison to the duration of the experiment and time interval between data points. Similarly, we assume that the oxygen partial pressure within the spheroid adapts to the change in *p*_∞_ instantaneously, which is reasonable since oxygen takes approximately 10 seconds to diffuse across a distance of 100 μm [10]. Then we estimate the oxygen partial pressure within the spheroid at *t*_s_ under normoxic and hypoxic conditions (Methods 4.1.2, Figure 4d). Immediately after *t*_s_ the predicted necrotic radius, denoted 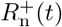 and implicitly defined by 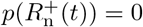, is greater than the actual necrotic radius, *R*_n_(*t*), specifically 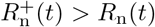 (Figure 4d-f).

Before considering the region 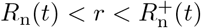, recall that parameter estimates from spheroids grown in normoxia and hypoxia differ. Specifically, *α* (Figure 3d), λ = *γs* (Figure 3l,n), *s* (Figure 3l), and 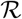 (Figure S23) are all different. Therefore, we expect that these parameter values will evolve in time after *t*_s_. To account for such changes we define the following, for *t* ≥ *t*_s_,

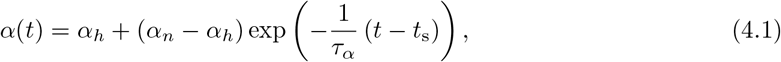

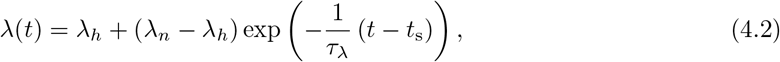

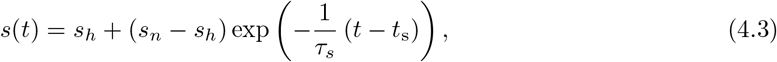

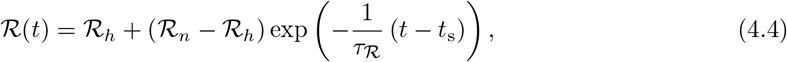

where *τ_α_* [days], *τ*_λ_ [days], *τ_s_* [days], and 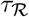 [days] denote timescales of adaptation for *α, λ, s*, and 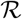, respectively. Further, the new constants in Equation (4) with subscripts *n* and *h*, for example *α_h_* and *α_n_*, represent parameter estimates from spheroids grown in normoxia and hypoxia, respectively. The other parameters (*k*, Ω, *κ*), are assumed to be constants. Hence, 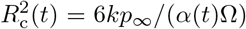 [μm^2^], 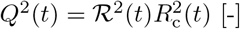, and 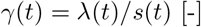 are functions of time.

In the region 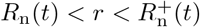 we assume cells die and increase the size of the necrotic core at a rate 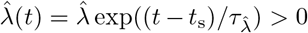 [day^-1^] per unit volume (Figure 4f). Conservation of volume for the necrotic core at time *t, V*_n_(*t*), gives (Supplementary Discussion C.2.1)

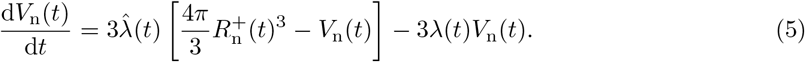

Volume is converted to radius for comparison with experimental data using 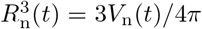. At later times the term involving 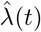 dominates the right hand side of Equation (5) and *R*_n_(*t*) tends to 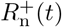. Equation (1), obtained by conservation of volume arguments, remains valid by including the time dependence in *s*(*t*) and λ(*t*). At *t*_s_ there is no immediate change in the waste concentration within the spheroid and so no immediate change to *R*_i_(*t*). However, as 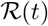 changes with time (Equation (4.4)) the waste concentration within the spheroid and *R*_i_(*t*) evolve over time in part directly due to deoxygenation.

In this new mathematical model (Equations (8.1)-(8.11)) there are fifteen parameters 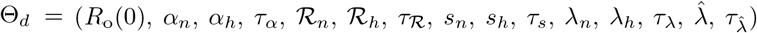. However, using Bayesian inference for parameter estimation the biological adaptation mechanisms become clearer (Methods 4.2). We identify fast adaptation to deoxygenation in *α*(*t*) with mean *τ_α_* = 0.23 [days] and for 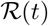 with mean 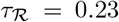 [days] (Figures 4g-j). Furthermore, simulating the new deoxygenation mathematical model we find good agreement with the experimental measurements of *R*_o_(*t*), *R*_n_ (*t*), and *R*_i_ (*t*) (Figure 4c, S26). Therefore, our new mathematical model provides a mechanistic description to the observed growth dynamics in the variable external environment and appears to capture key adaptation mechanisms.

### 2.3 Adaptation to re-oxygenation

We also perform re-oxygenation experiments, where spheroids grown in normoxia are transferred to hypoxic conditions at time *t*_s_. These re-oxygenation experiments exhibit a range of unexpected biological adaptation mechanisms for each cell line that appear to depend on: *t*_s_; spheroid size at re-oxygenation, *R*_o_(*t*_s_); and necrotic core fraction at re-oxygenation, *ξ*_n_(*t*_s_) = *R*_n_(*t*_s_)/*R*_o_(*t*_s_).

First we focus on slower growing WM793b spheroids and *t*_s_ = 2 [days] (Figure 5a,c). In hypoxic conditions prior to re-oxygenation, experimental images show a large hypoxic region (Day 2 of Figure 5a,c,g). However, after re-oxygenation at *t*_s_ + 1 [days] there is no hypoxic region (Day 3 of Figure 5c,g). Spheroid growth after deoxygenation appears to progress similar to spheroids that are grown in normoxia throughout (Days 3-8 in Figure 5c,g).

**Figure 5:**
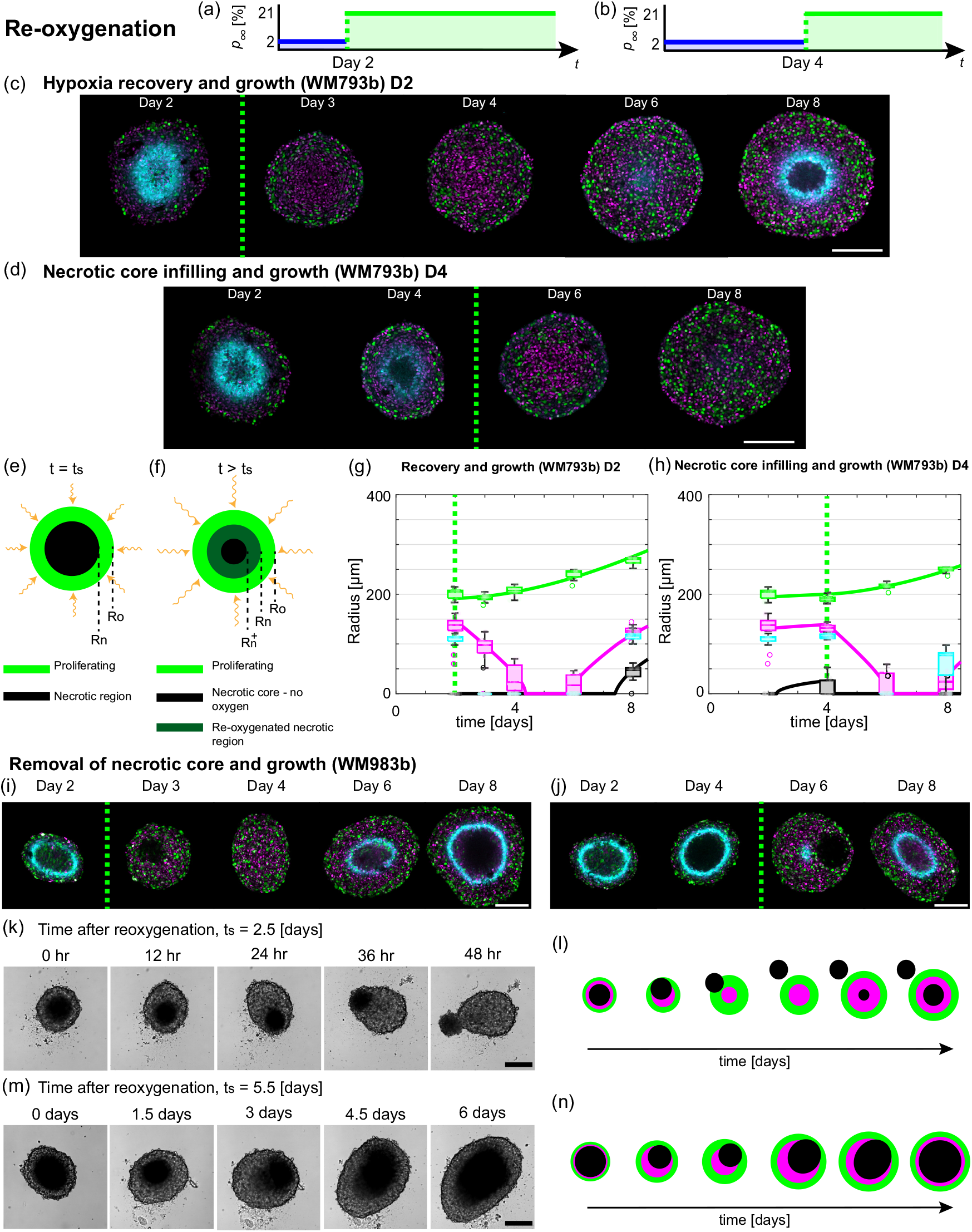
Tumour spheroids exhibit a range of adaption mechanisms in response to re-oxygenation. (a,b) Schematics for re-oxygenation experiments, where the external oxygen environment switches from hypoxia to normoxia at (a) *t*_s_ = 2 [days], and, (b) *t*_s_ =4 [days]. (c,g) Hypoxia recovery and growth for WM793b cell line with *t*_s_ = 2 [days] with (c) experimental images, and (g) radial measurements. (d,h) Necrotic core infilling and growth for WM793b cell line with *t*_s_ = 4 [days] with (d) experimental images, and (h) radial measurements. Green dashed line in (c,d,g,h,i,j) indicate *t*_s_. Colours in (g,h) *R*_o_(*t*) (green), *R*_i_(*t*) (magenta), *R*_n_(*t*) (black), and *R*_p_(*t*) (cyan). (e,f) Schematics for tumour spheroid structure due to re-oxygenation at *t*_s_. (i-l) Removal of necrotic core and growth for WM983b cell line. (i) Confocal images for experiment with *t*_s_ = 2 [days]. (j) Confocal images for experiment with *t*_s_ = 4 [days]. (k) Bright-field images for experiment with *t*_s_ = 2.5 [days]. (l) Schematic for removal of necrotic core and growth in WM983b cell line. (m) Bright-field images for experiment with *t*_s_ = 5.5 [days]. (n) Schematic for movement of necrotic core and growth in WM983b cell line. Colours in (c,d,i,j) correspond to cell cycle schematic shown in Figure 2(c) and hypoxic regions are shown by pimonidazole staining (cyan). Scale bars in (c,d,i,j,k,m) are 200μm.

For WM793b spheroids and *t*_s_ = 4 [days], a necrotic core forms before ts (Day 4 in Figure 5d,h). However, after deoxygenation, at *t*_s_ + 2 [days] there is no necrotic core (Day 6 of Figure 5d,h). At *t*_s_ + 4 [days] spheroids are either in phase (i) or phase (ii) (Day 8 of Figure 5d,h). While traditional tumour spheroid experiments progress through phase (i), (ii), and (iii) sequentially, these experiments show that spheroids can transition transiently through the growth phases in reverse order before subsequently growing in the usual order.

To interpret the WM793b re-oxygenation experiments we proceed analogously to the deoxygenation experiments. We extend Greenspan’s mathematical model to account for differences in parameter estimates between normoxia and hypoxia. Estimating the oxygen partial pressure within the spheroid at *t*_s_, the predicted necrotic core, denoted 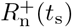, is smaller than the actual necrotic core, *R*_n_(*t*_s_), provided *R*_n_(*t*_s_) > 0. In the region 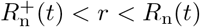 where the necrotic core is now supplied with oxygen, we assume that size of the necrotic core decreases at a rate 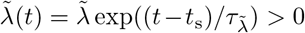 [day^-1^] per unit volume. We assume that a fraction, 0 ≤ *v* ≤ 1, of the volume lost from the necrotic core recovers from the harsh oxygen conditions and increases the population of living cells, and the remaining volume lost from the necrotic core diffuses out of the spheroid and does not influence *R*_i_(*t*). Using Bayesian inference we estimate the seventeen model parameters, 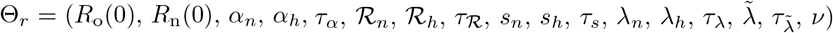 and simulate the mathematical model. We observe good agreement with the experimental data suggesting our new re-oxygenation mathematical model captures key mechanisms underlying adaptation and growth (Figures 5g,h, S28, S29).

Results for WM983b spheroids are unexpected. We hypothesised, based on the exploration of the mathematical model, that spheroid growth dynamics may occur in reverse as observed for WM793b spheroids. However, we did not anticipate that WM983b spheroids would lose their symmetrical internal structure and necrotic core due to re-oxygenation (Figures 5i-n, Movies S1,S2). The WM983b experiments are performed at four different re-oxygenation times *t*_s_ = 2, 2.5,4, and 5.5 [days] (Figures 5i,j,k,m). For all experiments, spheroids at *t*_s_ prior to re-oxygenation are in phase (iii). For *t*_s_ = 2, confocal microscopy reveals that at *t*_s_ + 1 [days] there is a necrotic region but it is not at the centre of the spheroid (Day 3 of Figure 5i). At later times there is no necrotic region and growth proceeds analogous to spheroids in normoxia. Similarly, for *t*_s_ = 4 (Figure 5j).

To explore this unusual behaviour for *t*_s_ = 2.5 [days] and *t*_s_ = 5.5 [days] we perform experiments using the IncuCyte S3 live cell imaging system (Sartorius, Goettingen, Germany) and obtain hourly bright-field images after re-oxygenation. In Figure 5k and Movie S1 with *t*_s_ = 2.5 [days], initially the necrotic core of the spheroid is visible as a dark central region. At later times the necrotic core is located closer to the edge of the spheroid and the radially symmetric internal structure is lost (12, 24 and 36 hours after re-oxygenation in Figure 5k). At *t*_s_ + 2 [days] the necrotic core appears to have exited the spheroid as a single object (48 hours after re-oxygenation in Figure 5k). Tracking the position of the necrotic core relative to the spheroid suggests the necrotic core moves randomly (Supplementary Discussion D.3.1).

We observe similar behaviour for WM983b spheroids with *t*_s_ = 5.5 [days] (Figure 5m, Movie S2). However, likely due to the larger *R*_o_(*t*_s_) and *ξ*_n_ (*t*_s_) here, the necrotic core is close to the edge of the spheroid but does not exit as a single object (1.5 and 3 days after *t*_s_ in Figure 5m). Instead as the spheroid grows necrotic matter forms at the centre of the spheroid and appears to merge with the necrotic matter located closer to the periphery (3, 4.5 and 6 days after *t*_s_ in Figure 5m). As the WM983b spheroids do not maintain spherical symmetry we do not interpret these experimental data with the re-oxygenation mathematical model. Schematics describing the behaviours are presented in Figure 5l,n.

Confocal images for WM164 spheroids suggest that symmetry is maintained (Figures S16, S17), and that WM164 spheroids transition transiently through the growth phases in reverse order before subsequently growing in the usual order. In contrast to WM793b spheroids, WM164 spheroids can resume growth while maintaining a necrotic core (Figure S29). These different behaviours appear to be dependent on *t*_s_, *R*_o_(*t*_s_) and *ξ*_n_(*t*_s_). To interpret WM164 re-oxygenation experiments we use the re-oxygenation model (Supplementary Discussion F). However, additional care should be exercised interpreting WM164 re-oxygenation results. Brightfield timelapse images show that mass from the necrotic core can move to the periphery and exit the spheroid (Figure S18, Movie S3).

## 3 Discussion

Tumours grow in complicated fluctuating external environments. However, spheroid experiments used to study tumours are typically performed in constant external environments and with oxygen partial pressures that are much greater than *in vivo*. To explore this gap between *in vitro* and *in vivo* conditions, we analyse a series of tumour spheroid experiments using mathematical modelling and statistical uncertainty quantification. Growing spheroids in time-dependent external oxygen conditions reveals a range of behaviours not observed in standard experimental protocols. For fifty years, tumour spheroid growth has been characterised by three sequential growth phases: phase (i) exponential growth; phase (ii) reduced exponential growth; and phase (iii) saturation. However, here in re-oxygenation experiments, spheroids can transiently undergo these growth phases in reverse. Furthermore, spheroids can lose their spherically symmetric structure and necrotic core. Deoxygenation experiments also show that large changes to the internal structure of spheroids can occur while the overall size remains constant. Overall, our results suggest that oxygen and internal structure play pivotal roles in spheroid growth and should be taken into account when interpreting spheroid experiments. This is important as many studies do not provide sufficient information to replicate oxygen conditions and do not measure spheroid internal structures.

Tumour spheroid growth is a complex process involving multiple mechanisms. However, the contribution of each mechanism to growth is unclear using experimentation alone. To quantitatively explore which mechanisms contribute to spheroid growth we use the seminal Greenspan mathematical model [10]. The model describes the growth of spheroids in normoxia and hypoxia remarkably well. Moreover, our analysis suggests that growth and formation of the necrotic core is reasonably described by oxygen diffusion and consumption, whereas the growth and formation of the inhibited region is more accurately described by waste production and diffusion. Using statistical uncertainty quantification we show that the rates at which different biological processes occur differ between normoxia and hypoxia. Therefore, external environmental conditions should be taken into account when interpreting tumour spheroid experiments. Previous studies analysing previously available experimental data with Greenspan’s model [15, 16] have not been able to distinguish between the two mechanisms. Further, our results build on studies analysing spheroid snapshots only [17, 35]. As the model captures the key dynamics, we do not include other biological mechanisms that may be relevant in future studies, for example glucose [4]. Introducing additional mechanisms prior to developing the deoxygenation and re-oxygenation models would complicate the analysis, and likely result in parameters being non-identifiable and not physically interpretable. Both of which we aim to avoid. Many mathematical models have been developed with additional mechanisms, but they have not been quantitatively tested with experimental data [25,31–34]. The experimental data and framework that we provide here are suitable to test such models.

Deoxygenation and re-oxygenation experiments reveal how spheroid overall size and internal structure adapt to changes in the external environment. However, without a mathematical modelling and statistical uncertainty quantification framework like we use here, the mechanisms underlying adaptation are challenging to identify and interpret. Extending Greenspan’s mathematical model allows us to interpret, analyse, and describe these deoxygenation and re-oxygenation experiments remarkably well. Parameter estimation and identifiability analysis, using profile likelihood analysis and Bayesian inference, allows us to identify and quantify the contributions of key biological mechanisms to adaptation and growth. For both models, the analysis identifies a narrow range of behaviours that differ in terms of the rates of adaptation in *t*_s_ < *t* < *t*_s_ + 1 [days] and long term dynamics (Supplementary Discussion D.2.1). These modelling predictions raise interesting questions (Supplementary Discussion D.2.1). In comparison to standard experimental protocols, we collect more data per time point and over a longer duration. Further, we build on previous studies [13,15,16] to improve on standard experimental designs by measuring the internal structure of spheroids and hypoxic regions in addition to the overall size. Even these improvements to standard protocols are insufficient to identify all adaptation processes. As with all studies, additional data would be beneficial. In particular, frequent measurements of oxygen and internal structure at early times would be useful but are challenging to obtain experimentally (Supplementary Discussion D.2).

Re-oxygenation experiments reveal unexpected necrotic core removal in WM983b spheroids. To the best of our knowledge this behaviour has not been previously described. The exact mechanisms underlying this behaviour are unclear. We hypothesise that small asymmetry at re-oxygenation, possibly in the distribution of proliferating cells, in combination with changes to cell-cell adhesion and physical interactions contribute. We also observe that mass from the necrotic core of WM164 spheroids can move to the periphery and exit the spheroid. It is unclear whether these observations are relevant to other cell lines. Interesting future work is to explore these unusual behaviours in greater detail. For example, can this phenomenon be induced by other external environmental changes, drug treatments, and *in vivo*.

This work lays the foundation for further studies bridging the gap between clinical conditions and standard experimental protocols. Here, we consider normoxia and hypoxia and switches between normoxia and hypoxia. Other oxygen conditions are also worth consideration, for example to mimic *in vivo* oxygen gradients [39], *in vivo* vascularisation, disrupted oxygen supplies, or cyclic hypoxia [1]. Microfluidic devices may be one useful approach [25], but challenges visualising the internal structure of spheroids throughout such experiments must be overcome. Intracellular responses to oxygen changes are also of interest [40,41]. While we focus here on the impact of changing the oxygen conditions, our framework is well suited to explore the role of other changing external conditions on spheroid growth, for example nutrient availability and mechanical confinement [4, 11]. Further, the framework can be extended to explore different treatment strategies, for example radiotherapy and chemotherapy [31, 42].

## 4 Methods

### 4.1 Mathematical modelling

#### 4.1.1 Greenspan’s mathematical model

Key elements of Greenspan’s mathematical model for fixed *p*_∞_ are included in the main text. Further details of the model derivation are included in Supplementary Discussion C.1. Recall that this model has two interpretations, that we refer to as hypotheses 1 and 2.

- Hypothesis 1 assumes that the necrotic and inhibited regions are both driven by oxygen diffusion and consumption.
- Hypothesis 2 assumes that the necrotic region is driven by oxygen diffusion and consumption whereas the inhibited region is driven by waste production and diffusion.

Here, we present the governing differential-algebraic system of equations for the outer radius, *R*_o_(*t*), necrotic radius, *R*_n_(*t*), and inhibited radius, *R*_i_(*t*), for both hypotheses 1 and 2,

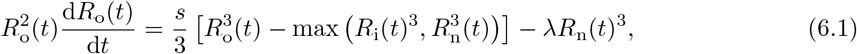

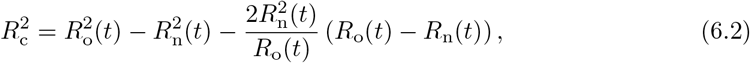

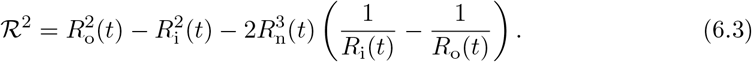

In these experiments, Equation (6.1) simplifies as *R*_i_(*t*) ≥ *R*_n_(*t*), consistent with our parameter choices (Methods 4.2) [16]. For both hypothesis 1 and 2: the outer radius when the necrotic region forms is *R*_c_ = [6kp_∞_/(*α*Ω)]^1/2^; Equation (6.1) arises from conservation of volume; and, Equation (6.2) is obtained by evaluating the oxygen partial pressure within the spheroid at the necrotic threshold. For hypothesis 1, Equation (6.3) is obtained by evaluating the oxygen partial pressure within the spheroid at the oxygen inhibited threshold, *p_i_*, and *Q*^2^ = (*p*_∞_ – *p*_i_)/*p*_∞_ so the left hand side of Equation (6.3) is 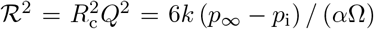. In contrast, for hypothesis 2, Equation (6.3) arises by evaluating the waste concentration within the spheroid at the waste inhibited threshold, *β_i_*, and *Q*^2^ = *β*_i_*κα*Ω/(*Pkp*_∞_) so the left hand side of Equation (6.3) is 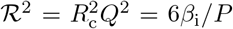. We solve the system of Equations (6.1)-(6.3) numerically (Supplementary Material C.1.2).

Analysing the model, the inhibited region forms at [10]

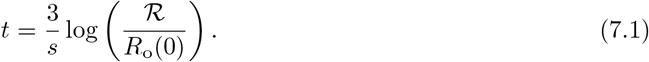

For hypothesis 1, Equation (7.1) is *t* = (3/*s*) log ([6*k*(*p*_∞_ – *p*_i_)/*α*]^1/2^/*R*_o_(0)). For hypothesis, 2 Equation (7.1) is *t* = (3/s)log ([6*β*_i_*κ/P*]^1/2^/*R*_o_(0)).

#### 4.1.2 Mathematical model to interpret deoxygenation experiments

Key elements of the mathematical model derived to interpret deoxygenation experiments are included in the main text. Further details of the model derivation are included in Supplementary Discussion C.2. Here, we present the governing equations for 0 < *t* < *t*_s_ and *t* > *t*_s_.

For 0 < *t* < *t*_s_ we solve Greenspan’s mathematical model [10] in normoxia interpreting the governing mechanisms with hypothesis 2. The differential-algebraic system of Equations (6.1)-(6.3) are solved to determine *R*_o_(*t*), *R*_n_(*t*), and *R*_i_(*t*).

After deoxygenation, *t* > *t*_s_, we extend Greenspan’s mathematical model to account for adaptation to hypoxia. Rewriting Equations (4), (5), (6.1), and solving Equation (6.2) for 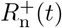 instead of *R*_n_(*t*), gives the governing differential-algebraic system of equations

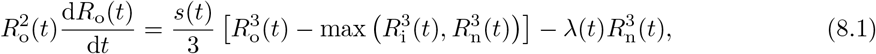

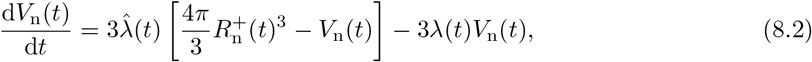

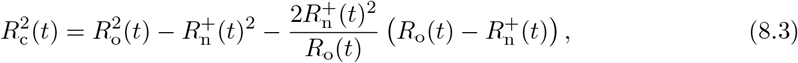

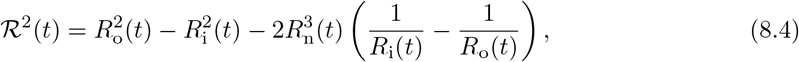

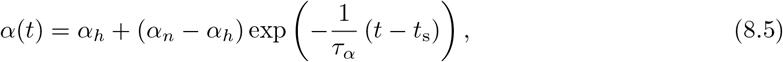

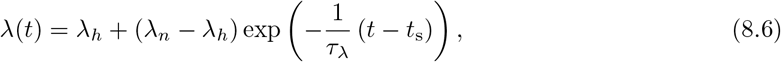

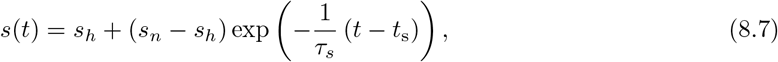

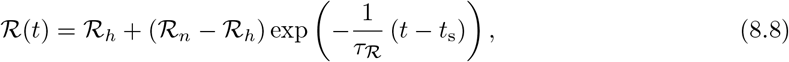

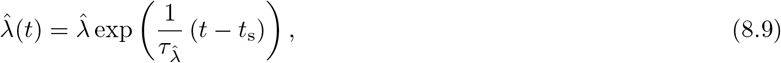

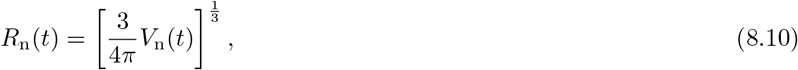

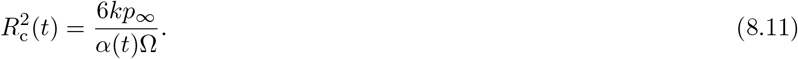

Note that in the long time limit *t* → ∞, we recover Greenspan’s mathematical model for normoxia (Equations (6.1)-(6.3)). Specifically, *α*(*t*) → *α_h_*, *λ*(*t*) → *λ*_*h*_, *s*(*t*) → *s_h_* and 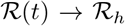 as *t* → ∞. Further, the term involving 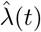 dominates the right hand side of Equation (8.2) as *t* → ∞, so 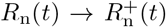 as *t* → ∞. We solve the system of equations (8.1)-(8.11) numerically numerically (Supplementary Material C.2.2).

#### 4.1.3 Mathematical model to interpret re-oxygenation experiments

Key details of the mathematical model to interpret re-oxygenation experiments are included in Supplementary Discussion C.3.

### 4.2 Parameter estimation and identifiability analysis

Parameter estimation and identifiability analysis are performed using profile likelihood analysis [15, 16, 43, 44], Bayesian inference [45–47], and global optimisation techniques, as now detailed. Throughout we exclude outliers in the experimental data. To detect outliers for each experiment we analyse each measurement type, *R*_o_(*t*), *R*_n_(*t*), and *R*_i_(*t*) at each time point independently using MATLABs `isoutlier` function with method *quartiles.*

#### 4.2.1 Greenspan’s model

Parameter estimation and identifiability analysis for Greenspan’s model are first performed using profile likelihood analysis (Figure 3), see [16]. The Bayesian inference approach to estimate parameters of Greenspan’s model is discussed in the following.

#### 4.2.2 Deoxygenation experiments

Parameter estimation for the deoxygenation model (Equations (8.1)-(8.11)) is performed using global optimisation and Bayesian inference techniques. We now explain how we estimate the fifteen model parameters, 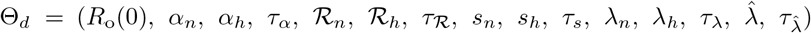. Informed by experimental measurements we set *R*_n_(*t*_s_) = 0 and *R*_i_(*t*_s_) = 0, but they could be included as additional parameters in future work.

We revisit Greenspan’s mathematical model for spheroids grown in normoxia and hypoxia for first estimates of *s_n_, s_h_, λ_n_* and *λ_h_*. Starting with experimental data from spheroids grown in normoxia, we fit a normal distribution to the initial outer radius measurements using the MATLAB `fitdist` function. Then for each spheroid we estimate *R*_c_ using Equation (6.2) given measurements of *R*_o_(*t*) and *R*_n_(*t*) and fit a normal distribution using the MATLAB `fitdist` function. Similarly, we estimate 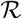 using Equation (6.3) and measurements of *R*_o_(*t*), *R*_n_(*t*) and *R*_i_(*t*) and fit a normal distribution using the MATLAB `fitdist` function. Next, we seek to estimate the five parameters of Greenspan’s model, 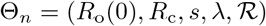, using the MATLAB package *MCMCstat* developed by Marko Laine [48,49]. Detailed information on the *MCMCstat* package is available on the GitHub repository (https://mjlaine.github.io/mcmcstat/).

Before we can use the *MCMCstat* package we require good first estimates of Θ_*n*_ and the mean squared error, *mse.* To provide a good first estimate for Θ_*n*_, we perform global optimisation using the MATLAB `GlobalSearch` function with settings: `fmincon` *sqp* algorithm; *MaxTime* = 60 [minutes]; *NumTrialPoints* = 5000; and *lowerbounds* and *upperbounds* informed by fitted normal distributions for *R*_o_(0), *R*_c_, and 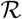 and previous results [16]. To estimate the *mse* we use experimental measurements as observations and simulate the deoxygenation model with the estimate of Θ_*n*_ from global optimisation to obtain predicted values. Next we use the *MCMCstat* package to generate four MCMC chains with 250,000 samples and enable automatic sampling and estimation of the error standard deviation. Next we discard the first 50,000 samples of each of the four chains as burn-in, resulting in each chain containing 200,000 samples. For other *MCMCstat* package options we use the default settings. To assess convergence of the MCMC chains we compute the potential scale reduction factor, 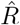, [47] for each parameter and observe convergence, where convergence corresponds to 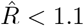 (Table S2). Performing posterior checks, using 50,000 samples from the chains to generate prediction intervals and comparing with the experimental data, suggests the parameter estimates are reasonable. This process is repeated for the hypoxia data to estimate Θ*h* (Table S3). The posteriors generated here are for *s_n_, s_h_, λ_n_* and *λ_h_*.

Next, we analyse the deoxygenation experimental data at each spheroid and time point independently. To estimate 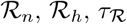 from Equation (8.8) we use global minimisation and the *MCMCstat* package. For estimates of 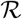 for each spheroid and at each time point independently we use Equation (6.2) and measurements of *R*_o_(*t*) and *R*_n_(*t*). Similarly, to estimate *α_n_, α_h_, τ_α_* from Equation (8.5) we use global minimisation and the *MCMCstat* package. For estimates of *α* we use Equations (8.11) and (6.3) and measurements of *R*_o_(*t*), *R*_n_(*t*) and *R*_i_(*t*). Note that to estimate *R*_c_ we assume that 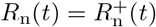. To estimate *R*_o_(0) we fit a normal distribution using the MATLAB `fitdist` function to measurements of *R*_o_(*t*) at the first time point.

Next we perform a global minimisation to estimate Θ_*d*_, using the estimates of 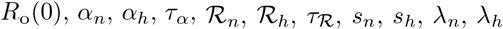 to inform *lowerbounds* and *upperbounds.* To estimate Θ_*d*_ we use global minimisation and the *MCMCstat* package (Table S4). Performing posterior checks, using 50,000 samples from the chain to generate prediction intervals and comparing with the experimental data, suggests the parameter estimates are reasonable (Figures S26, S28, S29).

#### 4.2.3 Re-oxygenation experiments

Parameter estimation for the re-oxygenation model formed by Equations (S.36.1)-(S.36.10), with parameters 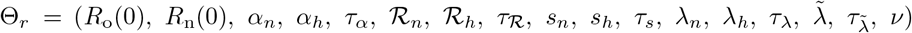, is analogous to the approach used for the deoxygenation model. Summary statistics for the MCMC chains and MCMC diagnostics are shown in Tables S5 and S6. Comparing prediction intervals and with the experimental data suggests the parameter estimates are reasonable (Figures S28, S29).

### 4.3 Experimental methods

#### Cell culture

The human melanoma cell lines established from primary (WM793b) and metastatic cancer sites (WM983b, WM164) were provided by Prof. Meenhard Herlyn, The Wistar Institute, Philadelphia, PA, [50]. All cell lines were previously transduced with fluorescent ubiquitination-based cell cycle indicator (FUCCI) constructs [13, 14]. Cell lines were previously genotypically characterised [13,51–53], and authenticated by short tandem repeat fingerprinting (QIMR Berghofer Medical Research Institute, Herston, Australia). The cells were cultured in melanoma cell medium (“Tu4% medium”): 80% MCDB-153 medium (Sigma-Aldrich, M7403), 20% L-15 medium (Sigma-Aldrich, L1518), 4% fetal bovine serum (ThermoFisher Scientific, 25080-094), 5mg mL^-1^ insulin (Sigma-Aldrich, I0516), 1.68 mM CaCl_2_ (Sigma-Aldrich, 5670) [14]. Cell lines were checked routinely for mycoplasma and tested negative using the MycoAlert MycoPlasma Detection Kit (Lonza) and polymerase chain reaction [54].

#### Spheroid generation, culture, and experiments

Spheroids were generated in 96-well cell culture flat-bottomed plates (3599, Corning), with 5000 total cells/well, using 50 μL total/well non-adherent 1.5% agarose to promote formation of a single spheroid per well [30]. From previous work we expect that different seeding densities, in the range 1250-10000 total cells/well, will provide similar results [15,16]. For all experiments spheroids formed after 2 days for WM793b, WM164 and WM983b. On day 3 and 7 of each experiment 50% of the medium in each well was replaced with fresh medium (200μL total/well). Each experiment was performed for 8 days, informed by previous experiments with these cell lines so that necrotic and inhibited region form prior to the end of the experiments [16].

For normoxia experiments, cells and spheroids were grown and formed in an incubator with standard settings: 37°C, 5% CO_2_ [13,14]; referred to as the normoxia incubator. For hypoxia experiments, cells were cultured in the normoxia incubator and then spheroids were grown in a hypoxia incubator with settings: 37 °C, 5% CO_2_, 2% O_2_. For deoxygenation experiments, cells and spheroids were formed and grown in the normoxia incubator. At the time of deoxygenation, the relevant plate(s) of spheroids were manually transferred from the normoxia incubator to the hypoxia incubator. Similarly, for the re-oxygenation experiments spheroids were grown in the hypoxia incubator then at the time of re-oxygenation the relevant plate(s) of spheroids were manually transferred to the normoxia incubator. The time to move plates was 1-2 minutes.

To estimate the outer, necrotic, inhibited, and hypoxic radii, we use a high-throughput method of mounting, clearing and imaging [55]. Spheroids maintained in the relevant incubator were harvested, fixed with 4% paraformaldehyde (PFA), and stored in phosphate buffered saline solution (PBS), sodium azide (0.02%), Tween-20 (0.1%), and DAPI (1:2500) at 4 °C, on days 2, 3, 4, 6 and 8 after seeding. For hypoxia measurements, spheroids were stained with 100mM pimonidazole for three hours, prior to fixation. Spheroids were then permeabilized with 0.5% Triton X-100 in PBS for one hour; blocked in antibody dilution buffer (Abdil) [56] for 24 hours; stained with a 1:50 anti-pimonidazole mouse IgG1 monoclonal antibody (Hypoxyprobe-1 MAb1) in Abdil for 48 hours; washed in PBS with 0.1% Tween-20 for 6 hours; placed in a 1:100 solution of goat anti-mouse Alexa Flour 647 in Abdil for 48 hours; and, finally washed for 6 hours in PBS. Then for imaging, fixed spheroids were set in place using low melting 2% agarose and optically cleared in 500 μL total/well high refractive index mounting solution (Quadrol 9 % wt/wt, Urea 22 % wt/wt, Sucrose 44 % wt/wt, Triton X-100 0.1 % wt/wt, water) for 2 days in a 24-well glass bottom plate (Cellvis, P24-1.5H-N) before imaging to ensure accurate measurements [55,57,58]. Images were then captured using an Olympus FV3000 confocal microscope with the 10 × objective focused on the equatorial plane of each spheroid.

As the unexpected necrotic core movement for WM983b cell line was observed in the re-oxygenation experiments, the re-oxygenation experiments were repeated for all cell lines alongside a control normoxia condition. Spheroids were cultured into three 96-well plates (3599, Corning): plate (i) control for normoxia; plate (ii) re-oxygenation 2.5 days after seeding; and, plate (iii) re-oxygenation 5.5 days after seeding. Each plate consisted of 32 spheroids of each cell line. The plates were placed inside the IncuCyte S3 live cell imaging system (Sartorius, Goettingen, Germany) incubator (37°C, 5% CO_2_). IncuCyte S3 settings were chosen to image with the 4× objective. For plate (i) images were captured every 2 hours for the first three days and then every 4 hours for the remainder of the experiment. For plate (ii) and (iii) images were captured every hour for three and seven days, respectively.

#### Image processing

Confocal microscopy images were converted to TIFF files in ImageJ and then processed with custom MATLAB scripts that use standard MATLAB image processing toolbox functions. Area was converted to an equivalent radius (*r*^2^ = *A/π*). These scripts are freely available on Zenodo with DOI:10.5281/zenodo.5121093 [59], with modifications to account for pimonidazole staining and blurred central regions due to hypoxia and deoxygenation discussed in Supplementary Discussion B. Images captured with the IncuCyte S3 were processed with custom MATLAB scripts that use standard MATLAB image processing toolbox functions and are detailed in Supplementary Discussion D.3.1.

#### Statistics and Reproducibility

Details of practical parameter identifiability analysis and the Bayesian inference are presented in Section 4.2. Each radial measurements is represented as an individual data point in relevant figures, with non-filled circles representing outliers (Section 4.2), and are summarised using box charts. Supplementary Table S1 details the number of measurements at each time point for each cell line and experimental data analysed during the study are available on a GitHub repository (https://github.com/ryanmurphy42/Murphy2022SpheroidOxygenAdaptation). We note that some measurements could not be obtained primarily due to blurring of the automated imaging, spheroids not forming properly, or spheroids losing their structural integrity at late times. Data for these spheroids was excluded. In a previous study we assess experimental designs [16] and use this to inform that our sample size is sufficient in this study. Randomisation and blinding was not possible.

## Data Availability

The datasets generated during and analysed during the current study are available on a GitHub repository (https://github.com/ryanmurphy42/Murphy2022SpheroidOxygenAdaptation) and are summarised in the electronic supplementary material.

## Code Availability

Key computer code and all experimental data used to generate computational results are available on a GitHub repository (https://github.com/ryanmurphy42/Murphy2022SpheroidOxygenAdaptation) repository. The computer code for the mathematical modelling and statistical identifiability analysis was written in MATLAB R2021b (v9.11) with the Image Processing Toolbox (v11.4), Optimization Toolbox (v9.2), Global Optimization Toolbox (v4.6), and the Statistics and Machine Learning Toolbox (v12.2), and uses the *MCMCstat* package available on the GitHub repository (https://mjlaine.github.io/mcmcstat/).

## Author Contributions

All authors conceived and designed the study. R.J.M. performed the research and drafted the article. G.G. and R.J.M. performed experimental work. All authors provided comments and approved the final version of the manuscript. N.K.H. and M.J.S. contributed equally.

## Acknowledgements

We thank Dr Alexander P. Browning and Dr Patrick B. Thomas for helpful discussions, and John Blake for guidance using IncuCyte. This research was carried out at the Translational Research Institute (TRI), Woolloongabba, QLD. TRI is supported by a grant from the Australian Government. We thank the staff in the microscopy core facility at TRI for their technical support. We thank Prof. Atsushi Miyawaki, RIKEN, Wako-city, Japan, for providing the FUCCI constructs, Prof. Meenhard Herlyn, The Wistar Institute, Philadelphia, PA, for providing the cell lines.

## References

[1] Bader, S. B., Dewhirst, M. W. & Hammond, E. M. Cyclic hypoxia: An update on its characteristics, methods to measure it and biological implications in cancer. Cancers 13, 23 (2021).

[2] Kumar, B. et al. Tumor collection/processing under physioxia uncovers highly relevant signaling networks and drug sensitivity. Science Advances 8, eabh3375 (2022).

[3] Vaupel, P., Kallinowski, F. & Okunieff, P. Blood flow, oxygen and nutrient supply and metabolic enviroment of human tumours: A review. Cancer Research 49, 6449–6465 (1989).

[4] Mueller-Klieser, W., Freyer, J. & Sutherland, R. Influence of glucose and oxygen supply conditions on the oxygenation of multicellular spheroids. British Journal of Cancer 53, 345–353 (1986).

[5] Hirschhaeuser, F. et al. Multicellular tumor spheroids: An underestimated tool is catching up again. Journal of Biotechnology 148, 3–15 (2010).

[6] Costa, E. C. et al. 3D tumor spheroids: an overview on the tools and techniques used for their analysis. Biotechnology Advances 34, 1427–1441 (2016).

[7] Nath, S. & Devi, G. R. Three-dimensional culture systems in cancer research: Focus on tumor spheroid model. Pharmacology & Therapeutics 163, 94–108 (2016).

[8] Mehta, G., Hsiao, A. Y., Ingram, M., Luker, G. D. & Takayama, S. Opportunities and challenges for use of tumor spheroids as models to test drug delivery and efficacy. Journal of Controlled Release 10, 192–204 (2012).

[9] Weiswald, L., Bellet, D. & Dangles-Marie, V. Spherical cancer models in tumor biology. Neoplasia 17, 1–15 (2015).

[10] Greenspan, H. P. Models for the growth of a solid tumor by diffusion. Studies in Applied Mathematics 51, 317–340 (1972).

[11] Le Maout, V. et al. Role of mechanical cues and hypoxia on the growth of tumor cells in strong and weak confinement a dual in vitro in silico approach. Science Advances 6, eaaz7130 (2020).

[12] Beaumont, K. A., Mohana-Kumaran, N. & Haass, N. K. Modeling melanoma in vitro and in vivo. Healthcare 2, 27–46 (2014).

[13] Haass, N. K. et al. Real-time cell cycle imaging during melanoma growth, invasion, and drug response. Pigment Cell & Melanoma Research 27, 764–776 (2014).

[14] Spoerri, L., Beaumont, K. A., Anfosso, A. & Haass, N. K. Real-time cell cycle imaging in a 3D cell culture model of melanoma. Methods in Molecular Biology 1612, 401–416 (2017).

[15] Browning, A. P. et al. Quantitative analysis of tumour spheroid structure. eLife 10, e73020 (2021).

[16] Murphy, R. J., Browning, A. P., Gunasingh, G., Haass, N. K. & Simpson, M. J. Designing and interpreting 4D tumour spheroid experiments. Communications Biology 5, 91 (2022).

[17] Gomes, A. et al. Oxygen partial pressure is a rate-limiting parameter for cell proliferation in 3D spheroids grown in physioxic culture condition. PLoS One 11, e0161239 (2016).

[18] Grimes, D. R. et al. The role of oxygen in avascular tumor growth. PLoS One 11, e0153692 (2016).

[19] Muz, B., de la Puente, P., Azab, F. & Azab, A. B. The role of hypoxia in cancer progression, angiogenesis, metastasis, and resistance to therapy. Hypoxia 3, 83–92 (2015).

[20] McKeown, S. R. Defining normoxia, physoxia and hypoxia in tumours-implications for treatment response. British Journal of Radiology 87, 20130676 (2014).

[21] Carreau, A., El Hafny-Rahbi, B., Matejuk, A., Grillon, C. & Kieda, C. Why is the partial oxygen pressure of human tissues a crucial parameter? small molecules and hypoxia. Journal of Cellular and Molecular Medicine 15, 1239–1253 (2011).

[22] Celora, G. L. et al. A DNA-structured mathematical model of cell-cycle progression in cyclic hypoxia. Journal of Theoretical Biology 111104 (2022).

[23] Lee, P., Chandel, N. S. & Simon, M. C. Cellular adaptation to hypoxia through hypoxia inducible factors and beyond. Nature Reviews Molecular Cell Biology 21, 268–283 (2020).

[24] Goto, T., Kaida, A. & Miura, M. Visualizing cell-cycle kinetics after hypoxia/reoxygenation in heLa cells expressing fluorescent ubiquitination-based cell cycle indicator (Fucci). Experimental Cell Research 339, 389–396 (2015).

[25] Grist, S. M. et al. Long-term monitoring in a microfluidic system to study tumour spheroid response to chronic and cycling hypoxia. Scientific Reports 9, 17782 (2019).

[26] Fridman, I. B., Ugolini, G. S., VanDelinder, V., Cohen, S. & Konry, T. High throughput microfluidic system with multiple oxygen levels for the study of hypoxia in tumor spheroids. Biofabrication 13, 035037 (2021).

[27] Riffle, S. & Hegde, R. S. Modeling tumor cell adaptations to hypoxia in multicellular tumor spheroids. Journal of Experimental & Clinical Cancer Research 36, 102 (2017).

[28] Al-Ani, A. et al. Oxygenation in cell culture: Critical parameters for reproducibility are routinely not reported. PLoS One 13, e0204269 (2018).

[29] Sakaue-Sawano, A. et al. Visualizing spatiotemporal dynamics of multicellular cell-cycle progression. Cell 132, 487–498 (2008).

[30] Spoerri, L., Gunasingh, G. & Haass, N. K. Fluorescence-based quantitative and spatial analysis of tumour spheroids: a proposed tool to predict patient-specific therapy response. Frontiers in Digital Health 3, 668390 (2021).

[31] Bull, J. A. & Byrne, H. M. The hallmarks of mathematical oncology. Proceedings of the IEEE 110, 523–540 (2022).

[32] Araujo, R. P. & McElwain, D. L. S. A history of the study of solid tumour growth: the contribution of mathematical modelling. Bulletin of Mathematical Biology 66, 1039 (2004).

[33] Byrne, H. M. Dissecting cancer through mathematics: from the cell to the animal model. Nature Reviews Cancer 10, 221–230 (2010).

[34] Roose, T., Chapman, S. J. & Maini, P. K. Mathematical models of avascular tumor growth. SIAM Review 49, 179–208 (2007).

[35] Grimes, D. R., Kelly, C., Bloch, K. & Partridge, M. A method for estimating the oxygen consumption rate in multicellular tumour spheroids. Journal of the Royal Society Interface 11, 20131124 (2014).

[36] Santilli, G. et al. Mild hypoxia enhances proliferation and multipotency of human neural stem cells. PLoS One 5, e8575 (2010).

[37] Grayson, W. L., Zhao, F., Bunnell, B. & Ma, T. Hypoxia enhances proliferation and tissue formation of human mesenchymal stem cells. Biochemical and Biophysical Research Communications 358, 948–953 (2007).

[38] Danet, G. H., Pan, Y., Luongo, J. L., Bonnet, D. A. & Simon, M. C. Expansion of human SCID-repopulating cells under hypoxic conditions. The Journal of Clinical Investigation 112, 126–135 (2003).

[39] Leedale, J. et al. In silico-guided optimisation of oxygen gradients in hepatic spheroids. Computational Toxicology 12, 100093 (2019).

[40] Cavadas, M. A. S., K., N. L. & Cheong, A. Hypoxia-inducible factor (HIF) network insights from mathematical models. Cell Communication and Signaling 11, 42 (2013).

[41] Leedale, J. et al. Modeling the dynamics of hypoxia inducible factor-1α (HIF-1α) within single cells and 3D cell culture systems. Mathematical Biosciences 258, 33–43 (2014).

[42] Lewin, T. D., Maini, P. K., Moros, E. G., Enderling, H. & Byrne, H. M. The evolution of tumour composition during fractionated radiotherapy implications for outcome. Bulletin of Mathematical Biology 80, 1207–1235 (2018).

[43] Pawitan, Y. In All Likelihood: Statistical Modelling And Inference Using Likelihood (Oxford University Press, Oxford, UK, 2001).

[44] Simpson, M. J., Baker, R. E., Vittadello, S. T. & Maclaren, O. J. Practical parameter identifiability for spatio-temporal models of cell invasion. Journal of the Royal Society Interface 17, 2020055 (2020).

[45] Browning, A. P., Haridas, P. & Simpson, M. J. A Bayesian sequential learning framework to parameterise continuum models of melanoma invasion into human skin. Bulletin of Mathematical Biology 81, 676–698 (2019).

[46] Collis, J. et al. Bayesian calibration, validation and uncertainty quantification for predictive modelling of tumour growth: a tutorial. Bulletin of Mathematical Biology 79, 939–974 (2017).

[47] Gelman, A. et al. Bayesian Data Analysis (Chapman and Hall/CRC, New York, 2013), 3 edn.

[48] Haario, H., Saksman, E. & Tamminen, J. An adaptive Metropolis algorithm. Bernoulli 7, 223–242 (2001).

[49] Haario, H., Laine, M., Mira, A. & Saksman, E. DRAM: Efficient adaptive MCMC. Statistics and Computing 16, 339–354 (2006).

[50] Hsu, M. Y., Elder, D. E. & Herylyn, M. Melanoma: the Wistar melanoma (WM) cell lines. Human Cell Culture 1, 259–274 (2002).

[51] Hoek, K. S. et al. Metastatic potential of melanomas defined by specific gene expression profiles with no BRAF signature. Pigment Cell Research 19, 290–302 (2006).

[52] Smalley, K. S. M. et al. An organometallic protein kinase inhibitor pharmacologically activates p53 and induces apoptosis in human melanoma cells. Cancer Research 67, 209–217 (2007).

[53] Smalley, K. S. M. et al. Ki67 expression levels are a better marker of reduced melanoma growth following MEK inhibitor treatment than phospho-ERK levels. British Journal of Cancer 96, 445–449 (2007).

[54] Uphoff, C. C. & Drexler, H. G. Detecting mycoplasma contamination in cell cultures by polymerase chain reaction. Methods in Molecular Biology 731, 93–103 (2011).

[55] Gunasingh, G., Browning, A. P. & Haass, N. K. Rapid optical clearing for high-throughput analysis of tumour spheroids. Preprints (2021).

[56] Cold Spring Harbor Laboratory Press. Antibody dilution buffer (Abdil) (2018). Accessed: November 2021.

[57] Costa, E. C., Silva, D. B., Moreira, A. F. & Correia, I. J. Optical clearing methods: an overview of the techniques used for the imaging of 3D spheroids. Biotechnology & Bioengineering 116, 2742–2763 (2019).

[58] Susaki, E. A. et al. Versatile whole-organ/body staining and imaging based on electrolyte-gel properties of biological tissues. Nature Communications 11, 1982 (2020).

[59] Browning, A. P. & Murphy, R. J. Image processing algorithm to identify structure of tumour spheroids with cell cycle labelling. Zenodo (2021). https://doi.org/10.5281/zenodo.5121093.

