## Supplementary Material for "Growth and adaptation mechanisms of tumour spheroids with time-dependent oxygen availability"

#### Supplementary Discussion **Page No.**


A. Experimental data 3

A.1 Data summary 3

A.2 Experimental images 4

B. Image processing 24

C. Mathematical model additional details 26

C.1 Greenspan's mathematical model 26

C.1.1 Model derivation 26

C.1.2 Numerical methods 29

C.2 Mathematical model to interpret deoxygenation experiments 30

C.2.1 Model derivation 30

C.2.2 Numerical methods 31

C.3 Mathematical model to interpret re-oxygenation experiments 33

C.3.1 Model derivation 33

C.3.2 Numerical methods 36

D. Additional results for WM983b spheroids 38

D.1 Oxygen diffusion alone is insufficient to describe spheroid growth 38

D.1.1 Analysing spheroid snapshots independently to explore oxygen assumptions 39

D.1.2 Analysing spheroid snapshots independently to explore waste assumptions 40

D.1.3 Parameter estimation 41

D.2 Deoxygenation 42

D.2.1 Parameter estimation 43

D.3 Re-oxygenation 44

D.3.1 Necrotic core movement in WM983b spheroids 44

E. Additional results for WM793b cell line 45

F. Additional results for WM164 cell line 46

G. Additional results: Summary statistics and MCMC diagnostics 47

H. Supplementary Movie Descriptions 52

H.1 Movie S1 52

H.2 Movie S2 52

H.3 Movie S3 52

### A Experimental data

#### A.1 Data summary

Here we summarise the experimental data analysed in this study. In Table S1 we present the total number of spheroids measured for each experiment type and cell line. For each spheroid we use confocal microscopy and image processing to measure the outer radius, $R_{\mathrm{o}}(t)$, inhibited radius, $R_{\mathrm{i}}(t)$, necrotic radius, $R_{\mathrm{n}}(t)$ and hypoxic radius, $R_{\mathrm{p}}(t)$. Note that each spheroid is only measured once as we harvest, fix, and mount each spheroid before imaging. Day 0 corresponds to the start of the experiment when the spheroids were seeded.

| Experiment description | Day | WM983b | WM793b | WM164 |
|---|---|---|---|---|
| 1 - Normoxia | 2 | 15 | 6 | 9 |
| | 3 | 7 | 8 | 4 |
| | 4 | 15 | 13 | 9 |
| | 6 | 10 | 15 | 7 |
| | 8 | 10 | 10 | 13 |
| 2 - Hypoxia | 2 | 4 | 12 | 7 |
| | 4 | 7 | 11 | 8 |
| | 6 | 5 | 11 | 1 |
| | 8 | 12 | 7 | 5 |
| 3 - Deoxygenation on Day 2 | 3 | 12 | 12 | 11 |
| | 4 | 11 | 12 | 12 |
| | 6 | 9 | 10 | 8 |
| | 8 | 7 | 14 | 6 |
| 4 - Re-oxygenation on Day 2 | 3 | * | 15 | 11 |
| | 4 | * | 13 | 9 |
| | 6 | * | 13 | 13 |
| | 8 | * | 14 | 7 |
| 5 - Re-oxygenation on Day 4 | 6 | * | 11 | 14 |
| | 8 | * | 12 | 14 |

Table S1: Number of spheroids imaged with confocal microscopy for the WM983b, WM793b, and WM164 cell lines. For WM983b re-oxygenation experiments, denoted by *, we focus on brightfield images.

#### A.2 Experimental images

In Figures S1-S19 we present confocal microscopy and brightfield images of spheroids formed with the WM983b, WM793b, and WM164 human melanoma FUCCI transduced cell lines.

*Confocal images.* To clearly visualise the internal structure and hypoxic regions of spheroids we show each spheroid twice. In the top set of images we outline each spheroids outer boundary, inhibited region, and necrotic region obtained by analysing FUCCI fluorescence. In the bottom set of images we present the pimonidazole staining and outline the boundary which we convert to the hypoxic radius, $R_\mathrm{p}(t)$. Note that both sets of images are of the same spheroids. By using different channels in confocal microscopy we include or exclude the pimonidazole signal (far-red channel, shown as cyan) without interfering with the FUCCI signals (green and red channels, shown as green and magenta respectively). The boundary of each detected region is manually reviewed post-image processing. On occasion, the necrotic and hypoxic regions are not accurately identified and so we use ImageJ to measure the respective regions (Supplementary Discussion B).

*Brightfield images.* Re-oxygenation experiments with $t_\mathrm{s} = 2.5$ [days] and $t_\mathrm{s} = 5.5$ [days] are shown with brightfield images.

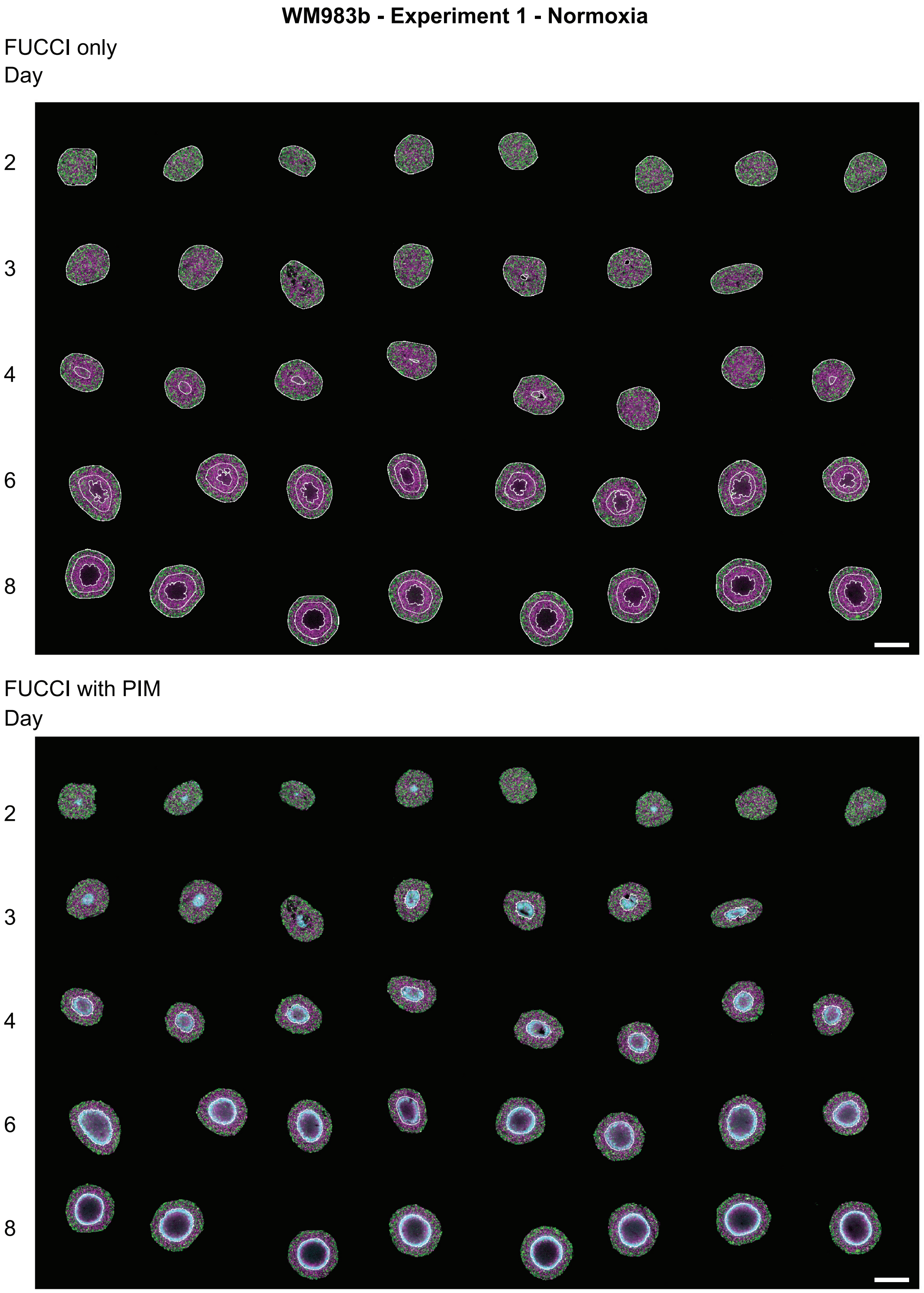


Figure S1: Experimental images of WM983b tumour spheroids in Experiment 1 - Normoxia. Top set of images shows spheroids with FUCCI signal only. Bottom set of images show spheroids with FUCCI signal and pimonidazole staining. Scale bars are 400µm.

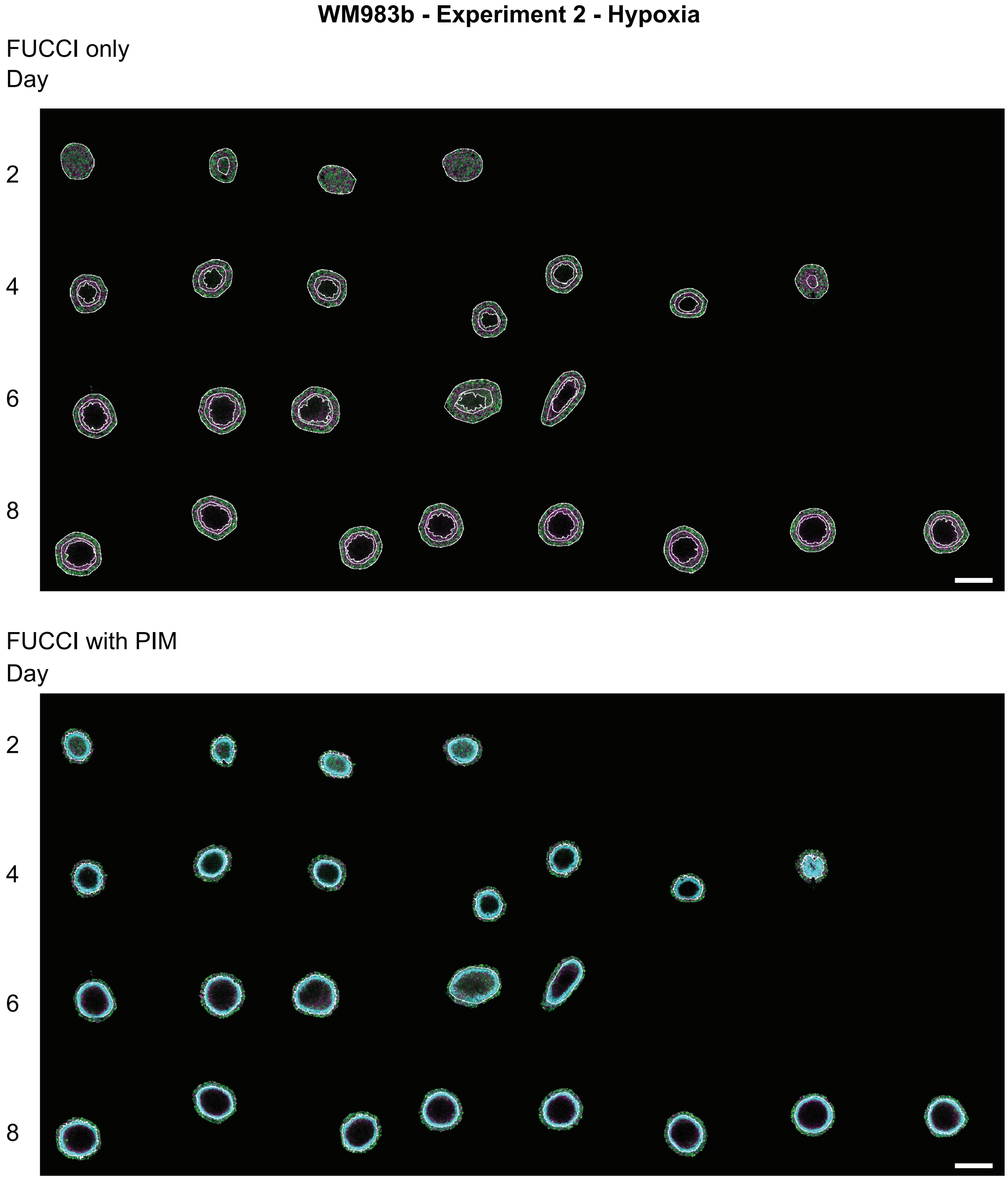


Figure S2: Experimental images of WM983b tumour spheroids in Experiment 2 - hypoxia. Top set of images shows spheroids with FUCCI signal only. Bottom set of images show spheroids with FUCCI signal and pimonidazole staining. Scale bars are 400µm.

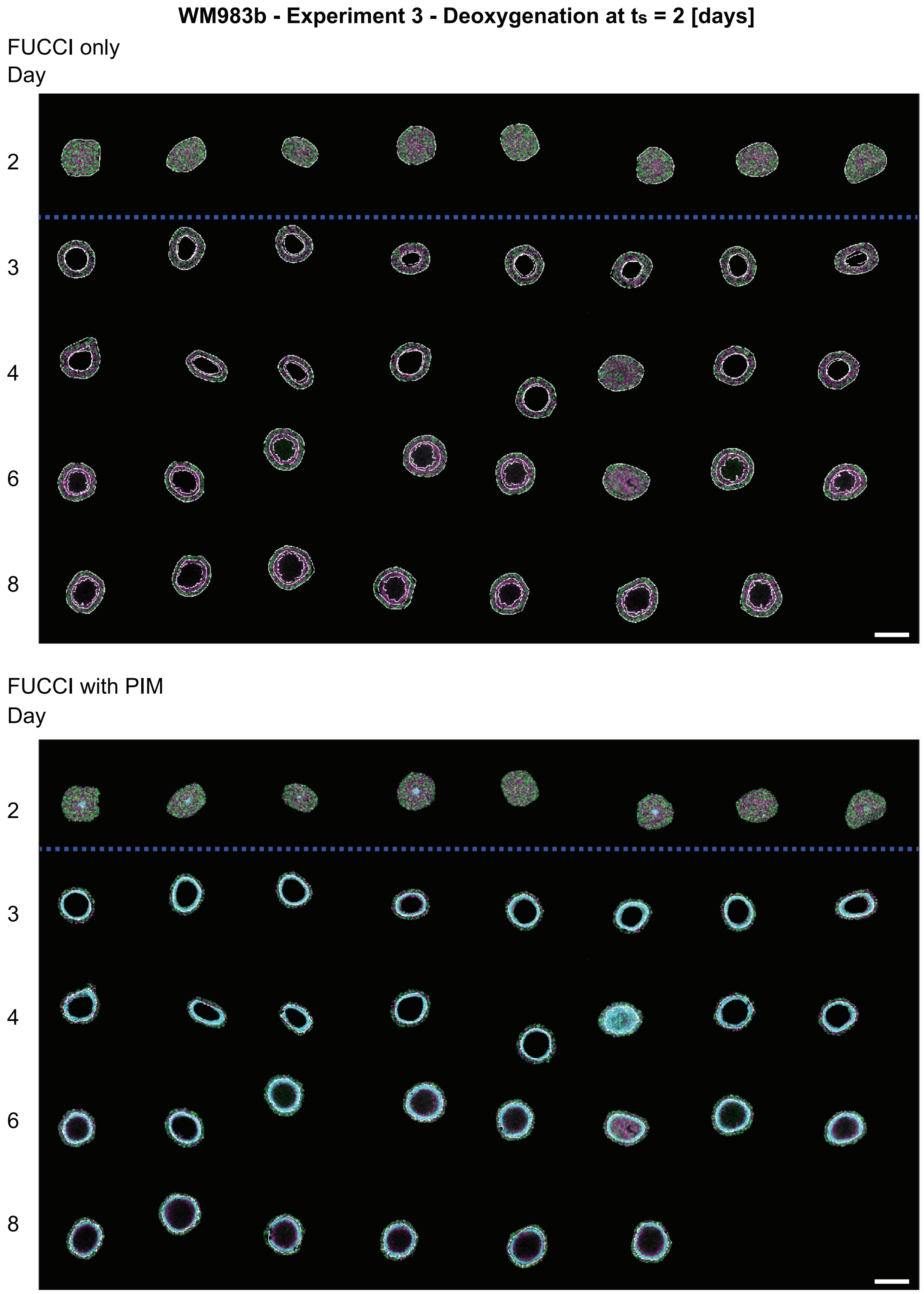


Figure S3: Experimental images of WM983b tumour spheroids in Experiment 3 - deoxygenation at $t_\mathrm{s} = 2$ [days] (blue dashed line).Top set of images shows spheroids with FUCCI signal only. Bottom set of images show spheroids with FUCCI signal and pimonidazole staining. Scale bars are 400µm.

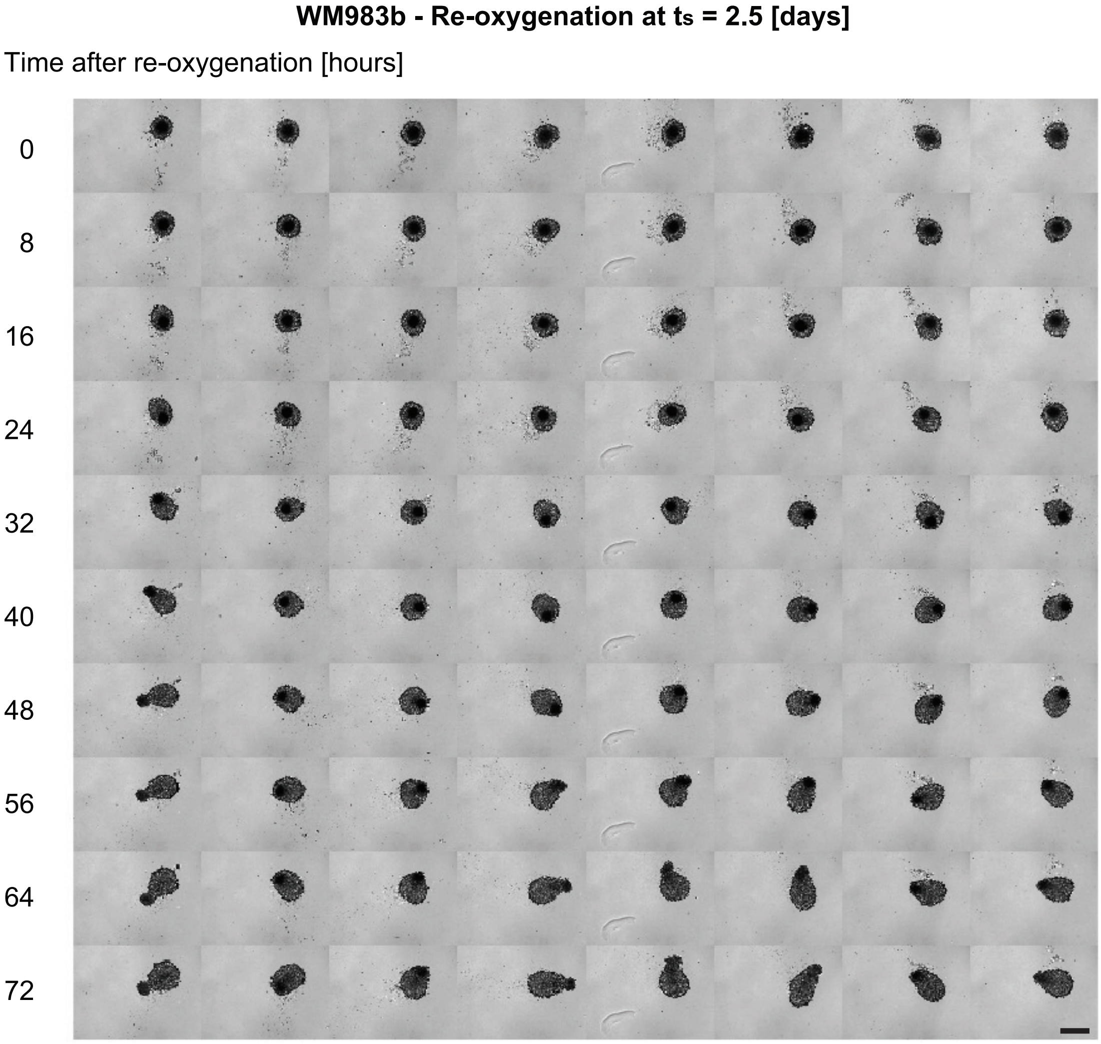


Figure S4: Experimental images of WM983b tumour spheroids in Experiment 6 - Re-oxygenation at $t_s = 2.5$ [days] (green dashed line). Scale bars are 400μm.

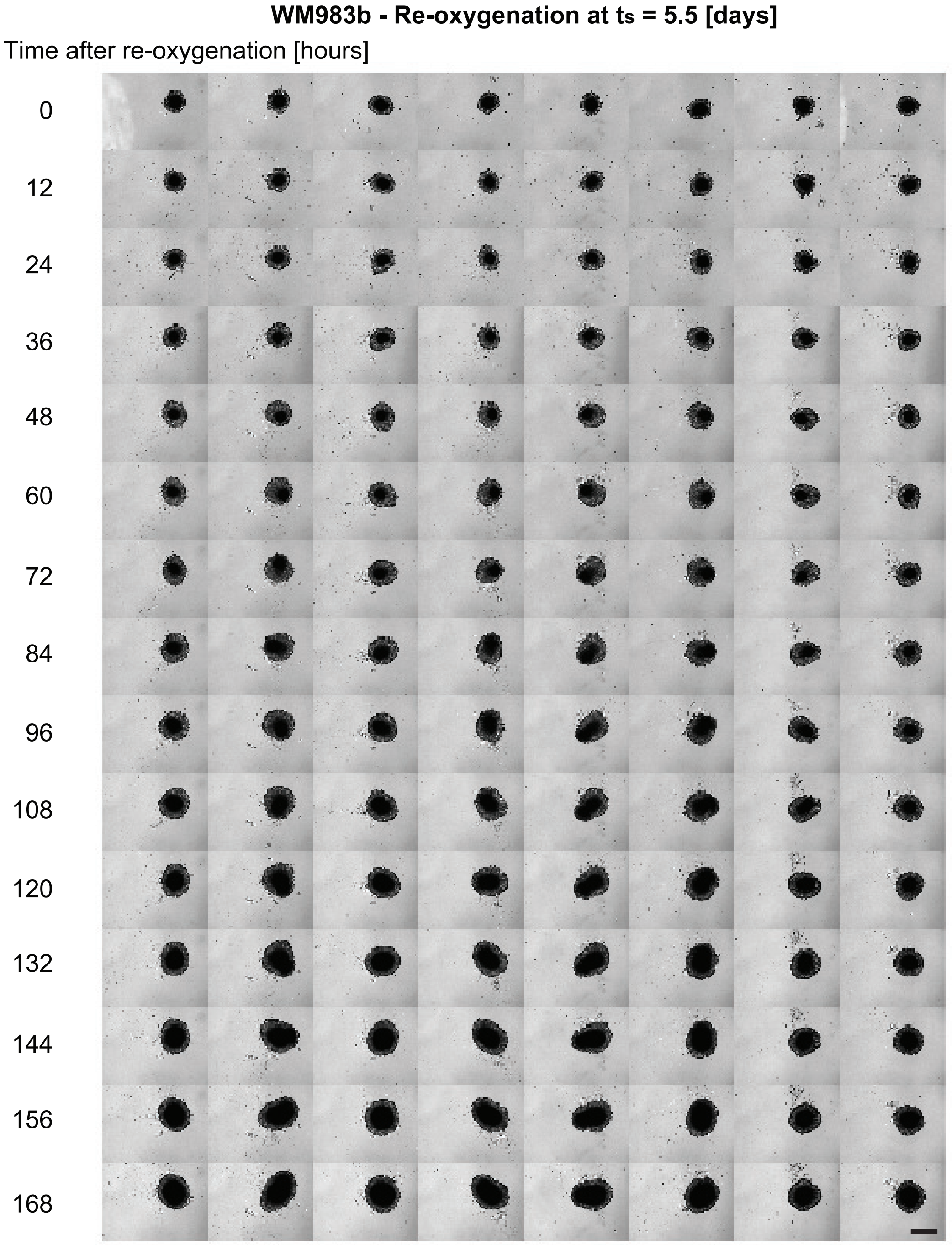


Figure S5: Experimental images of WM983b tumour spheroids in Experiment 7 - Re-oxygenation at $t_s = 5.5$ [days] (green dashed line). Scale bars are 400µm.

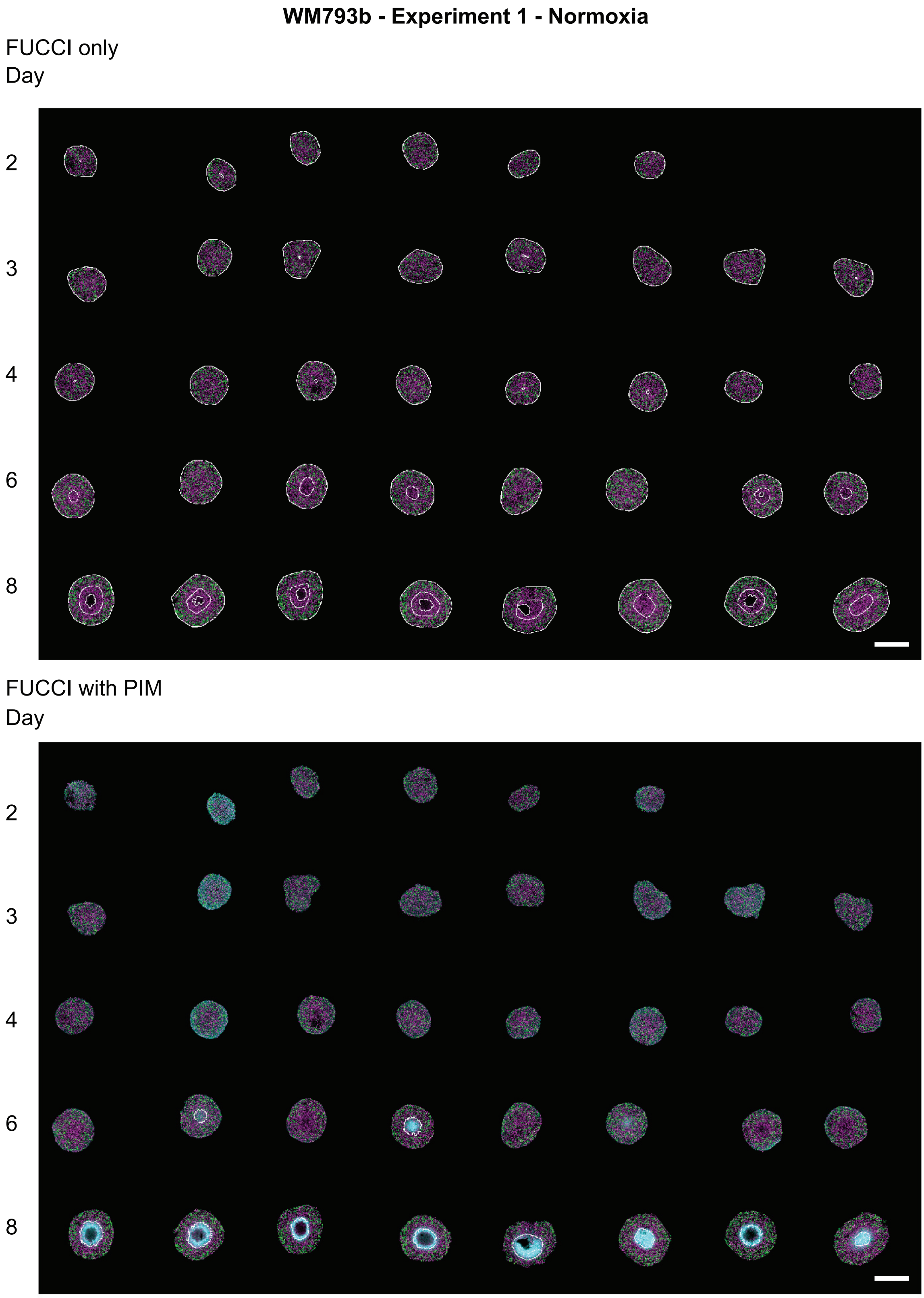


Figure S6: Experimental images of WM793b tumour spheroids in Experiment 1 - Normoxia.Top set of images shows spheroids with FUCCI signal only. Bottom set of images show spheroids with FUCCI signal and pimonidazole staining. Scale bars are 400µm.

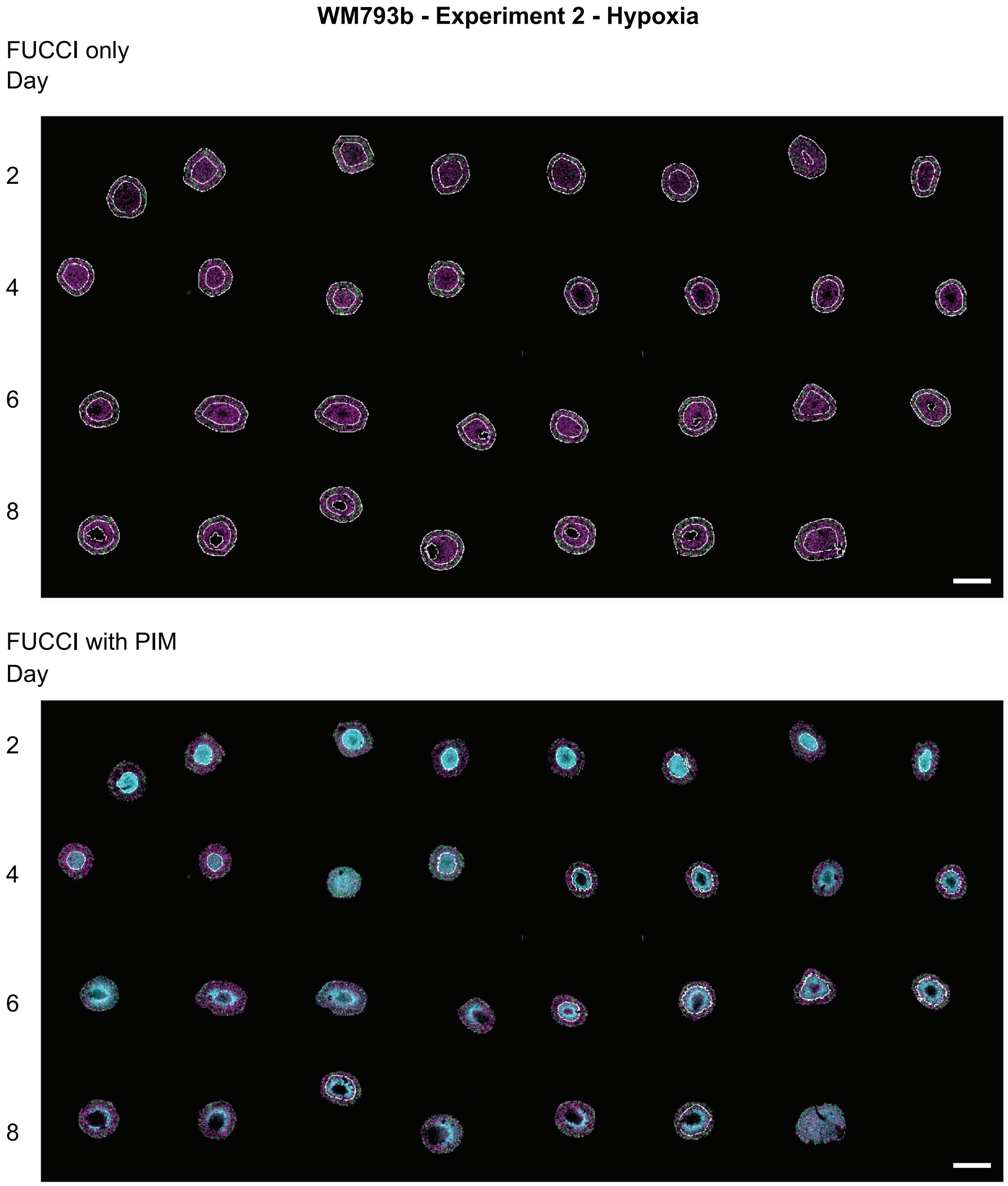


Figure S7: Experimental images of WM793b tumour spheroids in Experiment 2 - hypoxia. Top set of images shows spheroids with FUCCI signal only. Bottom set of images show spheroids with FUCCI signal and pimonidazole staining. Scale bars are 400µm.

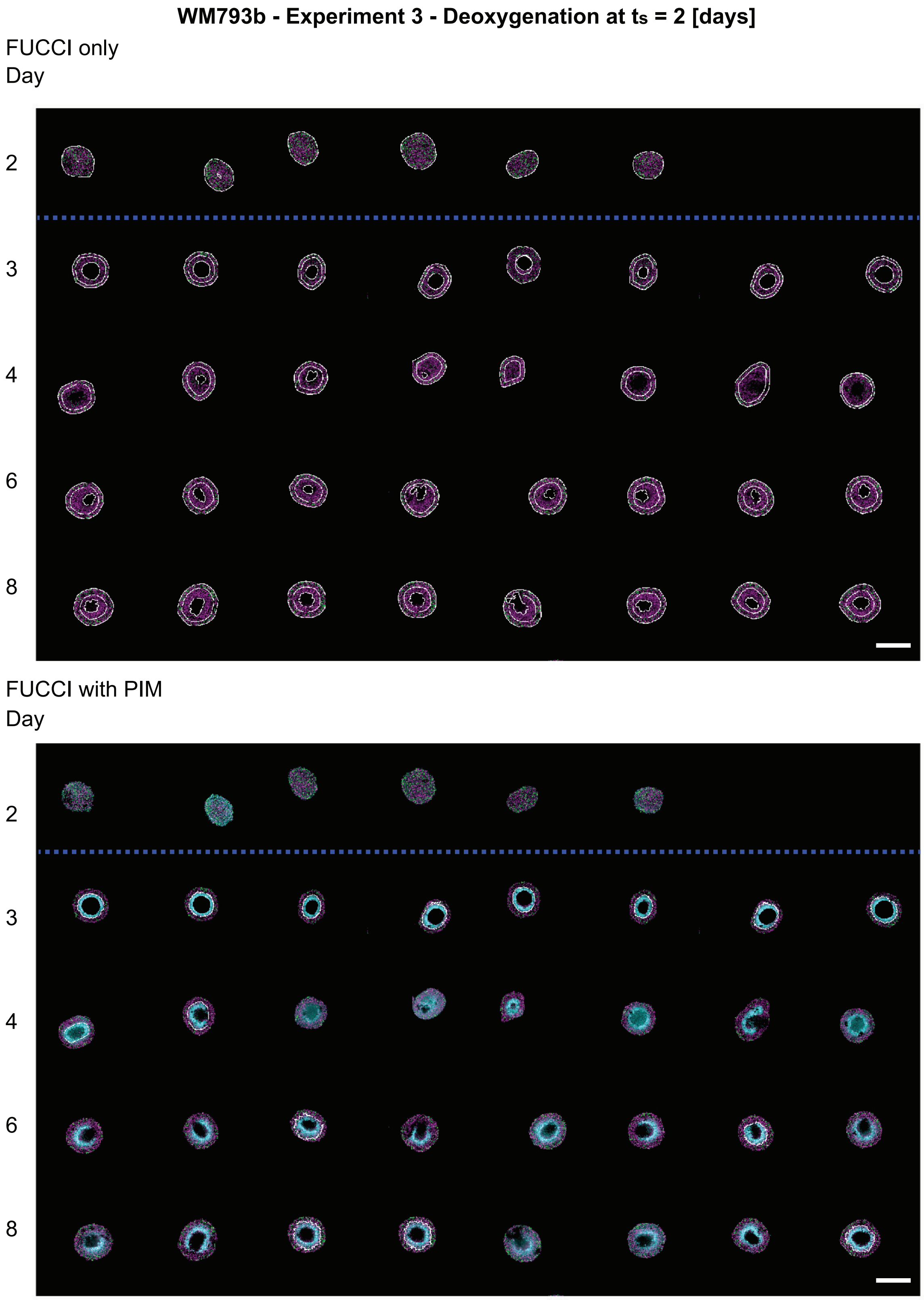


Figure S8: Experimental images of WM793b tumour spheroids in Experiment 3 - deoxygenation at $t_s = 2$ [days] (blue dashed line). Top set of images shows spheroids with FUCCI signal only. Bottom set of images show spheroids with FUCCI signal and pimonidazole staining. Scale bars are 400µm.

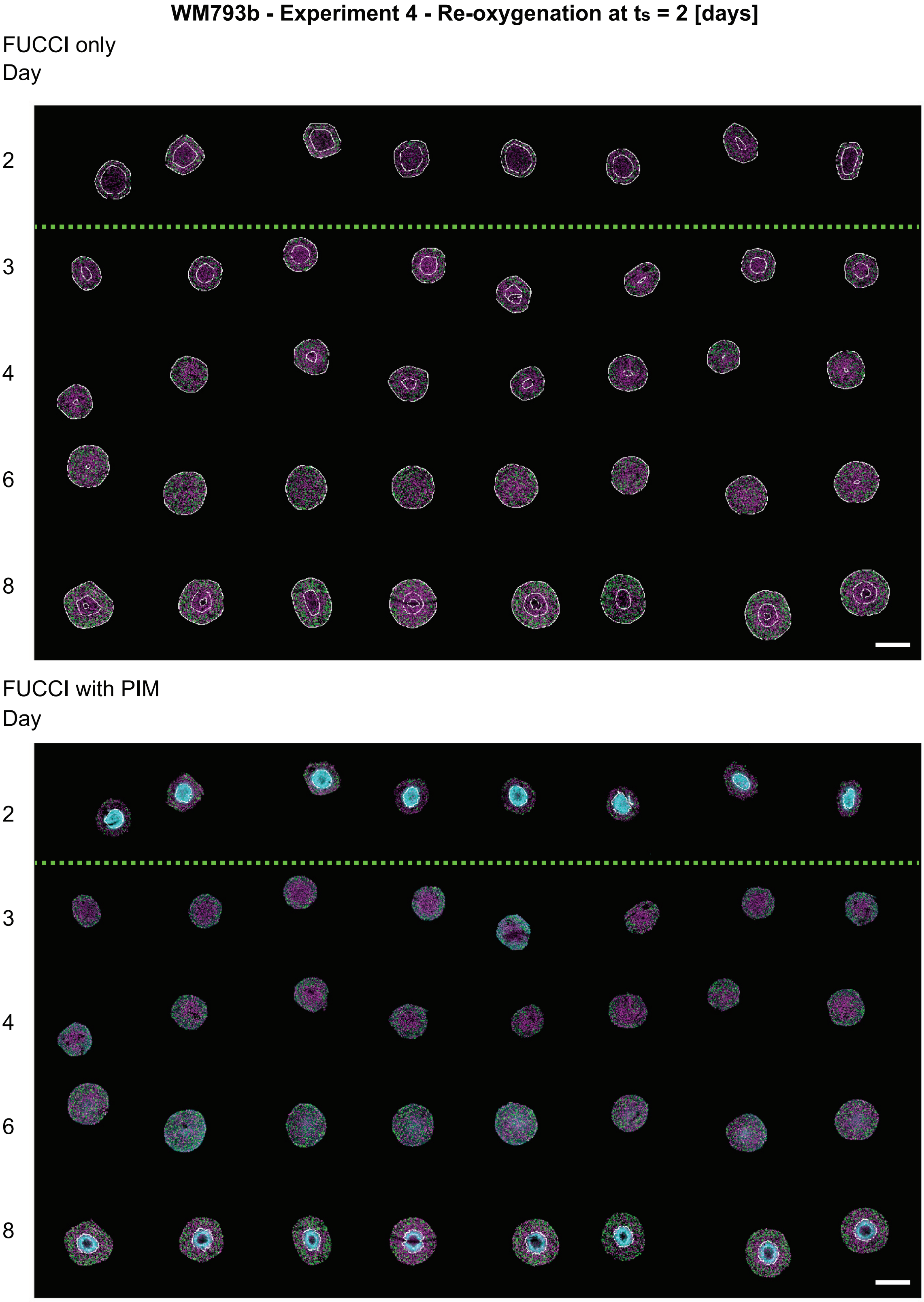


Figure S9: Experimental images of WM793b tumour spheroids in Experiment 4 - Re-oxygenation at $t_s = 2$ [days] (green dashed line). Top set of images shows spheroids with FUCCI signal only. Bottom set of images show spheroids with FUCCI signal and pimonidazole staining. Scale bars are 400µm.

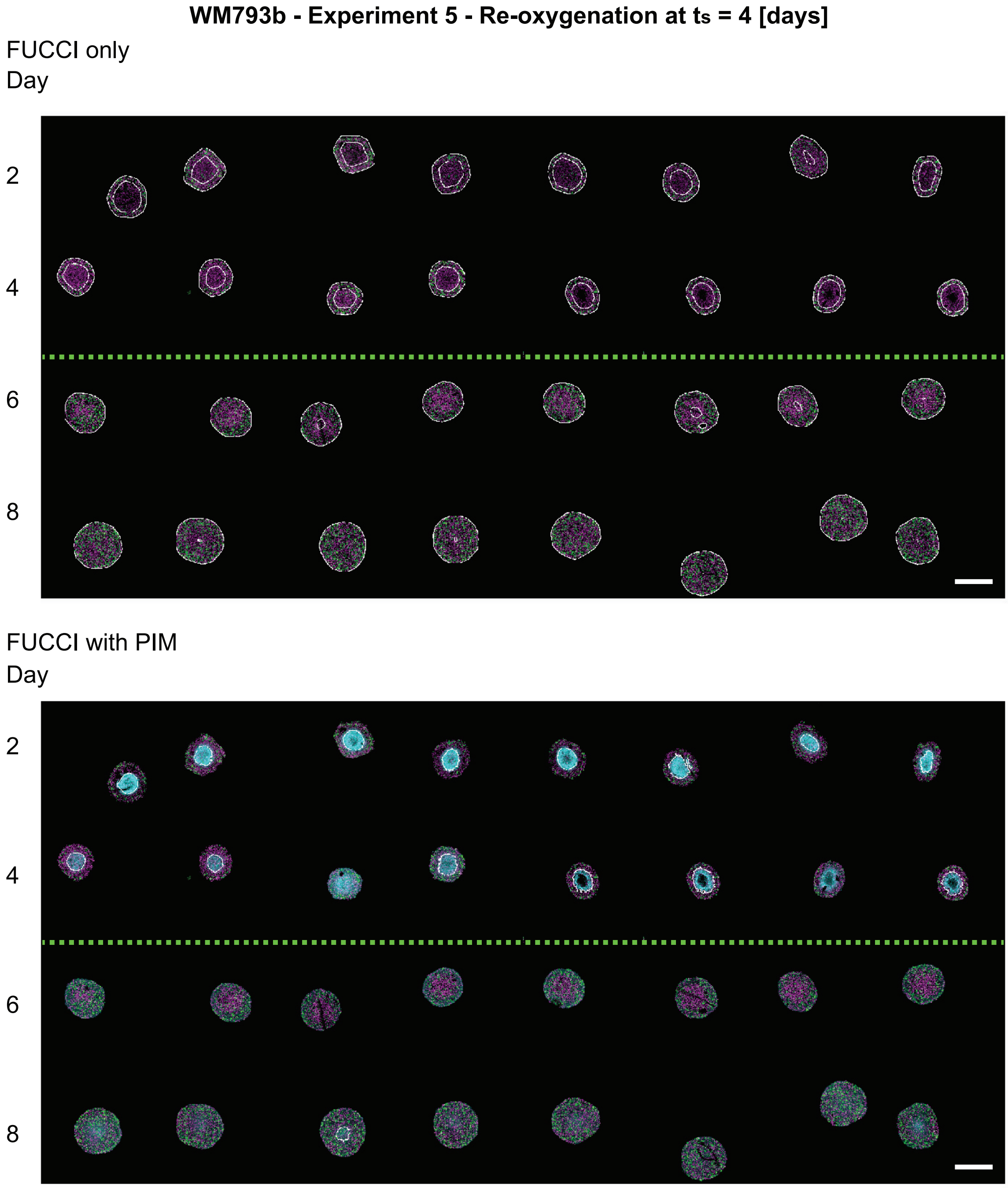


Figure S10: Experimental images of WM793b tumour spheroids in Experiment 5 - Re-oxygenation at $t_s = 4$ [days] (green dashed line). Top set of images shows spheroids with FUCCI signal only. Bottom set of images show spheroids with FUCCI signal and pimonidazole staining. Scale bars are 400µm.

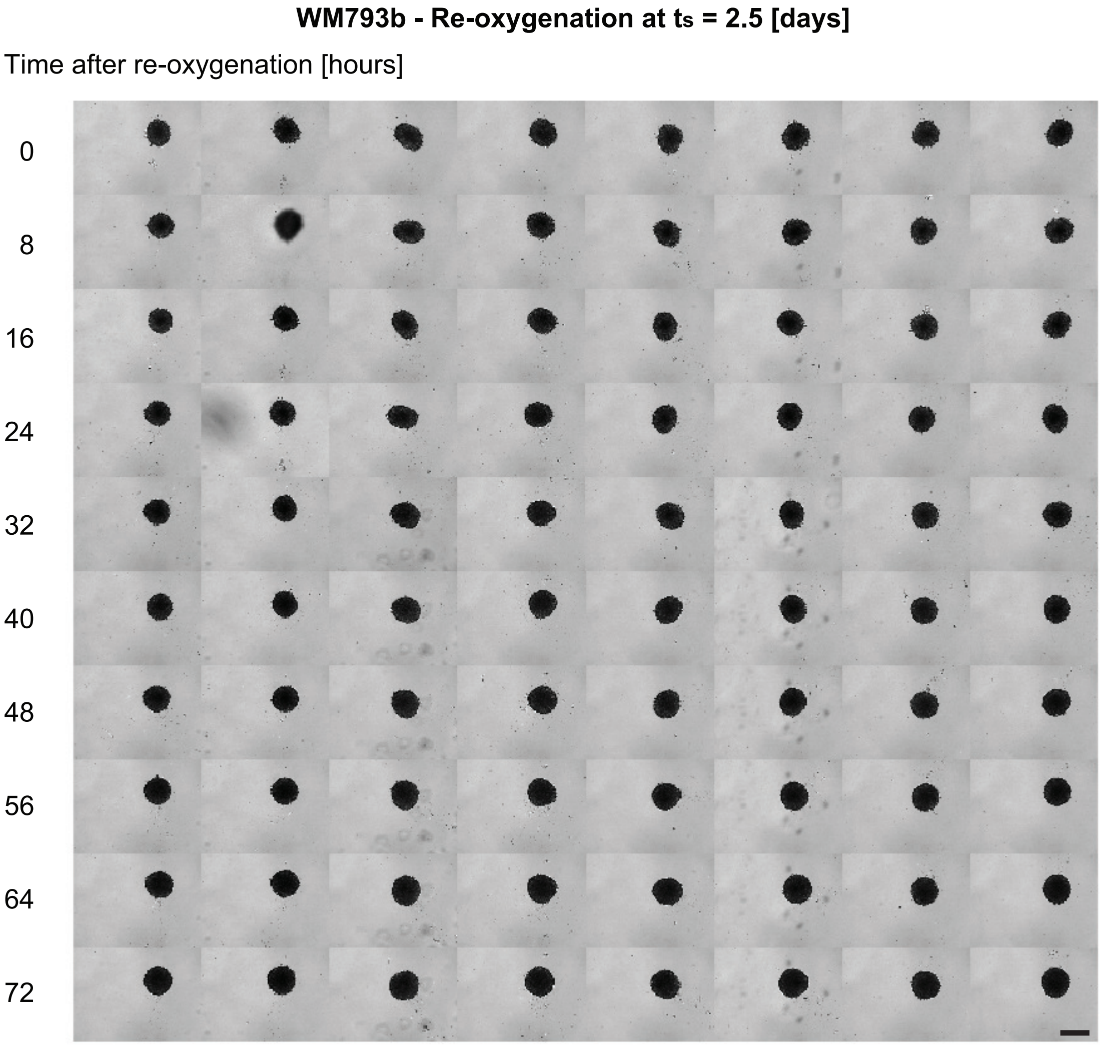


Figure S11: Experimental images of WM793b tumour spheroids in Experiment 6 - Re-oxygenation at $t_s = 2.5$ [days] (green dashed line). Scale bars are 400µm.

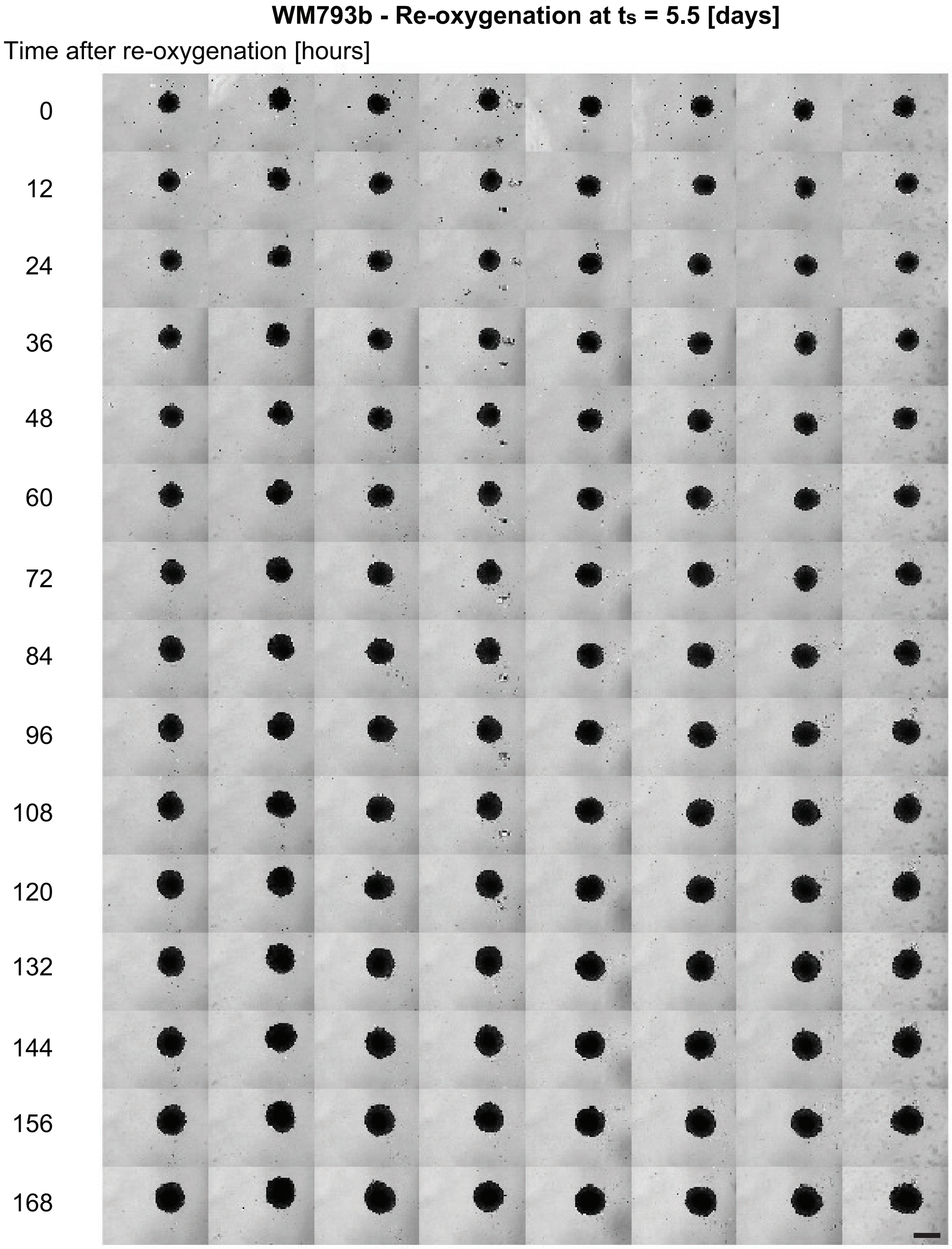


Figure S12: Experimental images of WM793b tumour spheroids in Experiment 7 - Re-oxygenation at $t_s = 5.5$ [days] (green dashed line). Scale bars are 400µm.

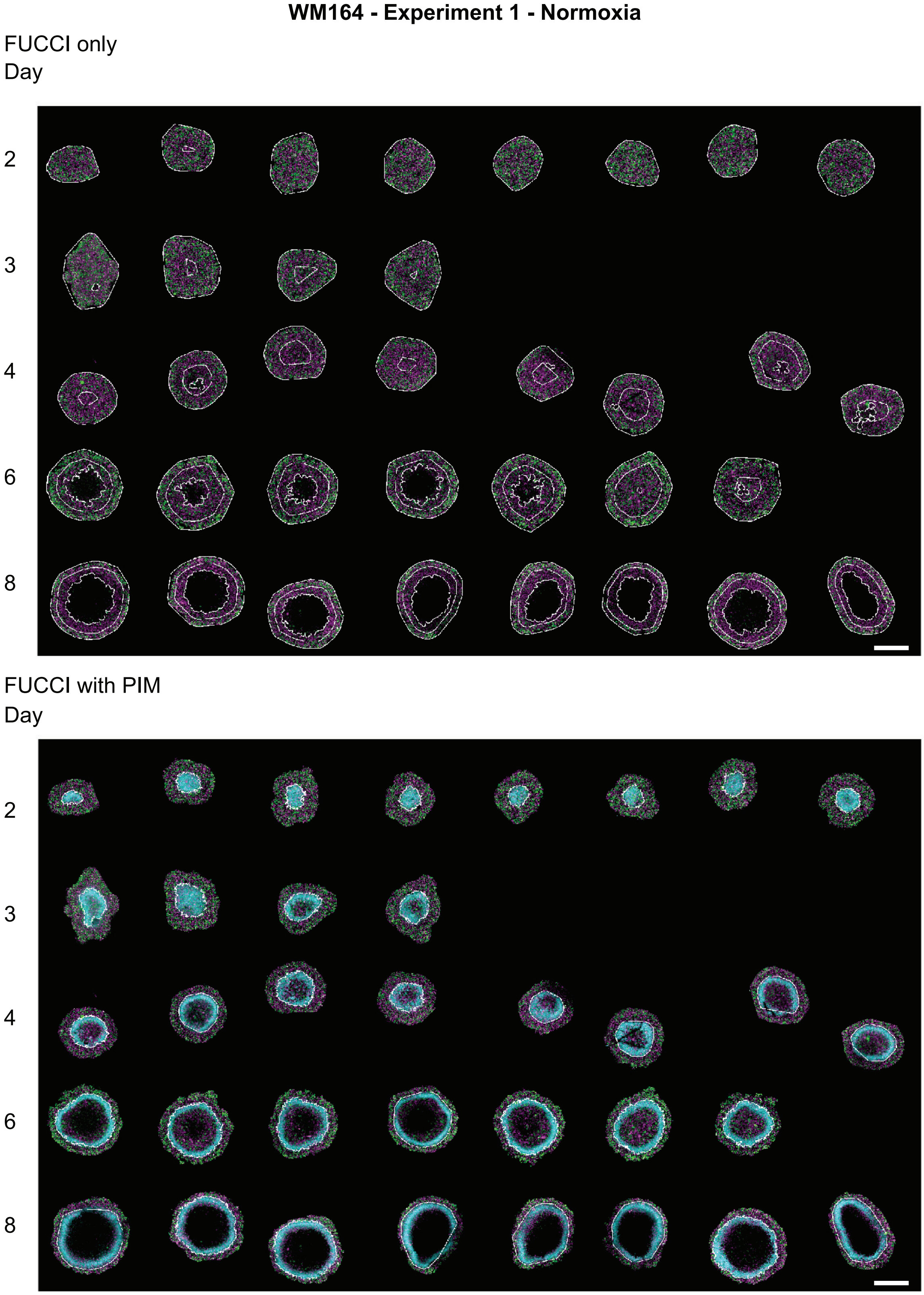


Figure S13: Experimental images of WM164 tumour spheroids in Experiment 1 - Normoxia. Top set of images shows spheroids with FUCCI signal only. Bottom set of images show spheroids with FUCCI signal and pimonidazole staining. Scale bars are 400µm.

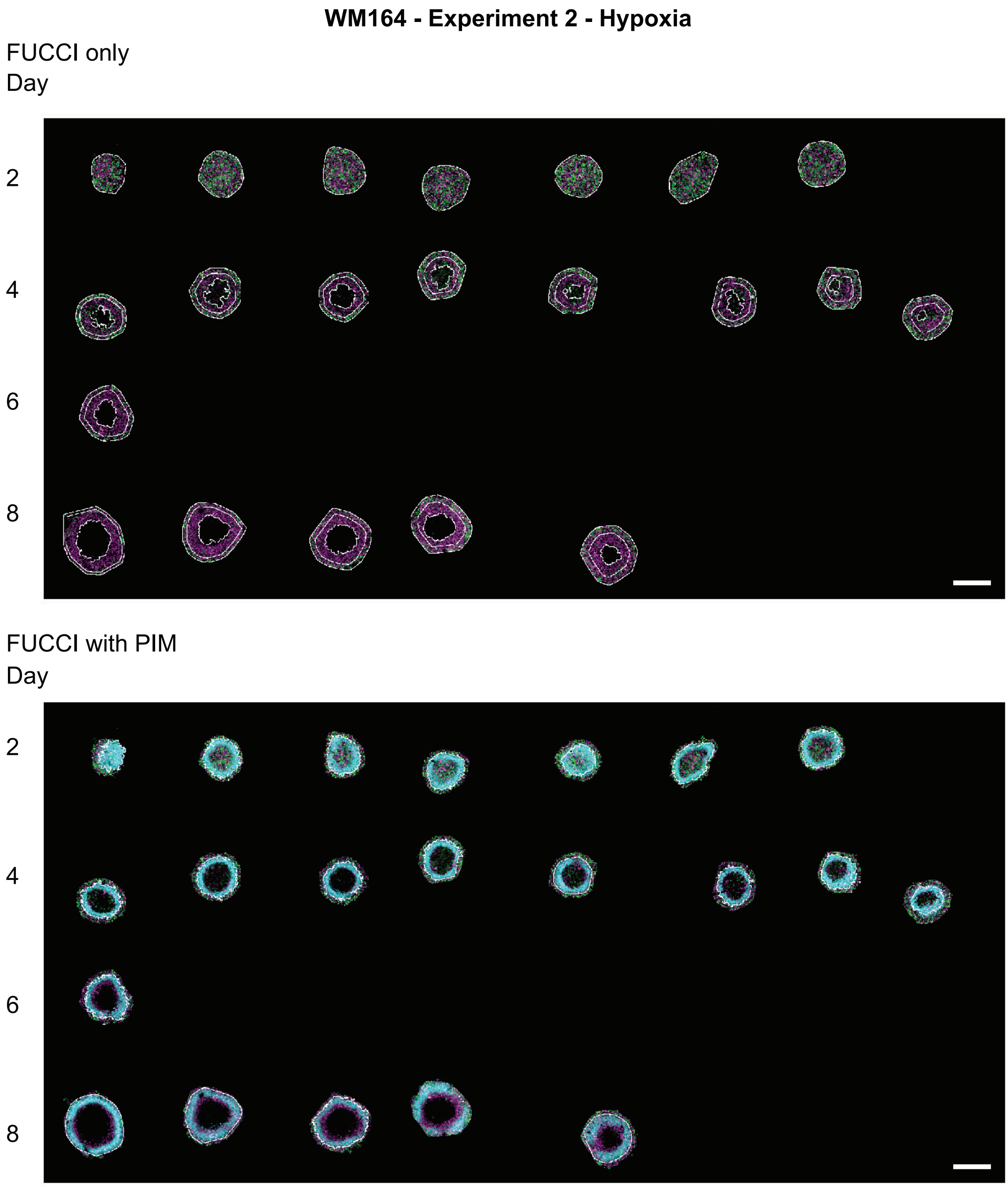


Figure S14: Experimental images of WM164 tumour spheroids in Experiment 2 - hypoxia. Top set of images shows spheroids with FUCCI signal only. Bottom set of images show spheroids with FUCCI signal and pimonidazole staining. Scale bars are 400μm.

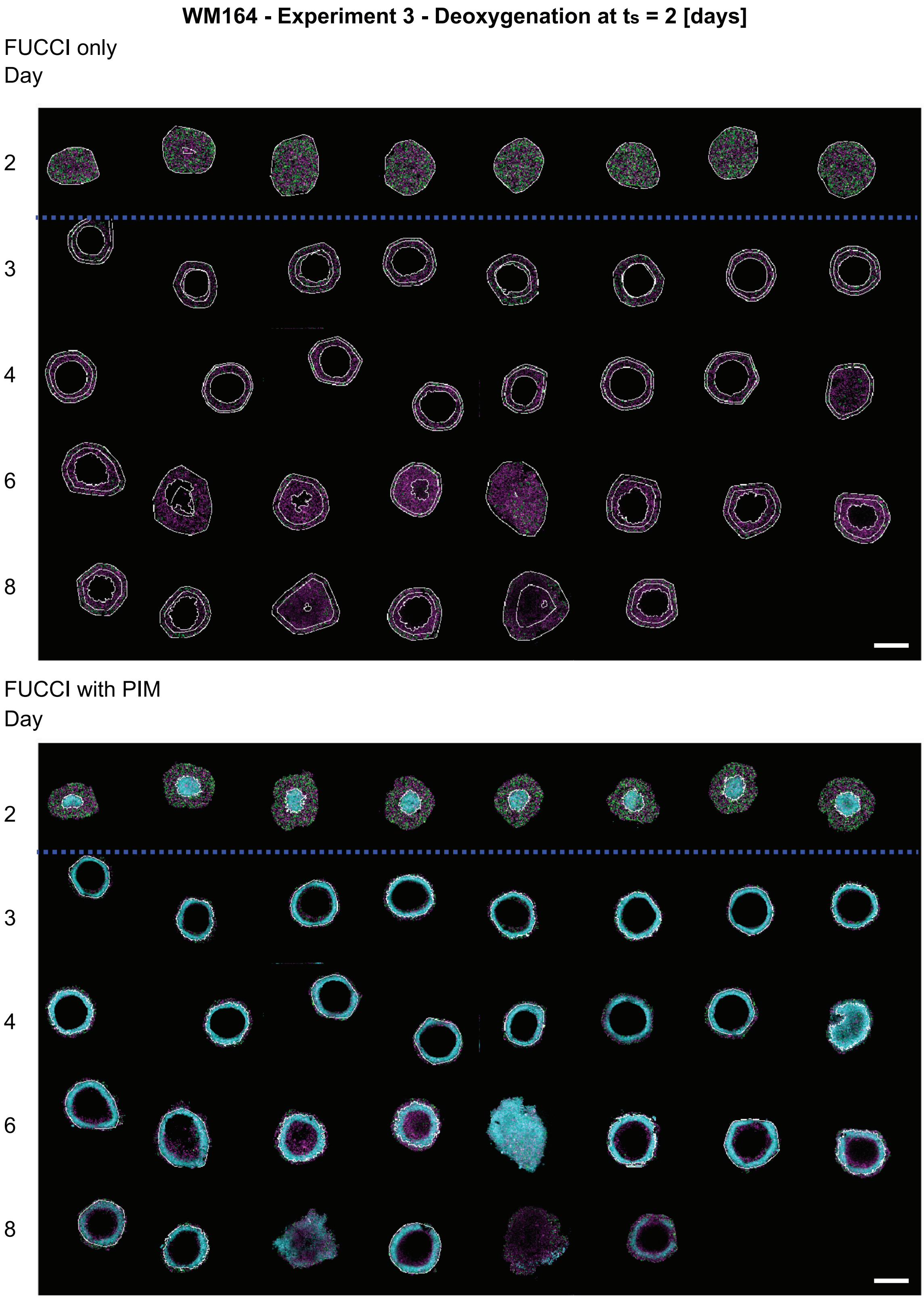


Figure S15: Experimental images of WM164 tumour spheroids in Experiment 3 - deoxygenation at $t_s = 2$ [days] (blue dashed line). Top set of images shows spheroids with FUCCI signal only. Bottom set of images show spheroids with FUCCI signal and pimonidazole staining. Scale bars are 400µm.

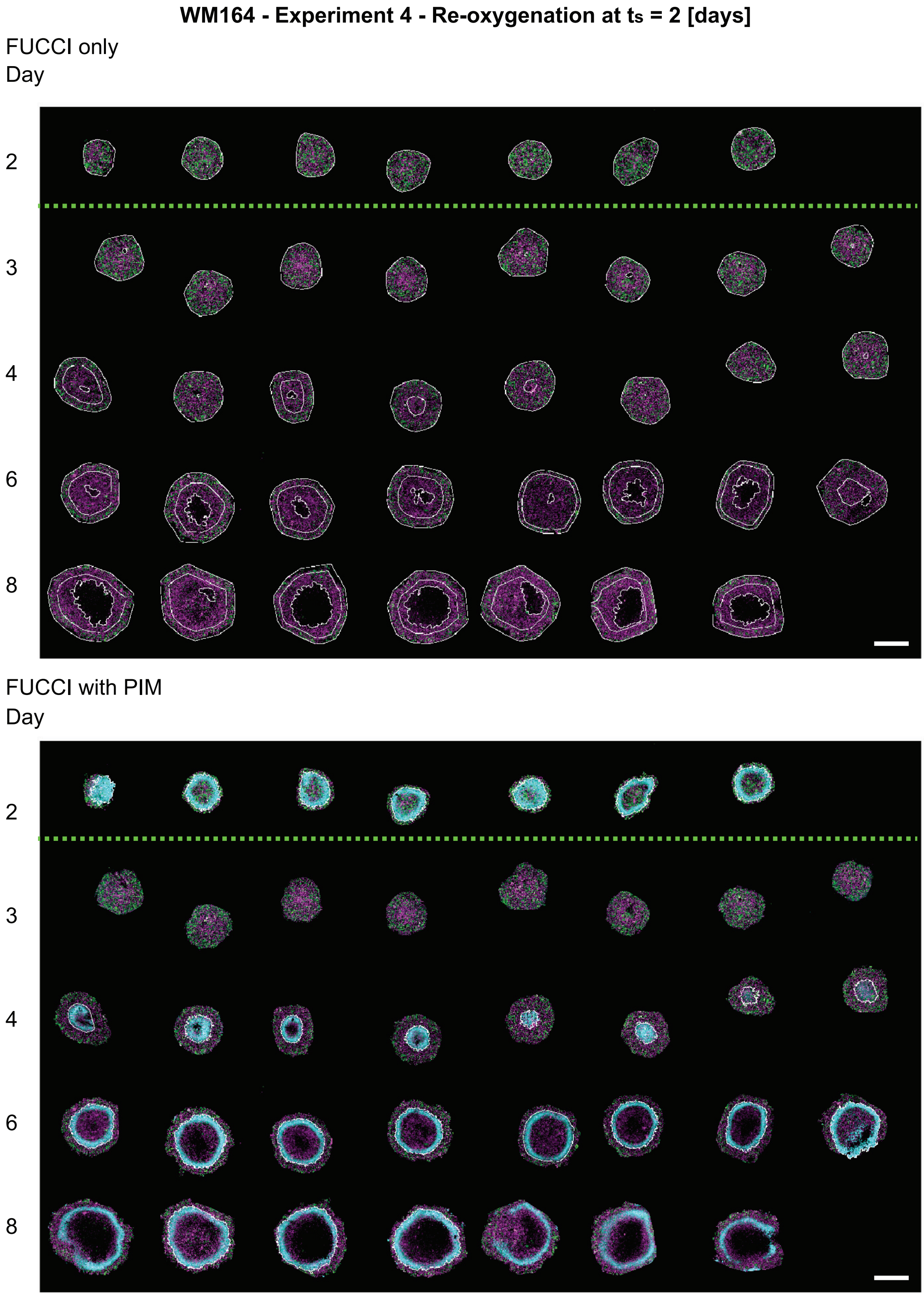


Figure S16: Experimental images of WM164 tumour spheroids in Experiment 4 - Re-oxygenation at $t_s = 2$ [days] (green dashed line). Top set of images shows spheroids with FUCCI signal only. Bottom set of images show spheroids with FUCCI signal and pimonidazole staining. Scale bars are 400µm.

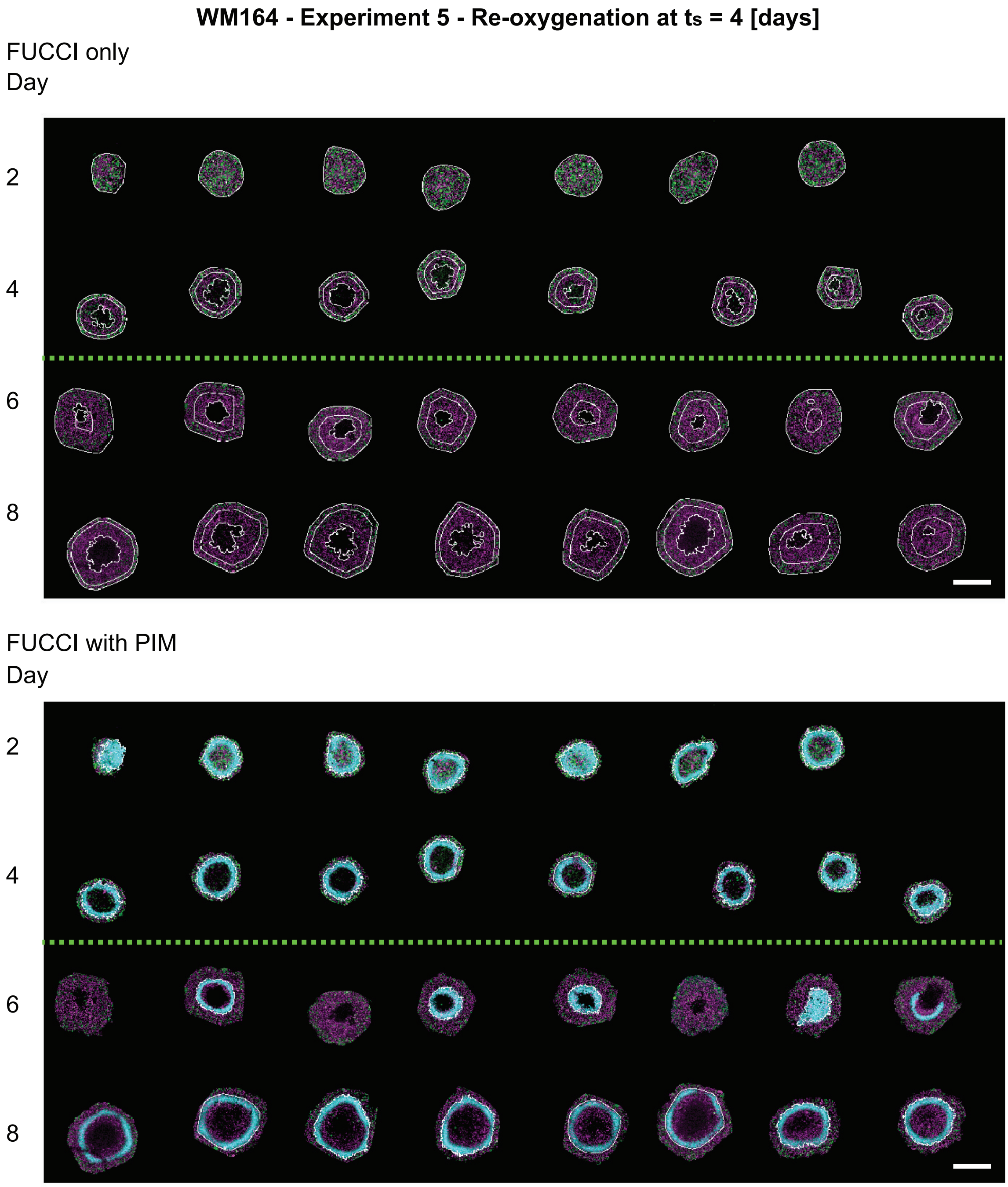


Figure S17: Experimental images of WM164 tumour spheroids in Experiment 5 - Re-oxygenation at $t_s$ = 4 [days] (green dashed line). Top set of images shows spheroids with FUCCI signal only. Bottom set of images show spheroids with FUCCI signal and pimonidazole staining. Scale bars are 400µm.

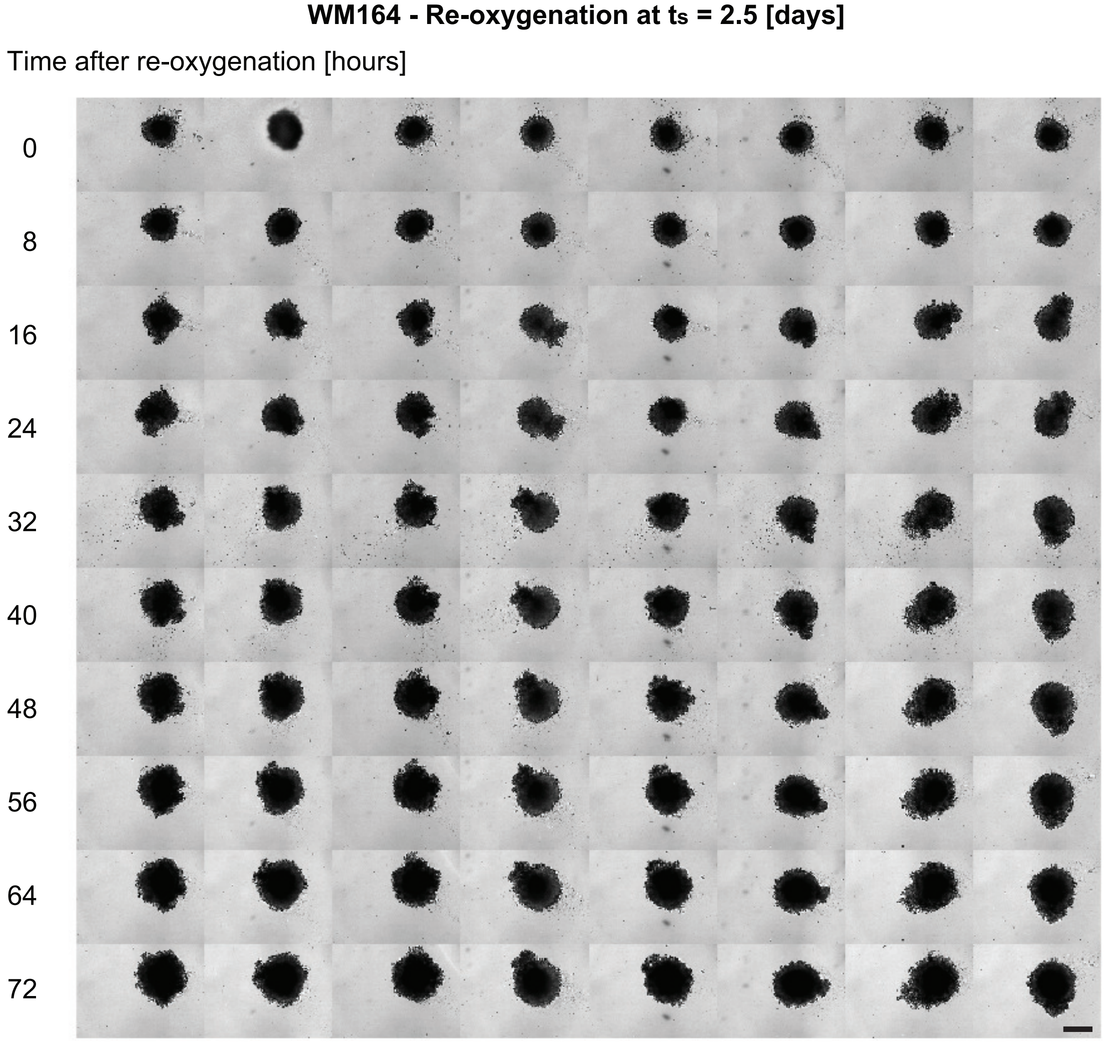


Figure S18: Experimental images of WM164 tumour spheroids in Experiment 6 - Re-oxygenation at $t_s = 2.5$ [days] (green dashed line). Scale bars are 400μm.

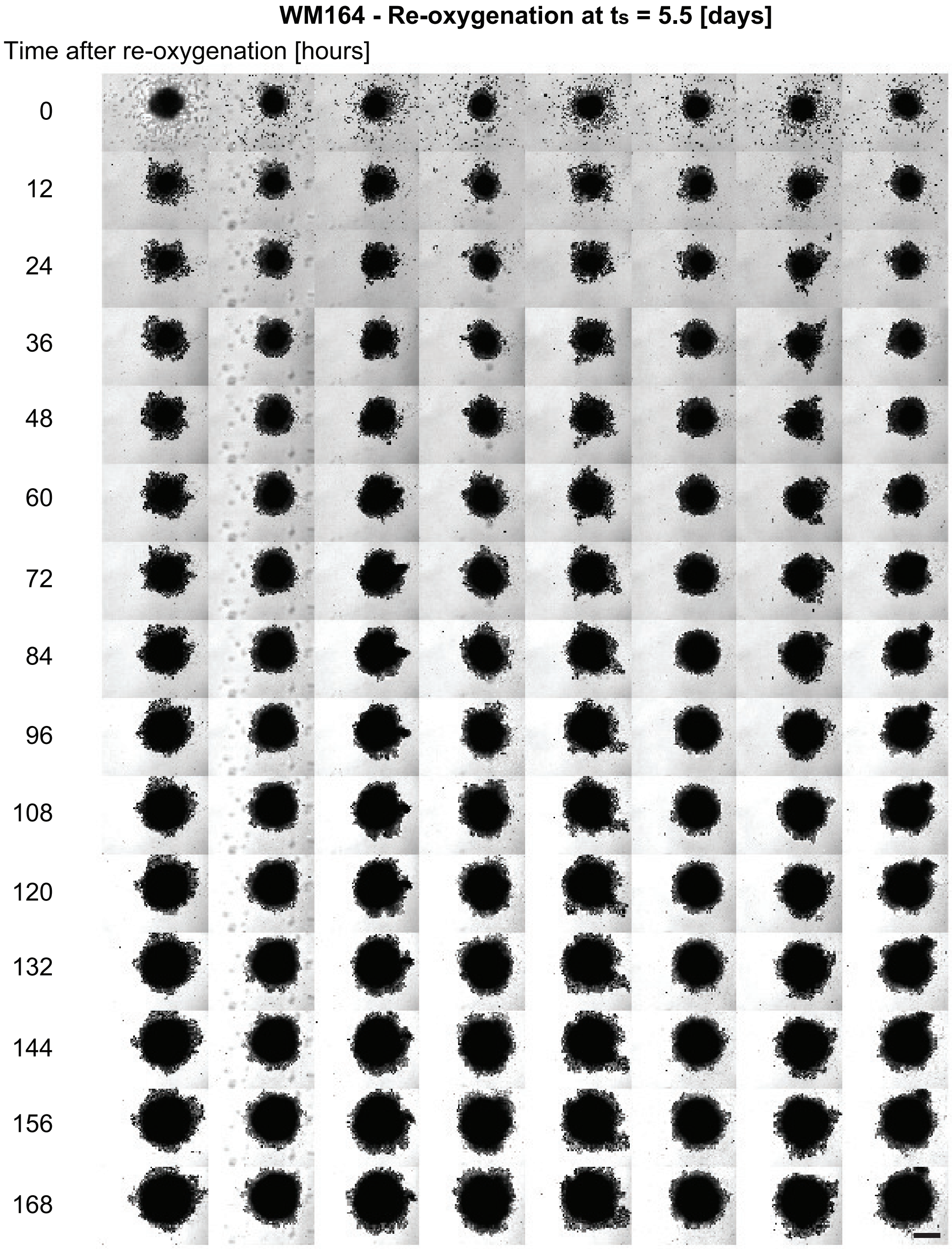


Figure S19: Experimental images of WM164 tumour spheroids in Experiment 7 - Re-oxygenation at $t_s = 5.5$ [days] (green dashed line). Scale bars are 400µm.

### B Image processing additional details

To estimate $R_\mathrm{o}(t)$, $R_\mathrm{n}(t)$ and $R_\mathrm{i}(t)$ we use MATLAB scripts that are freely available on Zenodo with DOI:10.5281/zenodo.5121093 [1]. These MATLAB scripts have been tested, developed, and used in previous studies on similar spheroid images [2–4]. Results from the image processing are manually inspected after using the scripts. For hypoxia and deoxygenation experiments the image processing scripts do not always accurately capture $R_\mathrm{n}(t)$. Therefore, we include an additional pre-processing step.

In deoxygenation experiments, and sometimes in hypoxia experiments, we observe that the FUCCI signal in the central region of the spheroid is blurred (Figure S20a-b), indicating dying or dead cells. Therefore, we identify this central region as the necrotic core. To identify this blurred region we open the relevant spheroid image in ImageJ, select the red FUCCI channel (shown in magenta), and draw a polygon with twenty points using the *polygon section* tool (Figure S20d). For accuracy, we compare the polygon with the FUCCI signal from the green channel and pimonidazole signal (cyan) (Figure S20c,d,e). Then we delete the polygon from the image (Figure S20f,g). This additional pre-processing step allows us to use MATLAB scripts on Zenodo with no further changes.

To estimate the hypoxic radius, $R_\mathrm{p}(t)$, we adapt existing MATLAB code on Zenodo [1] to account for the additional channel used to detect the pimonidazole staining. Specifically, we adapt the code used to identify the spheroids outer boundary and estimate $R_\mathrm{o}(t)$. Due to gradients in the pimonidazole staining, as opposed to the sharp transition at the edge of the spheroid, we adjust the signal boost from 1.1 to 2. Further, we perform the image processing with and without standard deviation filtering. We manually choose between the image processing results with and without standard deviation filtering to accurately identify the hypoxic region. Standard deviation filtering is used to estimate $R_\mathrm{o}(t)$ and so we use this option when results are similar. For a few images both methods do not detect the hypoxic region, due to the gradient in the signal. For these images we use the ImageJ *polygon section* and *measure* tools to estimate the hypoxic region.

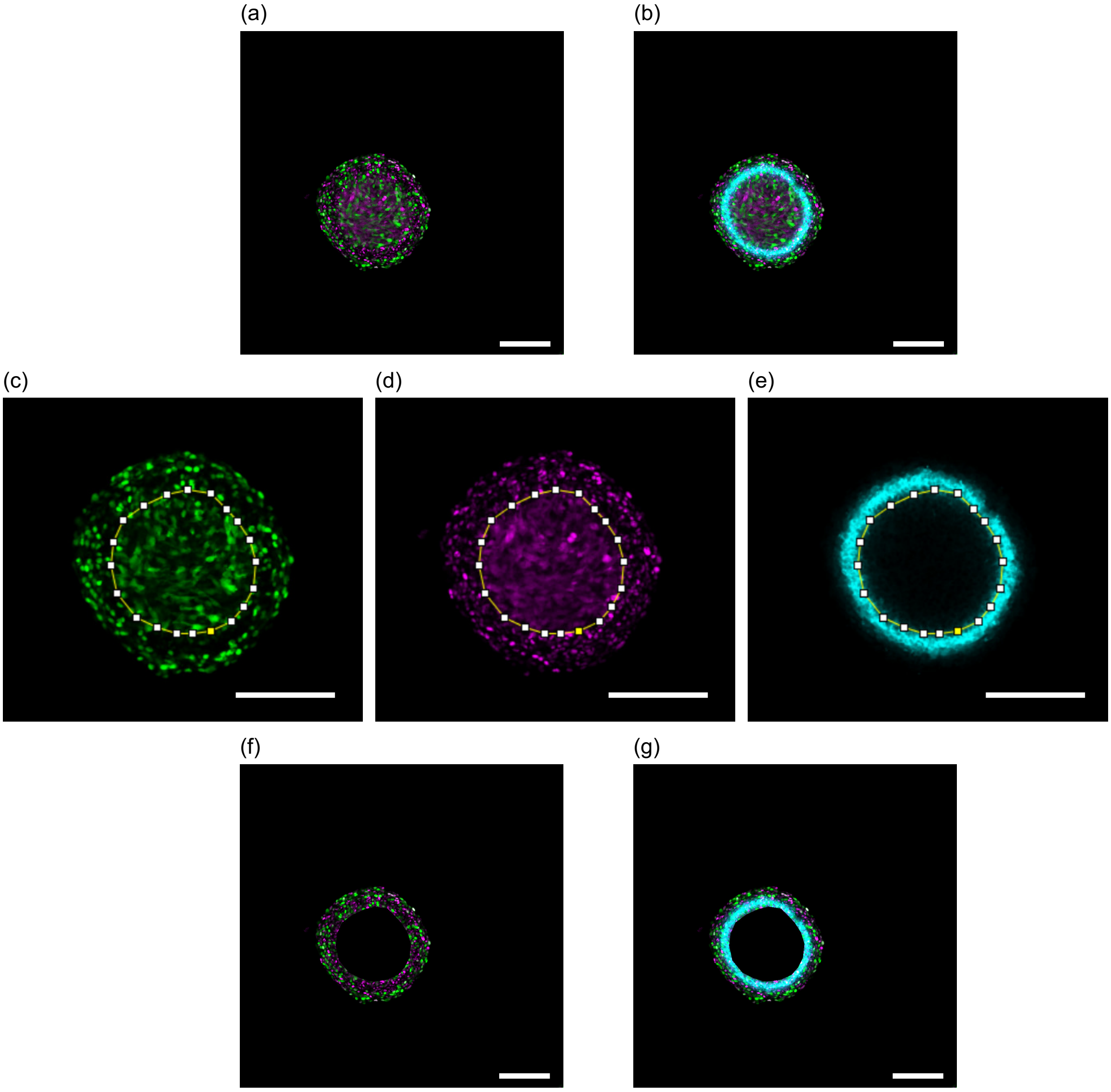


Figure S20: Image processing to estimate $R_\text{n}(t)$ when the FUCCI signal in the central region of the spheroid is blurred. Images show one WM983b spheroid one day after deoxygenation. Scale bars are 200 µm. (a) Spheroid with FUCCI signals (green and magenta). (b) Spheroid with FUCCI signals and pimonidazole staining (cyan). (c-e) Blurred region identified using the *polygon section* tool in ImageJ. Spheroid with: (c) FUCCI-green signal only (green), (d) FUCCI-red signal only (magenta), and (e) pimonidazole staining only (cyan). (f-g) Images in (a) and (b) with blurred region removed.

### C Mathematical modelling additional details

#### C.1 Greenspan's mathematical model

##### C.1.1 Model derivation

Key details are outlined in the main manuscript. Here, we provide further details about Greenspan's mathematical model [5] to describe spheroid growth in oxygen conditions, such as normoxia and hypoxia. The model is derived by considering conservation of volume,

$$A = B + C - D - E, \tag{S.1}$$

where $A$ is the total volume of the living cells at time $t$; $B$ is the initial volume of living cells; $C$ is the total volume of living cells produced in $t \geq 0$; $D$ is the volume of the necrotic core at time $t$ and, $E$ is the total volume lost from the necrotic core in $t \geq 0$. Writing $A$, $B$, $C$, $D$, and $E$ in their mathematical forms gives, recalling that the surface area and volume of a sphere of radius, $r$, are $4\pi r^2$ and $4\pi r^3/3$, respectively,

$$A = \frac{4\pi}{3}\left(R_{\mathrm{o}}^3(t) - R_{\mathrm{n}}^3(t)\right), \tag{S.2.1}$$

$$B = \frac{4\pi}{3} R_{\mathrm{o}}^3(0), \tag{S.2.2}$$

$$C = 4\pi \int_0^t \int_{R_{\mathrm{i}}(t)}^{R_{\mathrm{o}}(t)} s r^2 \; \mathrm{d}r \; \mathrm{d}t, \tag{S.2.3}$$

$$D = \frac{4\pi}{3} R_{\mathrm{n}}^3(t), \tag{S.2.4}$$

$$E = \frac{4\pi}{3} \int_0^t 3\lambda R_{\mathrm{n}}^3(t) \; \mathrm{d}t, \tag{S.2.5}$$

Substituting Equations (S.2.1)-(S.2.5) into Equation (S.1), differentiating with respect to time and simplifying gives governing Equation (1).

*Oxygen partial pressure governing equation and solution.* In Greenspan's mathematical model [5] oxygen is reported in terms of oxygen concentration and is assumed to be consumed by living cells at a constant rate. However, oxygen is typically reported in the experimental literature in terms of oxygen partial pressure [6]. To be consistent with the experimental literature and to avoid dimensional inconsistencies [7] we choose to present results in terms of oxygen partial pressure. Following the work of Grimes et al. [7], we define $c(r,t)$ [m$^3$ kg$^{-1}$] as the volume of oxygen gas per unit tumour mass. We assume that oxygen diffuses within the spheroid with diffusivity $k$ [m$^2$ s$^{-1}$], and that the rate of volume of oxygen gas per unit tumour mass that is consumed by living cells is a constant $\alpha$ [m$^3$ kg$^{-1}$ s$^{-1}$]. The volume of oxygen gas per unit tumour mass is assumed to be at diffusive equilibrium at all times so we write $c(r,t) = c(r)$. However, as $R_{\mathrm{o}}(t)$ is growing, oxygen diffusion occurs on a growing domain and we write $c(r) = c(r(t))$. The equation governing the volume of oxygen gas per unit tumour mass within the spheroid is,

$$\frac{1}{r^2}\frac{\partial}{\partial r}\left(r^2\frac{\partial}{\partial r}c(r(t))\right) = \frac{\alpha}{k}\mathrm{H}\left(r - R_{\mathrm{n}}(t)\right)\mathrm{H}\left(R_{\mathrm{o}}(t) - r\right), \quad 0 < r < R_{\mathrm{o}}(t). \tag{S.3}$$

We convert the volume of oxygen gas per unit tumour mass to partial pressure using Henry's law [7],

$$p(r(t)) = \Omega c(r(t)) \text{ [mmHg]}, \tag{S.4}$$

where $\Omega = \rho_T \rho_{\mathrm{O}_2} K = 3.0318 \times 10^7$[mmHg kgm$^{-3}$] is composed of: $\rho_T$, the density of the tumour that we assume is similar to that of water $k = 2 \times 10^{-9}$ [m$^2$ s$^{-1}$]; $\rho_{\mathrm{O}_2}$, the density of oxygen gas 1.331 [kg m$^{-3}$]; and, $K$, Henry's law constant which for oxygen gas at human body temperature, consistent with incubator settings (37 °C), is $2.2779 \times 10^{-4}$[m$^3$ mmHg kg$^{-1}$] [7]. Then we rewrite Equation (S.4) in terms of oxygen partial pressure to obtain Equation (2), rewritten here for clarity,

$$\frac{1}{r^2}\frac{\partial}{\partial r}\left(r^2\frac{\partial}{\partial r}p(r(t))\right) = \frac{\Omega\alpha}{k}\mathrm{H}\left(r - R_{\mathrm{n}}(t)\right)\mathrm{H}\left(R_{\mathrm{o}}(t) - r\right), \quad 0 < r < R_{\mathrm{o}}(t), \tag{S.5}$$

with external oxygen partial pressure $p_\infty$ [mmHg]. Oxygen partial pressure is commonly reported in units of percentage of standard atmospheric pressure, and we choose to do so here for consistency with standard cell culture incubators settings, using the conversion 160 mmHg = 21 %. For normoxic oxygen conditions $p_\infty = 21\%$ and for hypoxic conditions $p_\infty = 2\%$. Further, the hypoxic region is indicated by activation of pimonidazole at oxygen partial pressures of 1.32% [8].

The solution of Equation (S.5) is,

$$p(r(t)) = \begin{cases} p_\infty - \dfrac{\alpha\Omega}{6k}\left(R_{\mathrm{o}}^2(t) - r^2\right) + \dfrac{\alpha\Omega R_{\mathrm{n}}^3(t)}{3k}\left(\dfrac{1}{r} - \dfrac{1}{R_{\mathrm{o}}(t)}\right), & R_{\mathrm{n}}(t) \le r \le R_{\mathrm{o}}(t), \\ p_{\mathrm{n}}, & 0 \le r \le R_{\mathrm{n}}(t), \end{cases} \tag{S.6}$$

where

$$p_\infty - p_{\mathrm{n}} = \frac{\alpha\Omega}{6k}\left[R_{\mathrm{o}}^2(t) - R_{\mathrm{n}}^2(t) - 2\frac{R_{\mathrm{n}}^2(t)}{R_{\mathrm{o}}(t)}\left(R_{\mathrm{o}}(t) - R_{\mathrm{n}}(t)\right)\right]. \tag{S.7}$$

We set $p_{\mathrm{n}} = 0$ [7]. Hypothesis 1 and 2 both assume that $R_{\mathrm{n}}(t)$ is implicitly defined as the greatest radial position when the oxygen partial pressure is equal to zero. Specifically $p(R_{\mathrm{n}}(t)) = 0$ provided the oxygen partial pressure is sufficiently small, and $R_{\mathrm{n}}(t) = 0$ otherwise. Next we introduce a convenient parameter: the outer radius when the necrotic region first forms $R_{\mathrm{c}}$, defined as

$$R_{\mathrm{c}}^2 = \frac{6kp_\infty}{\alpha\Omega}. \tag{S.8}$$

Using Equation (S.8) we rewrite Equation (S.7) as

$$R_{\mathrm{c}}^2 = R_{\mathrm{o}}^2(t) - R_{\mathrm{n}}^2(t) - \frac{2R_{\mathrm{n}}^2(t)}{R_{\mathrm{o}}(t)} \left(R_{\mathrm{o}}(t) - R_{\mathrm{n}}(t)\right). \tag{S.9}$$

Equation (S.9) is convenient to estimate $R_{\mathrm{c}}$ given measurements of $R_{\mathrm{o}}(t)$ and $R_{\mathrm{n}}(t)$, provided $R_{\mathrm{n}}(t) > 0$. Then we can estimate $\alpha$ given estimates of $R_{\mathrm{c}}$, $\Omega$, $k$, and $p_\infty$ and by rearranging to solve for $\alpha$ in Equation (S.8).

Hypothesis 1 also assumes that oxygen diffusion and consumption describes the formation and growth of the inhibited region. Specifically that $R_{\mathrm{i}}(t)$ is implicitly defined by $p(R_{\mathrm{i}}(t)) = p_{\mathrm{i}}$, provided the oxygen partial pressure is sufficiently small, and $R_{\mathrm{i}}(t) = 0$ otherwise. Evaluating Equation (S.6) at $R_{\mathrm{i}}(t)$, provided $R_{\mathrm{i}}(t) > R_{\mathrm{n}}(t)$, gives the following convenient equation to explore hypothesis 1,

$$\mathcal{R}^2 = R_{\mathrm{o}}^2(t) - R_{\mathrm{i}}^2(t) - 2R_{\mathrm{n}}^3(t) \left(\frac{1}{R_{\mathrm{i}}(t)} - \frac{1}{R_{\mathrm{o}}(t)}\right), \tag{S.10}$$

where we have defined the outer radius when the inhibited region first forms as $\mathcal{R}$ defined as,

$$\mathcal{R}^2 = \frac{6k\left(p_\infty - p_{\mathrm{i}}\right)}{\alpha\Omega}. \tag{S.11}$$

*Waste concentration governing equation and solution.* Hypothesis 2 proposes that production of waste from living cells and diffusion of waste within the spheroid is responsible for the formation and growth of the inhibited region. Following Greenspan's original model [5] we report waste within the spheroid in terms of waste concentration. Equation (3) governs the time evolution of waste concentration within the spheroid, $\beta(r(t))$, and the solution is,

$$\beta(r(t)) = \begin{cases} \dfrac{P}{6\kappa}\left[R_{\mathrm{o}}^2(t) - r^2 - 2R_{\mathrm{n}}^3(t)\left(\dfrac{1}{r} - \dfrac{1}{R_{\mathrm{o}}(t)}\right)\right], & R_{\mathrm{n}}(t) \le r \le R_{\mathrm{o}}(t), \\ \dfrac{P}{6\kappa}\left[R_{\mathrm{o}}^2(t) - R_{\mathrm{n}}^2(t) - 2R_{\mathrm{n}}^3(t)\left(\dfrac{1}{R_{\mathrm{n}}(t)} - \dfrac{1}{R_{\mathrm{o}}(t)}\right)\right], & 0 \le r \le R_{\mathrm{n}}(t). \end{cases} \tag{S.12}$$

The inhibited radius, $R_{\mathrm{i}}(t)$, is implicitly defined through $\beta_i = \beta(R_{\mathrm{i}}(t))$ provided the waste concentration is sufficiently large, and $R_{\mathrm{i}}(t) = 0$ otherwise. Evaluating Equation (S.12) at $R_{\mathrm{i}}(t)$, provided $R_{\mathrm{i}}(t) > R_{\mathrm{n}}(t)$, gives the following convenient equation to explore hypothesis 2,

$$\mathcal{R}^2 = R_{\mathrm{o}}^2(t) - R_{\mathrm{i}}^2(t) - 2R_{\mathrm{n}}^3(t) \left(\frac{1}{R_{\mathrm{i}}(t)} - \frac{1}{R_{\mathrm{o}}(t)}\right). \tag{S.13}$$

where we have defined the outer radius when the inhibited region first forms as $\mathcal{R}$ defined as,

$$\mathcal{R}^2 = \frac{6\beta_i\kappa}{P}. \tag{S.14}$$

##### C.1.2 Numerical methods

The code to simulate Greenspan's model to interpret normoxia and hypoxia experiments is available on the GitHub repository listed in Methods: Code Availability. Here, we outline the key details. First, we rewrite governing Equations (6.1)-(6.3),

$$R_{\mathrm{o}}^{2}(t)\frac{\mathrm{d}R_{\mathrm{o}}(t)}{\mathrm{d}t}=\frac{s}{3}\left[R_{\mathrm{o}}^{3}(t)-\max\left(R_{\mathrm{i}}(t)^{3},R_{\mathrm{n}}^{3}(t)\right)\right]-\lambda R_{\mathrm{n}}(t)^{3}, \tag{S.15.1}$$

$$R_{\mathrm{c}}^{2}=R_{\mathrm{o}}^{2}(t)-R_{\mathrm{n}}^{2}(t)-\frac{2R_{\mathrm{n}}^{2}(t)}{R_{\mathrm{o}}(t)}\left(R_{\mathrm{o}}(t)-R_{\mathrm{n}}(t)\right), \tag{S.15.2}$$

$$\mathcal{R}^{2}=R_{\mathrm{o}}^{2}(t)-R_{\mathrm{i}}^{2}(t)-2R_{\mathrm{n}}^{3}(t)\left(\frac{1}{R_{\mathrm{i}}(t)}-\frac{1}{R_{\mathrm{o}}(t)}\right). \tag{S.15.3}$$

Note that Equation (S.15.1) is an ordinary differential equation and Equations (S.15.2)-(S.15.3) are algebraic equations.

To numerically solve the coupled differential-algebraic system of Equations (S.15.1)-(S.15.3) we use MATLAB's `ode15s` function. At time $t$ we assume that $R_{\mathrm{o}}(t)$ is known. To determine $R_{\mathrm{n}}(t)$, $R_{\mathrm{i}}(t)$, and $R_{\mathrm{o}}(t+\Delta t)$, where the time step $\Delta t$ is determined by the `ode15s` function, we,

1. solve Equation (S.15.2) using MATLAB's `roots` function and define $R_{\mathrm{n}}(t)$ as the root between 0 and $R_{\mathrm{o}}(t)$, otherwise $R_{\mathrm{n}}(t)=0$;
2. solve Equation (S.15.3) using MATLAB's `roots` function and define $R_{\mathrm{i}}(t)$ as the root between `max` $(0, R_{\mathrm{n}}(t))$ and $R_{\mathrm{o}}(t)$, otherwise $R_{\mathrm{i}}(t)=0$;
3. compute $R_{\mathrm{o}}(t+\Delta t)$ using Equation (S.15.1) given $R_{\mathrm{n}}(t)$ and $R_{\mathrm{i}}(t)$.

Note that in experiments spheroids formed two days after seeding. As the model is only appropriate once the spheroids have formed, the initial time in the mathematical model corresponds to two days after seeding.

#### C.2 Mathematical model to interpret deoxygenation experiments

##### C.2.1 Model derivation

Key details are outlined in the main manuscript. Here we provide further details extending Greenspan's mathematical model to interpret the deoxygenation experiments. The conservation of volume argument from Greenspan's model used to derive Equation (1) still holds, now with time dependence captured in $s(t)$ and $\lambda(t)$.

*Necrotic core.* Considering conservation of volume for the necrotic core gives

$$A_{\mathrm{n}} = B_{\mathrm{n}} + C_{\mathrm{n}} - E, \tag{S.16}$$

where $A_{\mathrm{n}}$ is the total volume of the necrotic core at time $t$; $B_{\mathrm{n}}$ is the volume of the necrotic core at time $t_{\mathrm{s}}$; $C_{\mathrm{n}}$ is the total volume of necrotic core produced in $t \geq t_{\mathrm{s}}$ due to to cells dying in the region $R_{\mathrm{n}}(t) < r < R_{\mathrm{n}}^{+}(t)$ at a rate $\hat{\lambda}(t)$ per unit volume; and, $E$ is the total volume lost in the necrotic core in $t \geq t_{\mathrm{s}}$ defined in Equation (S.2) of Greenspan's original mathematical model. Note that in the definition of $C_{\mathrm{n}}$ we assume that the necrotic core volume only increases due to to cells dying in the region $R_{\mathrm{n}}(t) < r < R_{\mathrm{n}}^{+}(t)$. Therefore, $C_{\mathrm{n}}$ does not include live cells dying to replenish the volume loss in the necrotic centre at the boundary $R_{\mathrm{n}}(t)$ as in Greenspan's original model. This assumption is to simplify the analysis, and also implicitly assumes that cells in $r > R_{\mathrm{n}}^{+}(t)$ are subject to small oxygen partial pressures for some time before dying. This mechanism is also implicitly assumed in Greenspan's original mathematical model. Writing $A_{\mathrm{n}}$, $B_{\mathrm{n}}$, and $C_{\mathrm{n}}$ in their mathematical forms gives,

$$A_{\mathrm{n}} = \frac{4\pi}{3} R_{\mathrm{n}}^{3}(t), \tag{S.17.1}$$

$$B_{\mathrm{n}} = \frac{4\pi}{3} R_{\mathrm{n}}^{3}(t_{\mathrm{s}}), \tag{S.17.2}$$

$$C_{\mathrm{n}} = 4\pi \int_{t_{\mathrm{s}}}^{t} \int_{R_{\mathrm{n}}(t)}^{R_{\mathrm{n}}^{+}(t)} 3\hat{\lambda}(t) r^{2} \ \mathrm{d}r \ \mathrm{d}t, \tag{S.17.3}$$

where the three in Equation (S.17.3) is included for mathematical convenience. Substituting Equations (S.17) and Equation (S.2.5) into the conservation of volume Equation (S.16) gives

$$\frac{4\pi}{3} R_{\mathrm{n}}^{3}(t) = \frac{4\pi}{3} R_{\mathrm{n}}^{3}(t_{\mathrm{s}}) + 4\pi \int_{t_{\mathrm{s}}}^{t} \int_{R_{\mathrm{n}}(t)}^{R_{\mathrm{n}}^{+}(t)} 3\hat{\lambda}(t) r^{2} \ \mathrm{d}r \ \mathrm{d}t - \frac{4\pi}{3} \int_{t_{\mathrm{s}}}^{t} 3\lambda(t) R_{\mathrm{n}}^{3}(t) \ \mathrm{d}t, \tag{S.18}$$

Differentiating Equation (S.18) with respect to time and simplifying gives the governing equation for $R_{\mathrm{n}}(t)$,

$$R_{\mathrm{n}}^{2}(t) \frac{\mathrm{d}R_{\mathrm{n}}(t)}{\mathrm{d}t} = \hat{\lambda}(t) \left[ R_{\mathrm{n}}^{+}(t)^{3} - R_{\mathrm{n}}^{3}(t) \right] - \lambda(t) R_{\mathrm{n}}^{3}(t). \tag{S.19}$$

However, Equation (S.19) is valid only when $R_{\mathrm{n}}(t) > 0$. For a general governing equation, valid for

$R_\mathrm{n}(t) \geq 0$ we reexpress Equation (S.19) in terms of necrotic volume at time $t$,

$$\frac{\mathrm{d}V_\mathrm{n}(t)}{\mathrm{d}t} = 3\hat{\lambda}(t)\left[\frac{4\pi}{3}R_\mathrm{n}^{+}(t)^3 - V_\mathrm{n}(t)\right] - 3\lambda(t)V_\mathrm{n}(t), \tag{S.20}$$

and then define $R_\mathrm{n}(t)$ as

$$R_\mathrm{n}(t) = \left[\frac{3}{4\pi}V_\mathrm{n}(t)\right]^{\frac{1}{3}}. \tag{S.21}$$

We set $\hat{\lambda}(t) = \hat{\lambda}\exp\left((t-t_\mathrm{s})/\tau_{\hat{\lambda}}\right)$ in Equation (S.20) to capture the assumption that $R_\mathrm{n}(t) \to R_\mathrm{n}^{+}(t)$ as $t \to \infty$.

##### C.2.2 Numerical methods

The code to simulate the deoxygenation model is available on the GitHub repository listed in Methods: Code Availability. Here, we outline the key details. First, we rewrite governing Equations (8.1)-(8.11) for $t \geq t_\mathrm{s}$,

$$R_\mathrm{o}^2(t)\frac{\mathrm{d}R_\mathrm{o}(t)}{\mathrm{d}t} = \frac{s(t)}{3}\left[R_\mathrm{o}^3(t) - \max\left(R_\mathrm{i}^3(t), R_\mathrm{n}^3(t)\right)\right] - \lambda(t)R_\mathrm{n}^3(t), \tag{S.22.1}$$

$$\frac{\mathrm{d}V_\mathrm{n}(t)}{\mathrm{d}t} = 3\hat{\lambda}(t)\left[\frac{4\pi}{3}R_\mathrm{n}^{+}(t)^3 - V_\mathrm{n}(t)\right] - 3\lambda(t)V_\mathrm{n}(t), \tag{S.22.2}$$

$$R_\mathrm{c}^2(t) = R_\mathrm{o}^2(t) - R_\mathrm{n}^{+}(t)^2 - \frac{2R_\mathrm{n}^{+}(t)^2}{R_\mathrm{o}(t)}\left(R_\mathrm{o}(t) - R_\mathrm{n}^{+}(t)\right), \tag{S.22.3}$$

$$\mathcal{R}^2(t) = R_\mathrm{o}^2(t) - R_\mathrm{i}^2(t) - 2R_\mathrm{n}^3(t)\left(\frac{1}{R_\mathrm{i}(t)} - \frac{1}{R_\mathrm{o}(t)}\right), \tag{S.22.4}$$

$$\alpha(t) = \alpha_h + (\alpha_n - \alpha_h)\exp\left(-\frac{1}{\tau_\alpha}(t-t_\mathrm{s})\right), \tag{S.22.5}$$

$$\lambda(t) = \lambda_h + (\lambda_n - \lambda_h)\exp\left(-\frac{1}{\tau_\lambda}(t-t_\mathrm{s})\right), \tag{S.22.6}$$

$$s(t) = s_h + (s_n - s_h)\exp\left(-\frac{1}{\tau_s}(t-t_\mathrm{s})\right), \tag{S.22.7}$$

$$\mathcal{R}(t) = \mathcal{R}_h + (\mathcal{R}_n - \mathcal{R}_h)\exp\left(-\frac{1}{\tau_\mathcal{R}}(t-t_\mathrm{s})\right), \tag{S.22.8}$$

$$\hat{\lambda}(t) = \hat{\lambda}\exp\left(\frac{1}{\tau_{\hat{\lambda}}}(t-t_\mathrm{s})\right), \tag{S.22.9}$$

$$R_\mathrm{n}(t) = \left[\frac{3}{4\pi}V_\mathrm{n}(t)\right]^{\frac{1}{3}}, \tag{S.22.10}$$

$$R_\mathrm{c}^2(t) = \frac{6kp_\infty}{\alpha(t)\Omega}. \tag{S.22.11}$$

Note that: Equations (S.22.1) and (S.22.2) are ordinary differential equations; Equations (S.22.3), (S.22.4), and (S.22.10)-(S.22.11) are algebraic equations; Equations (S.22.5)-(S.22.8) include exponential decay; and Equation (S.22.9) describes exponential growth.

To numerically solve the coupled differential-algebraic system of Equations (S.22.1)-(S.22.11) we use MATLAB's `ode15s` function. At time $t$ we assume that $R_\mathrm{o}(t)$ and $R_\mathrm{n}(t)$ are known. To determine

$R_\mathrm{i}(t)$, $R_\mathrm{n}(t+\Delta t)$, and $R_\mathrm{o}(t+\Delta t)$, where the time step $\Delta t$ is determined by the `ode15s` function, we,

1. evaluate Equations (S.22.5)-(S.22.9) to obtain $\alpha(t)$, $\lambda(t)$, $s(t)$, $\mathcal{R}(t)$, and $\hat{\lambda}(t)$;
2. evaluate $R_\mathrm{c}^2(t)$ using Equation (S.22.11);
3. solve Equation (S.22.10) using MATLAB's `roots` function and define $R_\mathrm{n}^+(t)$ as the root between 0 and $R_\mathrm{o}(t)$, otherwise $R_\mathrm{n}^+(t) = 0$;
4. solve Equation (S.22.4) using MATLAB's `roots` function and define $R_\mathrm{i}(t)$ as the root between $\mathtt{max}\,(0, R_\mathrm{n}(t))$ and $R_\mathrm{o}(t)$, otherwise $R_\mathrm{i}(t) = 0$;
5. compute $R_\mathrm{o}(t+\Delta t)$ and $V_\mathrm{n}(t+\Delta t)$ using Equations (S.22.1) and (S.22.2), respectively;
6. compute $R_\mathrm{n}(t+\Delta t)$ using Equation (S.22.10).

As $\hat{\lambda}(t)$ grows exponentially, in some parameter regimes MATLAB's `ode15s` function fails to evaluate Equation (S.22.2) within the time frame of the experiments. To avoid these numerical issues we evaluate $\hat{\lambda}(t)$ using Equation (S.22.9) and impose a threshold: if $\hat{\lambda}(t) > 2 \times 10^7$, then $\hat{\lambda}(t) = 2 \times 10^7$.

#### C.3 Mathematical model to interpret re-oxygenation experiments

##### C.3.1 Model derivation

The mathematical modelling approach to interpret re-oxygenation experiments, for spheroids where spherical symmetry is maintained, is similar to the mathematical modelling approach to interpret the deoxygenation experiments. Here, we present full details.

In the re-oxygenation experiments we set $p_\infty = 2$ [%] for $0 < t < t_\mathrm{s}$ [days] and $p_\infty = 21$ [%] for $t_\mathrm{s} < t < 8$ [days]. To interpret these re-oxygenation experiments we extend Greenspan's mathematical model [5]. Consistent with assumptions for deoxygenation experiments, we assume that the change in $p_\infty$ at $t_\mathrm{s}$ is instantaneous. Then we estimate the oxygen partial within the spheroid at $t_\mathrm{s}$ under normoxic and hypoxic conditions. Immediately after $t_\mathrm{s}$ the predicted necrotic radii, denoted $R_\mathrm{n}^+(t)$, is implicitly defined through $p(R_\mathrm{n}^+(t)) = 0$, is smaller than the actual necrotic radii, $R_\mathrm{n}(t)$, specifically $R_\mathrm{n}^+(t) < R_\mathrm{n}(t)$ (Figure 5e,f).

Before considering the region $R_\mathrm{n}(t) < r < R_\mathrm{n}^+(t)$, recall that parameter estimates from spheroids grown in normoxia and hypoxia differ. Specifically, $\alpha$ (Figure 3d), $\lambda = \gamma s$ (Figure 3l,n), $s$ (Figure 3l), and $\mathcal{R}$ (Figure S23) are all different. Therefore, we expect that these parameter values will evolve in time after $t_\mathrm{s}$. To account for such changes we define the following,

$$\alpha(t) = \begin{cases} \alpha_h, & t < t_\mathrm{s}, \\ \alpha_n + (\alpha_h - \alpha_n)\exp\left(-\frac{1}{\tau_\alpha}(t - t_\mathrm{s})\right), & t > t_\mathrm{s}, \end{cases} \tag{S.23}$$

$$\lambda(t) = \begin{cases} \lambda_h, & t < t_\mathrm{s}, \\ \lambda_n + (\lambda_h - \lambda_n)\exp\left(-\frac{1}{\tau_\lambda}(t - t_\mathrm{s})\right), & t > t_\mathrm{s}, \end{cases} \tag{S.24}$$

$$s(t) = \begin{cases} s_h, & t < t_\mathrm{s}, \\ s_n + (s_h - s_n)\exp\left(-\frac{1}{\tau_s}(t - t_\mathrm{s})\right), & t > t_\mathrm{s}, \end{cases} \tag{S.25}$$

$$\mathcal{R}(t) = \begin{cases} \mathcal{R}_h, & t < t_\mathrm{s}, \\ \mathcal{R}_n + (\mathcal{R}_h - \mathcal{R}_n)\exp\left(-\frac{1}{\tau_\mathcal{R}}(t - t_\mathrm{s})\right), & t > t_\mathrm{s}, \end{cases} \tag{S.26}$$

where $\tau_\alpha$, $\tau_\lambda$, $\tau_s$, and $\tau_\mathcal{R}$ denote timescales of adaptation for $\alpha$, $\lambda$, $s$, and $\mathcal{R}$ respectively. Further, the new constants in Equation (S.23) with subscripts $n$ and $h$, for example $\alpha_h$ and $\alpha_n$ represent parameter estimates from spheroids grown in normoxia and hypoxia, respectively. The other parameters parameters ($k$, $\Omega$, $\kappa$) are assumed to be constants. Hence, $R_\mathrm{c}^2(t) = 6kp_\infty/(\alpha(t)\Omega)$ [µm], $Q(t)^2 = \mathcal{R}^2(t)R_\mathrm{c}^2(t)$ [-], and $\gamma(t)$ [-] are functions of time.

In the region $R_\mathrm{n}(t) < r < R_\mathrm{n}^+(t)$ we assume that the necrotic core volume decreases a rate $\tilde{\lambda}(t) = \tilde{\lambda}\exp((t - t_\mathrm{s})/\tau_{\tilde{\lambda}}) > 0$ [day$^{-1}$]. We allow for the possibility that a fraction, $0 \leq \nu \leq 1$, of the volume lost from the necrotic core is due to cells recovering from the harsh oxygen conditions to increase the population of living cells. Note $\tilde{\lambda}$, $\tau_{\tilde{\lambda}}$, and $\nu$ are new parameters. To define $R_\mathrm{n}(t)$ we consider conservation of volume,

$$D = B_\text{n} - C_\text{n} - F_\text{n}, \tag{S.27}$$

where $D$ is the total volume of the necrotic core at time $t$ from Equation (S.2.4); $B_\text{n}$ is the volume of the necrotic core at time $t_\text{s}$; $C_\text{n}$ is the total volume of necrotic debris in the region $R_\text{n}^+(t) < r < R_\text{n}(t)$ that is lost in $t \geq t_\text{s}$; and $F_\text{n}$ is the total volume lost in the necrotic core in $t \geq t_s$ in the region $0 < r < R_\text{n}^+(t)$ when the necrotic region exists and $p(R_\text{n}^+(t)) = 0$. Writing $B_\text{n}$, $C_\text{n}$, and $F_\text{n}$ in their mathematical forms gives,

$$B_\text{n} = \frac{4\pi}{3} R_\text{n}^3(t_\text{s}), \tag{S.28.1}$$

$$C_\text{n} = 4\pi \int_{t_\text{s}}^{t} \int_{R_\text{n}^+(t)}^{R_\text{n}(t)} 3\tilde{\lambda}(t) r^2 \text{ d}r \text{ d}t, \tag{S.28.2}$$

$$F_\text{n} = \frac{4\pi}{3} \int_{t_\text{s}}^{t} 3\lambda(t) R_\text{n}^+(t_\text{s})^3 \text{ d}t. \tag{S.28.3}$$

where the three inside the integrals of Equations (S.28.2) and (S.28.3) is included for convenience.

Substituting Equations (S.28) and (S.2.4) into Equation (S.27) gives,

$$\frac{4\pi}{3} R_\text{n}^3(t) = \frac{4\pi}{3} R_\text{n}^3(t_\text{s}) - 4\pi \int_{t_\text{s}}^{t} \int_{R_\text{n}^+(t)}^{R_\text{n}(t)} 3\tilde{\lambda}(t) r^2 \text{ d}r \text{ d}t - \frac{4\pi}{3} \int_{t_s}^{t} 3\lambda(t) R_\text{n}^+(t)^3 \text{ d}t, \tag{S.29}$$

Differentiating Equation (S.29) with respect to time and simplifying gives,

$$R_\text{n}^2(t) \frac{\text{d}R_\text{n}(t)}{\text{d}t} = -\tilde{\lambda}(t) \left[ R_\text{n}^3(t) - R_\text{n}^+(t)^3 \right] - \lambda(t) R_\text{n}^+(t)^3. \tag{S.30}$$

However, Equation (S.30) is only valid for $R_\text{n}(t) > 0$. For a general equation, valid for $R_\text{n}(t) \geq 0$, we rexpress Equation (S.30) in terms of the necrotic core volume at time $t$, $V_\text{n}(t)$,

$$\frac{\text{d}V_\text{n}(t)}{\text{d}t} = -3\tilde{\lambda}(t) \left[ V_\text{n}(t) - \frac{4\pi}{3} R_\text{n}^+(t)^3 \right] - 3\lambda(t) \frac{4\pi}{3} R_\text{n}^+(t)^3. \tag{S.31}$$

Then we can define the necrotic radius, $R_\text{n}(t)$, as

$$R_\text{n}(t) = \left[ \frac{3}{4\pi} V_\text{n}(t) \right]^{\frac{1}{3}}. \tag{S.32}$$

Note that Equation (S.31) is similar to the analogous deoxygenation Equation (S.20), with the differences being $\tilde{\lambda}(t)$ instead of $\hat{\lambda}(t)$, and the last term on the right hand side is in terms of $R_\text{n}^+(t)$ instead of $R_\text{n}(t)$. We solve Equations (S.20) and (S.32) numerically using MATLAB's `ode15s` function.

As we assume that a fraction, $0 \leq \nu \leq 1$, of the matter lost from the necrotic core may become living cells we reconsider conservation of volume for living cells during this adaptation period.

Conservation of volume gives

$$A = B_\mathrm{l} + C_\mathrm{l} - D - F_\mathrm{n} + \nu C_\mathrm{n}, \tag{S.33}$$

where $A$ is the total volume of living cells at time $t$; $B_\mathrm{l}$ is the volume of living cells at time $t_\mathrm{s}$; $C_\mathrm{l}$ is the total volume of cells produced in $t \geq t_\mathrm{s}$ due to cell proliferation in $R_\mathrm{i}(t) < r < R_\mathrm{o}(t)$; $D$ is the total volume of necrotic debris at time $t$; $F_\mathrm{n}$ is the total volume lost in the necrotic core in $t \geq t_s$ in the region $0 < r < R_\mathrm{n}^+(t)$ when the necrotic region exists and $p(r(t)) < p_\mathrm{n}$ holds; and, $\nu C_\mathrm{n}$ is the total volume of necrotic debris in the region $R_\mathrm{n}^+(t) < r < R_\mathrm{n}(t)$ that recovers due to re-oxygenation to be classified as living cells in $t \geq t_\mathrm{s}$. Note that:

- $A$ and $D$ are the same as $A$ and $D$ defined in Equations (S.2.1) and (S.2.4), respectively,
- $B_\mathrm{l}$ and $C_\mathrm{l}$ are $B$ and $C$ from Equations (S.2.2) and (S.2.3), respectively, but starting at $t = t_\mathrm{s}$,
- $F_\mathrm{n}$ is defined in Equation (S.28.3) and is analogous to the definition of $E$ in Equation (S.2.5),
- $C_\mathrm{n}$ is defined in Equation (S.28.2).

Substituting the mathematical forms of $A$, $B_\mathrm{l}$, $C_\mathrm{l}$, $D$, $F_\mathrm{n}$, and, $C_\mathrm{n}$ into the conservation of volume Equation (S.33) and simplifying gives,

$$\begin{aligned}\frac{4\pi}{3}R_\mathrm{o}^3(t) =& \frac{4\pi}{3}R_\mathrm{o}^3(t_\mathrm{s}) + 4\pi \int_{t_\mathrm{s}}^{t}\int_{R_\mathrm{i}(t)}^{R_\mathrm{o}(t)} s(t) r^2 \ \mathrm{d}r \ \mathrm{d}t \\ &- 4\pi \int_{t_\mathrm{s}}^{t}\int_{0}^{R_\mathrm{n}^+(t)} 3\lambda(t) r^2 \ \mathrm{d}r \ \mathrm{d}t + 4\pi\nu \int_{t_\mathrm{s}}^{t}\int_{R_\mathrm{n}^+(t)}^{R_\mathrm{n}(t)} 3\tilde{\lambda}(t) r^2 \ \mathrm{d}r \ \mathrm{d}t.\end{aligned} \tag{S.34}$$

Differentiating Equation (S.34) with respect to time and simplifying gives,

$$R_\mathrm{o}^2(t)\frac{\mathrm{d}R_\mathrm{o}(t)}{\mathrm{d}t} = \frac{s(t)}{3}\left[R_\mathrm{o}^3(t) - R_\mathrm{i}^3(t)\right] - \lambda(t) R_\mathrm{n}^+(t)^3 + \nu\tilde{\lambda}(t)\left[R_\mathrm{n}^3(t) - R_\mathrm{n}^+(t)^3\right]. \tag{S.35}$$

Note that Equation (1) has an additional term on the right hand side in comparison to the right hand side of Equation (S.35). Further, the second term on the right hand side of Equation (S.35) is in terms of $R_\mathrm{n}^+(t)$ instead of $R_\mathrm{n}(t)$ as in Equation (1). At late times $R_\mathrm{n}(t)$ tends to $R_\mathrm{n}^+(t)$ and we recover Equation (1) in with parameters from hypoxia. As with the deoxygenation experiments, the time evolution of the inhibited region, $R_\mathrm{i}(t)$, in the re-oxygenation experiments is assumed to be governed by the waste mechanisms.

##### Complete re-oxygenation mathematical model

Here, we present the full governing system of equations for $0 < t < t_\mathrm{s}$ and $t > t_\mathrm{s}$.

For $0 < t < t_\mathrm{s}$ we solve Greenspan's mathematical model [5] in normoxia, using hypothesis 2, where $R_\mathrm{o}(t)$, $R_\mathrm{n}(t)$, and $R_\mathrm{i}(t)$ are determined from the differential-algebraic system of Equations (6.1) - (6.3).

For $t > t_\mathrm{s}$ when the spheroid adapts to normoxic conditions, we solve,

$$R_\mathrm{o}^2(t)\frac{\mathrm{d}R_\mathrm{o}(t)}{\mathrm{d}t} = \frac{s(t)}{3}\left[R_\mathrm{o}^3(t) - \max\left(R_\mathrm{i}^3(t), R_\mathrm{n}^3(t)\right)\right] + \nu\tilde{\lambda}(t)\left[R_\mathrm{n}^3(t) - R_\mathrm{n}^+(t)^3\right], \quad \text{(S.36.1)}$$

$$\frac{\mathrm{d}V_\mathrm{n}(t)}{\mathrm{d}t} = -3\tilde{\lambda}(t)\left[V_\mathrm{n}(t) - \frac{4\pi}{3}R_\mathrm{n}^+(t)^3\right] - 3\lambda(t)\frac{4\pi}{3}R_\mathrm{n}^+(t)^3, \quad \text{(S.36.2)}$$

$$R_\mathrm{c}^2(t) = R_\mathrm{o}^2(t) - R_\mathrm{n}^+(t)^2 - \frac{2R_\mathrm{n}^+(t)^2}{R_\mathrm{o}(t)}\left(R_\mathrm{o}(t) - R_\mathrm{n}^+(t)\right), \quad \text{(S.36.3)}$$

$$\mathcal{R}^2(t) = R_\mathrm{o}^2(t) - R_\mathrm{i}^2(t) - 2R_\mathrm{n}^3(t)\left(\frac{1}{R_\mathrm{i}(t)} - \frac{1}{R_\mathrm{o}(t)}\right), \quad \text{(S.36.4)}$$

$$\alpha(t) = \alpha_n + (\alpha_h - \alpha_n)\exp\left(-\frac{1}{\tau_\alpha}(t - t_\mathrm{s})\right), \quad \text{(S.36.5)}$$

$$\lambda(t) = \lambda_n + (\lambda_h - \lambda_n)\exp\left(-\frac{1}{\tau_\lambda}(t - t_\mathrm{s})\right), \quad \text{(S.36.6)}$$

$$s(t) = s_n + (s_h - s_n)\exp\left(-\frac{1}{\tau_s}(t - t_\mathrm{s})\right), \quad \text{(S.36.7)}$$

$$\mathcal{R}(t) = \mathcal{R}_n + (\mathcal{R}_h - \mathcal{R}_n)\exp\left(-\frac{1}{\tau_\mathcal{R}}(t - t_\mathrm{s})\right), \quad \text{(S.36.8)}$$

$$\tilde{\lambda}(t) = \tilde{\lambda}\exp\left(\frac{1}{\tau_{\tilde{\lambda}}}(t - t_\mathrm{s})\right), \quad \text{(S.36.9)}$$

$$R_\mathrm{n}(t) = \left[\frac{3}{4\pi}V_\mathrm{n}(t)\right]^{\frac{1}{3}}, \quad \text{(S.36.10)}$$

$$R_\mathrm{c}^2(t) = \frac{6kp_\infty}{\alpha(t)\Omega}. \quad \text{(S.36.11)}$$

Note that in the long time limit, $t \to \infty$, we recover Greenspan's mathematical model for hypoxia (Equations (6.1)-(6.3)) from Equations (S.22). Specifically, $\alpha(t) \to \alpha_n$, $\lambda(t) \to \lambda_n$, $s(t) \to s_n$ and $\mathcal{R}(t) \to \mathcal{R}_n$ as $t \to \infty$. Further, the term involving $\tilde{\lambda}(t)$ dominates the right hand side of Equation (8.10) as $t \to \infty$, so $R_\mathrm{n}(t) \to R_\mathrm{n}^+(t)$ as $t \to \infty$.

##### C.3.2 Numerical methods

The code to simulate the re-oxygenation model is available on the GitHub repository listed in Methods: Code Availability. For $0 < t < t_\mathrm{s}$ we numerically solve Greenspan's model as described in Section C.1.2. Here, we outline the key details to solve Equations (S.36.1)-(S.36.11) for $t \geq t_\mathrm{s}$.

For $t \geq t_\mathrm{s}$ note that: Equations (S.36.1) and (S.36.2) are ordinary differential equations; Equations (S.36.3), (S.36.4), and (S.36.10)-(S.36.11) are algebraic equations; Equations (S.36.5)-(S.36.8) include exponential decay; and Equation (S.36.9) describes exponential growth.

To numerically solve the coupled differential-algebraic system of Equations (S.36.1)-(S.36.11) we use MATLAB's `ode15s` function. At time $t$ we assume that $R_{\mathrm{o}}(t)$ and $R_{\mathrm{n}}(t)$ are known. To determine $R_{\mathrm{i}}(t)$, $R_{\mathrm{n}}(t+\Delta t)$, and $R_{\mathrm{o}}(t+\Delta t)$, where the time step $\Delta t$ is determined by the `ode15s` function, we,

1. evaluate Equations (S.36.5)-(S.36.9) to obtain $\alpha(t)$, $\lambda(t)$, $s(t)$, $\mathcal{R}(t)$, and $\tilde{\lambda}(t)$;
2. evaluate $R_{\mathrm{c}}^2(t)$ using Equation (S.36.11);
3. solve Equation (S.36.10) using MATLAB's `roots` function and define $R_{\mathrm{n}}^{+}(t)$ as the root between 0 and $R_{\mathrm{o}}(t)$, otherwise $R_{\mathrm{n}}^{+}(t)=0$;
4. solve Equation (S.36.4) using MATLAB's `roots` function and define $R_{\mathrm{i}}(t)$ as the root between $\mathtt{max}\,(0, R_{\mathrm{n}}(t))$ and $R_{\mathrm{o}}(t)$, otherwise $R_{\mathrm{i}}(t)=0$;
5. compute $R_{\mathrm{o}}(t+\Delta t)$ and $V_{\mathrm{n}}(t+\Delta t)$ using Equations (S.36.1) and (S.36.2), respectively;
6. compute $R_{\mathrm{n}}(t+\Delta t)$ using Equation (S.36.10).

As $\tilde{\lambda}(t)$ grows exponentially, in some parameter regimes MATLAB's `ode15s` function fails to evaluate Equation (S.36.2) within the time frame of the experiments. To avoid these numerical issues we evaluate $\tilde{\lambda}(t)$ using Equation (S.36.9) and impose a threshold: if $\tilde{\lambda}(t) > 2\times 10^7$, then $\tilde{\lambda}(t) = 2\times 10^7$.

### D Additional results for WM983b spheroids

#### D.1 Oxygen diffusion alone is insufficient to describe spheroid growth

In Figure S21 we show additional measurements of $\xi_{\mathrm{n}}(t) = R_{\mathrm{n}}(t)/R_{\mathrm{o}}(t)$, $\xi_{\mathrm{i}}(t) = R_{\mathrm{i}}(t)/R_{\mathrm{o}}(t)$, and $\xi_{\mathrm{p}}(t) = R_{\mathrm{p}}(t)/R_{\mathrm{o}}(t)$ for spheroids grown in normoxia and hypoxia. These results support and are consistent with the results and discussion in the main manuscript about Figure 3.

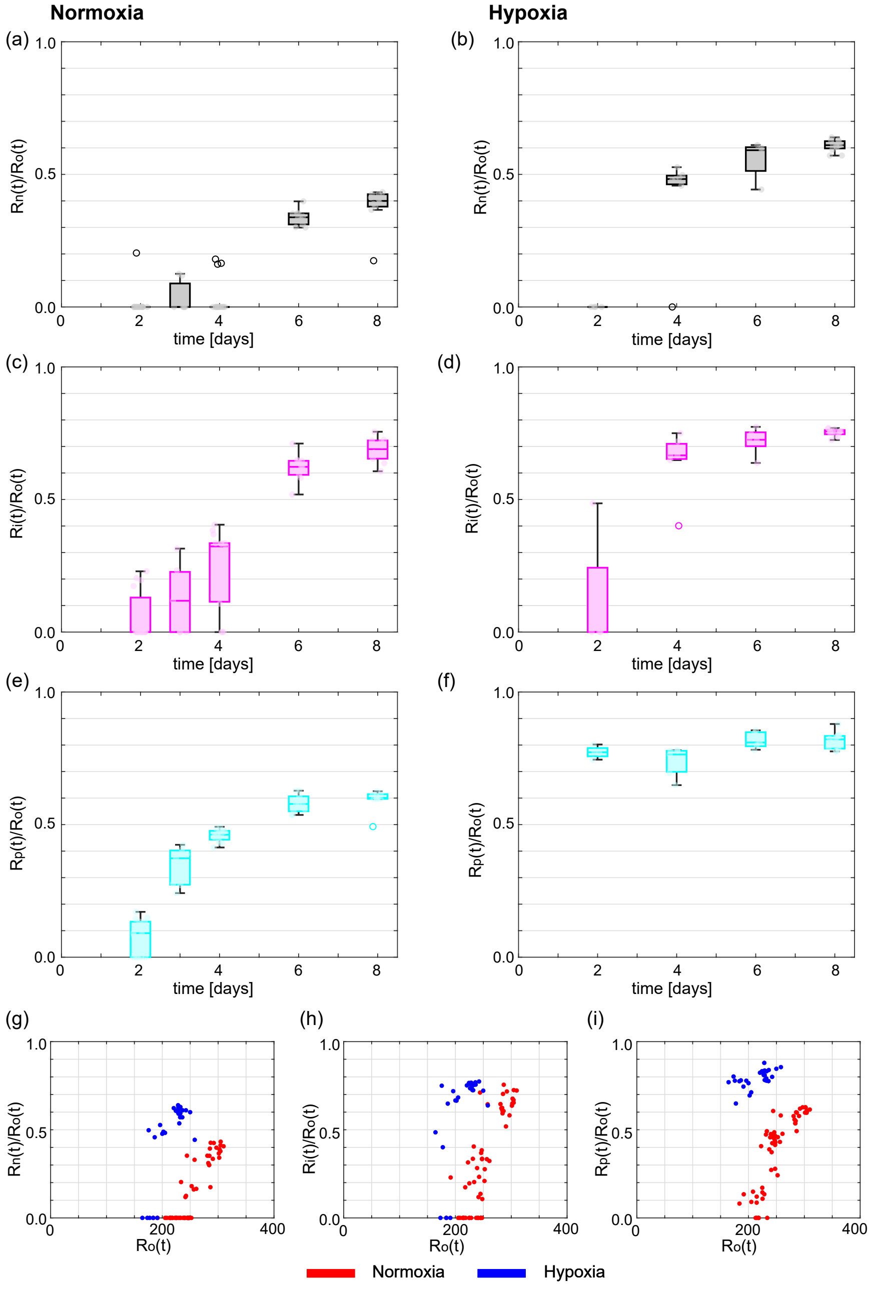


Figure S21: Additional experimental measurements comparing spheroids grown in normoxia and hypoxia. (a-b) Estimates of $\xi_{\mathrm{n}}(t) = R_{\mathrm{n}}(t)/R_{\mathrm{o}}(t)$. (a-b) Estimates of $\xi_{\mathrm{n}}(t) = R_{\mathrm{n}}(t)/R_{\mathrm{o}}(t)$ with time for (a) normoxia and (b) hypoxia. (c-d) Estimates of $\xi_{\mathrm{i}}(t) = R_{\mathrm{i}}(t)/R_{\mathrm{o}}(t)$ with time for (c) normoxia and (d) hypoxia. (e-f) Estimates of $\xi_{\mathrm{p}}(t) = R_{\mathrm{p}}(t)/R_{\mathrm{o}}(t)$ with time for (e) hypoxia and (f) hypoxia. (g-h) Estimates of (g) $\xi_{\mathrm{n}}(t)$, (h) $\xi_{\mathrm{i}}(t)$, and (i) $\xi_{\mathrm{p}}(t)$ with against outer radius, $R_{\mathrm{o}}(t)$.

##### D.1.1 Analysing spheroid snapshots independently to explore oxygen assumptions

In Figure S22 we present additional results analysing spheroid images independently to explore oxygen assumptions. Using Equation (S.9) we estimate the outer radius when the necrotic region forms, $R_c$ [µm]. Then we estimate the constant rate of volume of oxygen gas per unit tumour mass that is consumed by living cells, $\alpha$ [$m^3$ $kg^{-1}$ $s^{-1}$], by rearranging Equation (S.8) to solve for $\alpha$. Rearranging Equations (S.10) and (S.11) we estimate the oxygen threshold that defines the inhibited region according to hypothesis 1, $p_i$ [%]. These results support and are consistent with the results and discussion in the main manuscript about Figure 3.

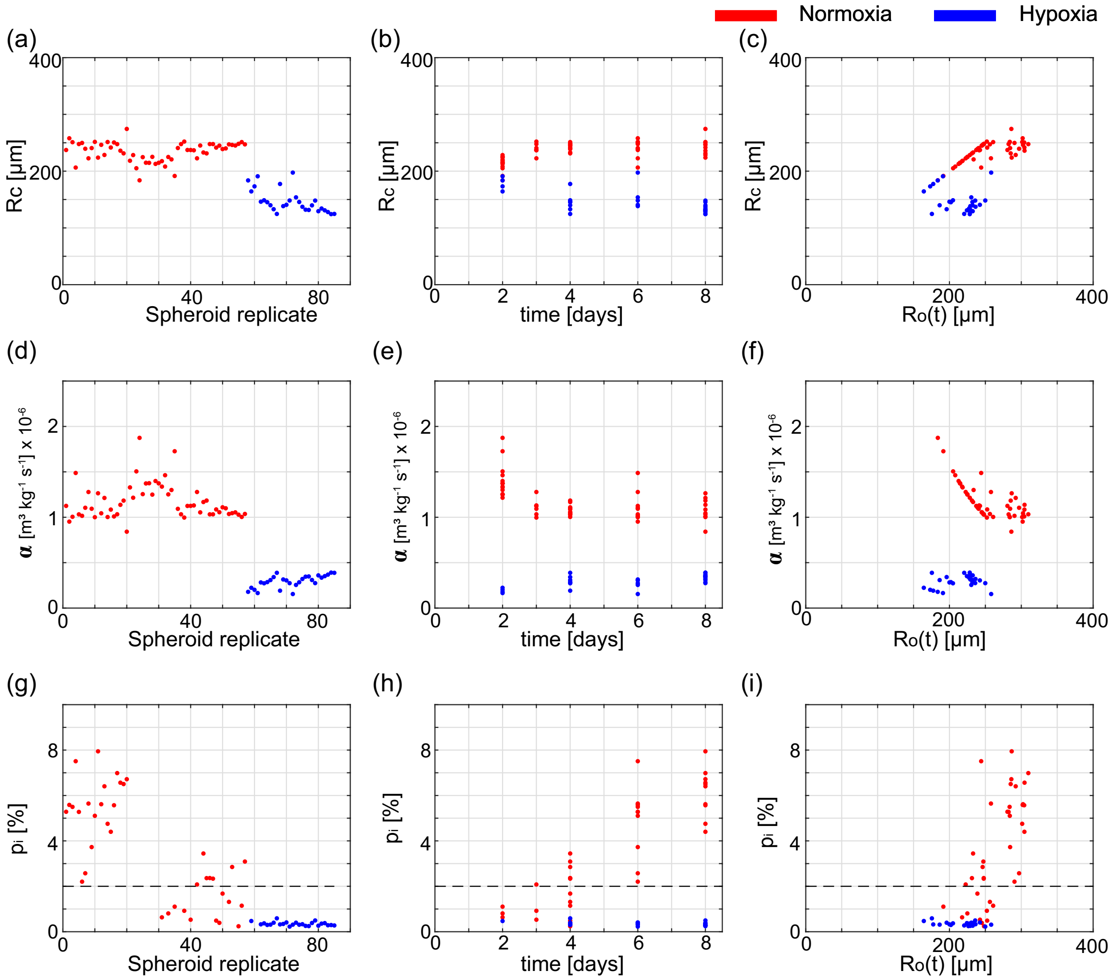


Figure S22: Oxygen diffusion and consumption is sufficient to describe formation and growth of necrotic core but not sufficient to describe formation and growth of inhibited region. Estimates of $R_c$ (a) per spheroid; (b) with time; (c) against $R_o(t)$. Estimates of $\alpha$ (d) per spheroid; (e) with time; (f) against $R_o(t)$. Estimates of $p_i$ (g) per spheroid; (h) with time; (i) against $R_o(t)$. In (a-i) each data point represents a single spheroid. Data points are only included for $p_i$ if the spheroid is in phase (ii) or phase (iii), consistent with when equations used to estimate $p_i$ are valid.

##### D.1.2 Analysing spheroid snapshots independently to explore waste assumptions

In Figure S23 we present additional results analysing spheroid images independently to estimate the outer radius when the inhibited region first forms, $\mathcal{R}$ [µm], using Equation (S.13). These results support and are consistent with the results and discussion in the main manuscript about Figure 3.

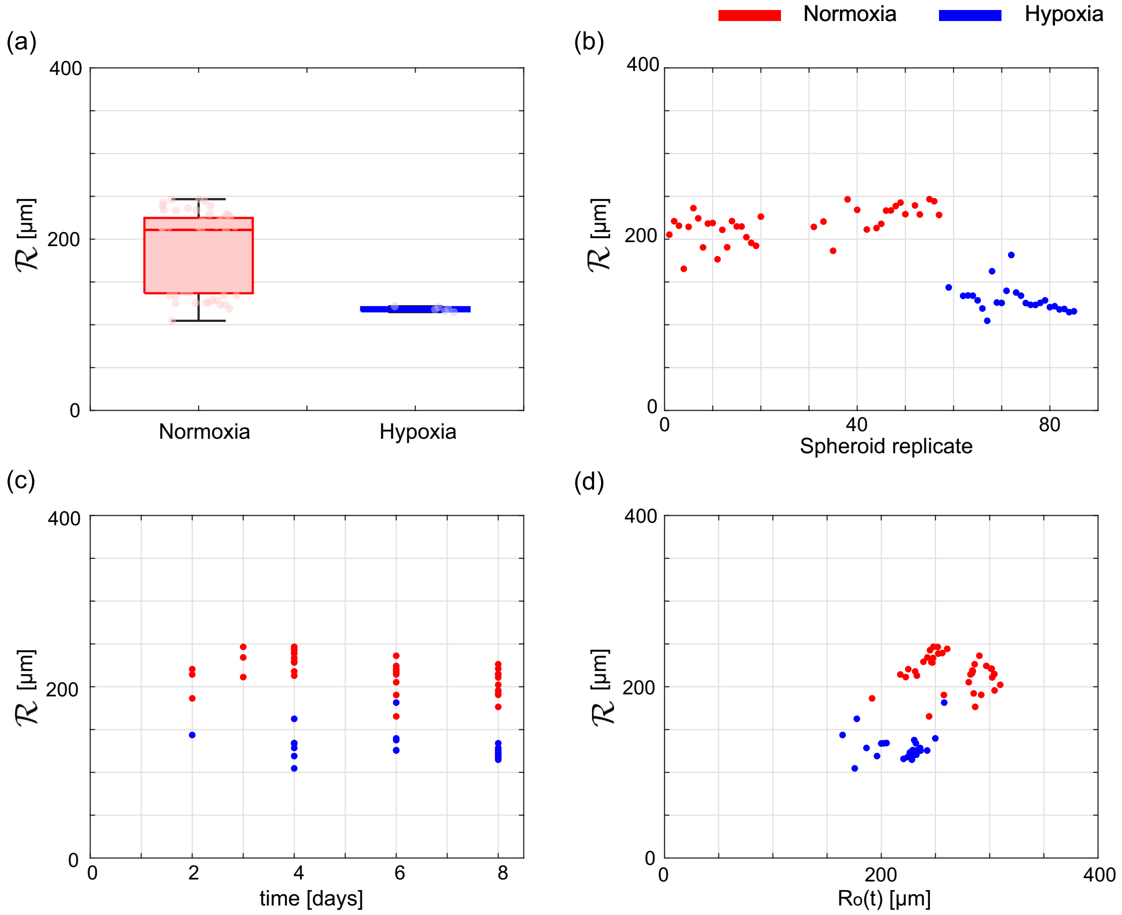


Figure S23: Analysing whether production and diffusion of waste describes formation and growth of inhibited region. Estimates of the outer radius when the inhibited region first forms, $\mathcal{R} = (\beta_{\mathrm{i}}\kappa/P)^{1/2}$ (a) box chart; (b) per spheroid; (c) with time; (d) against $R_{\mathrm{o}}(t)$. In (b-d) each data point represents a single spheroid. Data points are only included only if the spheroid is in phase (ii) or phase (iii), consistent with when the equations used to estimate $\mathcal{R}$ are valid.

##### D.1.3 Parameter estimation

In Figure S24 we show univariate posterior densities from Bayesian inference to estimate parameters of Greenspan's mathematical model for spheroids grown in normoxia and hypoxia oxygen conditions. Posterior densities for Greenspan's model parameters in Figure S24a,b,c are consistent with parameter estimates from profile likelihood analysis in Figure 3k,m,l, respectively. Note that here we show $\mathcal{R}$ (Figure S24d) instead of $Q$ and $\lambda$ (Figure S24e) instead of $\gamma$. Prediction intervals in Figure S24f-g accurately capture the experimental data suggesting that the parameter estimates are reasonable.

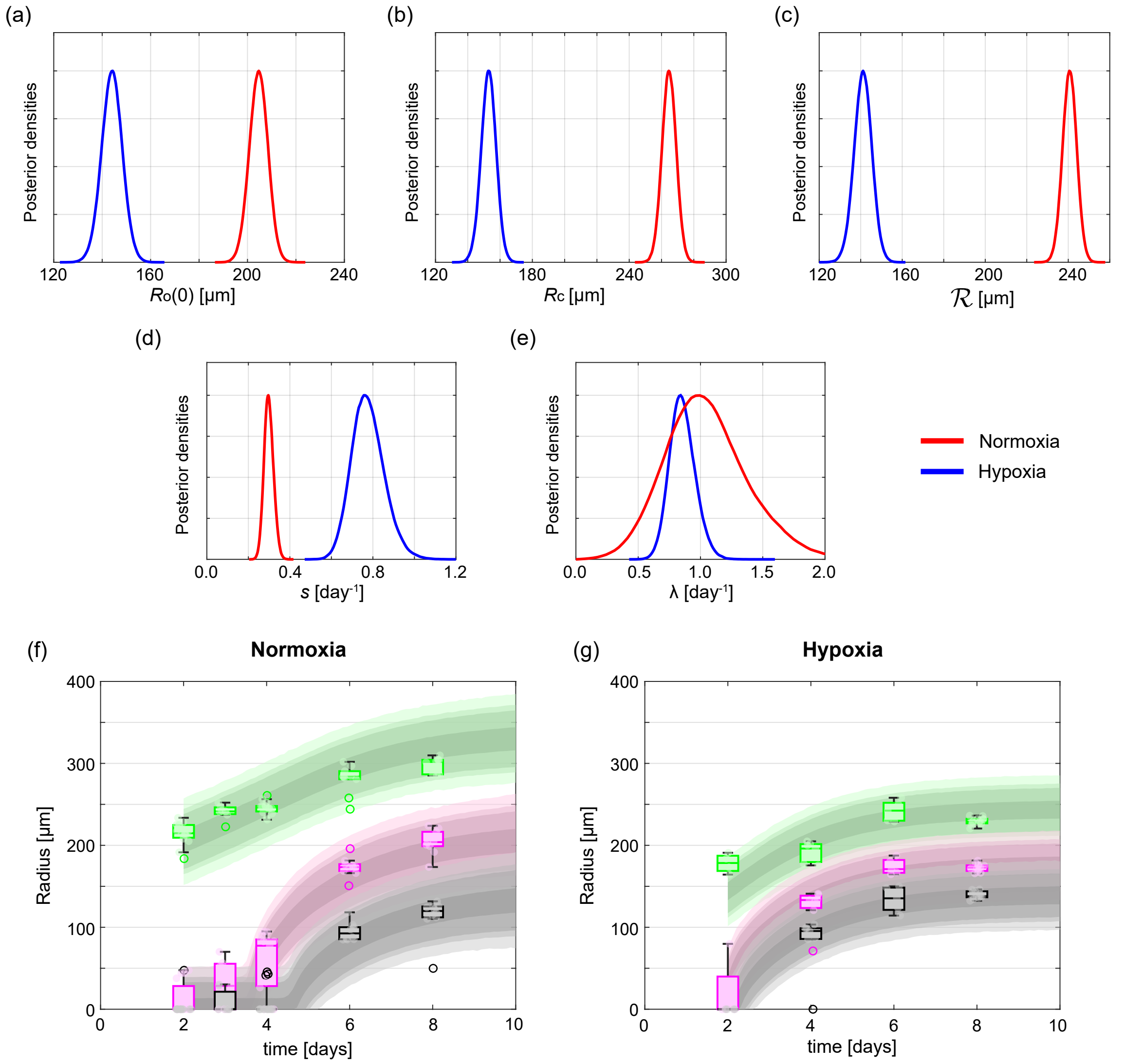


Figure S24: Bayesian inference to estimate parameters of Greenspan's mathematical model for spheroids grown in normoxia and hypoxia. (a-e) Posterior densities for Greenspan model parameters: (a) $R_o(0)$, (b) $R_c$, (c) $\mathcal{R}$, (d) $s$, (e) $\lambda$. Prediction intervals for (f) normoxia. (g) hypoxia. In (f-g) colour bands, in decreasing darkness, represent 50%, 75%, 95%, 97.5%, and 99.5% prediction intervals.

#### D.2 Deoxygenation

Here we present additional results analysing deoxygenation experiments. These results support and agree with comments in the main manuscript on Figure 4.

In Figure S25a-c, measurements of $\xi_{\mathrm{n}}(t) = R_{\mathrm{n}}(t)/R_{\mathrm{o}}(t)$, $\xi_{\mathrm{i}}(t) = R_{\mathrm{i}}(t)/R_{\mathrm{o}}(t)$, and $\xi_{\mathrm{p}}(t) = R_{\mathrm{p}}(t)/R_{\mathrm{o}}(t)$ suggest that deoxygenated spheroids approach spheroid structures observed in spheroids grown in hypoxia (Figure S21b,d,f).

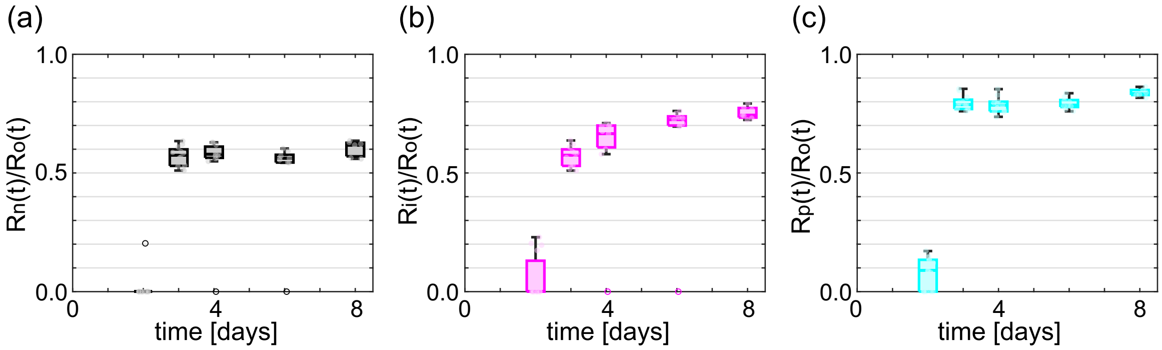


Figure S25: Additional results from deoxygenation experiments for WM983b spheroids. Measurements of (a) $\xi_{\mathrm{n}}(t) = R_{\mathrm{n}}(t)/R_{\mathrm{o}}(t)$, (b) $\xi_{\mathrm{i}}(t) = R_{\mathrm{i}}(t)/R_{\mathrm{o}}(t)$, and (c) $\xi_{\mathrm{p}}(t) = R_{\mathrm{p}}(t)/R_{\mathrm{o}}(t)$.

##### D.2.1 Parameter estimation

Prediction intervals for the deoxygenation model are shown in Figure S26a. Results suggest the model and parameter estimates accurately capture the experimental data. Further, we observe rapid growth of the necrotic region at early times consistent with the experimental data.

One advantage of using a mathematical model to analyse the experimental data is that we can explore what would happen if we had additional data. To investigate the predicted rapid growth of the necrotic region at early times, we suppose we have additional data at $t = 2.5$ [days]. To generate these additional synthetic data points we first compute the mean of each measurement type at $t = 2$ [days] and at $t = 3$ [days], which we denote $\bar{R}_{\mathrm{o}}(2)$, $\bar{R}_{\mathrm{n}}(2)$, $\bar{R}_{\mathrm{i}}(2)$, $\bar{R}_{\mathrm{o}}(3)$, $\bar{R}_{\mathrm{n}}(3)$, and $\bar{R}_{\mathrm{i}}(3)$. Then we average for each measurement type and add noise. For example, to generate eight synthetic measurements of $R_{\mathrm{o}}(2.5)$ we generate eight samples from a normal distribution with mean $(\bar{R}_{\mathrm{o}}(2) + \bar{R}_{\mathrm{o}}(3))/2$ and standard deviation set to the pooled standard deviation all measurements of $R_{\mathrm{o}}(t)$. Similarly, for $R_{\mathrm{n}}(2.5)$ and $R_{\mathrm{i}}(2.5)$. Prediction intervals in Figure S26b show that we still accurately capture the experimental data and with slower predicted growth of the necrotic core at early times. These results suggest that additional data at early times would be beneficial to understand the early time dynamics.

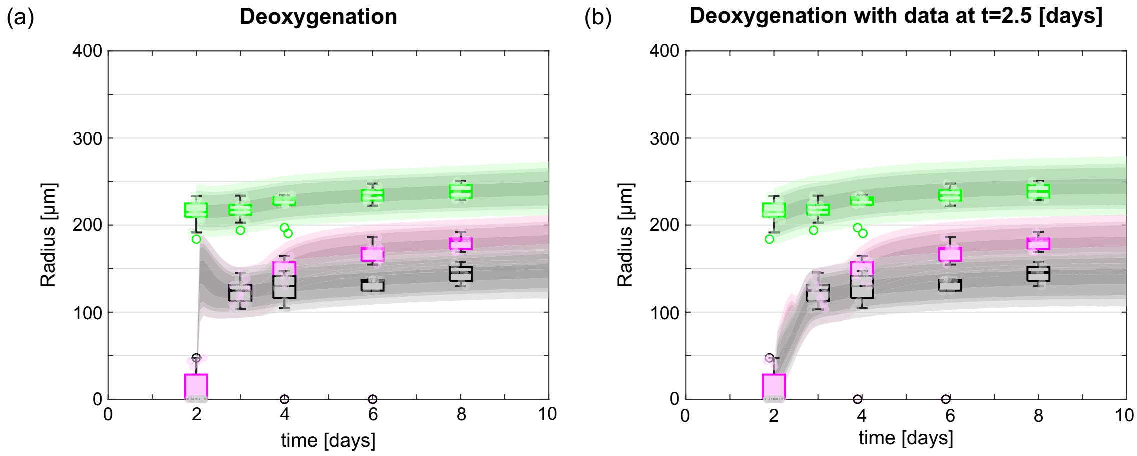


Figure S26: Prediction intervals for (a) deoxygenation experimental data and (b) deoxygenation experimental data with additional synthetic data points at $t = 2.5$ [days]. In (a-b) colour bands, in decreasing darkness, represent 50%, 75%, 95%, 97.5%, and 99.5% prediction intervals. Additional synthetic data at $t = 2.5$ not shown.

#### D.3 Re-oxygenation

Here we present additional results and discussion corresponding to the WM983b re-oxygenation experiments presented in Figure 5.

##### D.3.1 Necrotic core movement in WM983b spheroids

Direction of movement of necrotic core to edge of spheroid appears random in re-oxygenation experiments with WM983b spheroids. In Figure S27b-d we track the centroid of the necrotic core relative to the spheroid centroid over 24 hours and observe that the motion appears random since there is no obvious systematic direction. Similarly for other spheroids (Figure S27e).

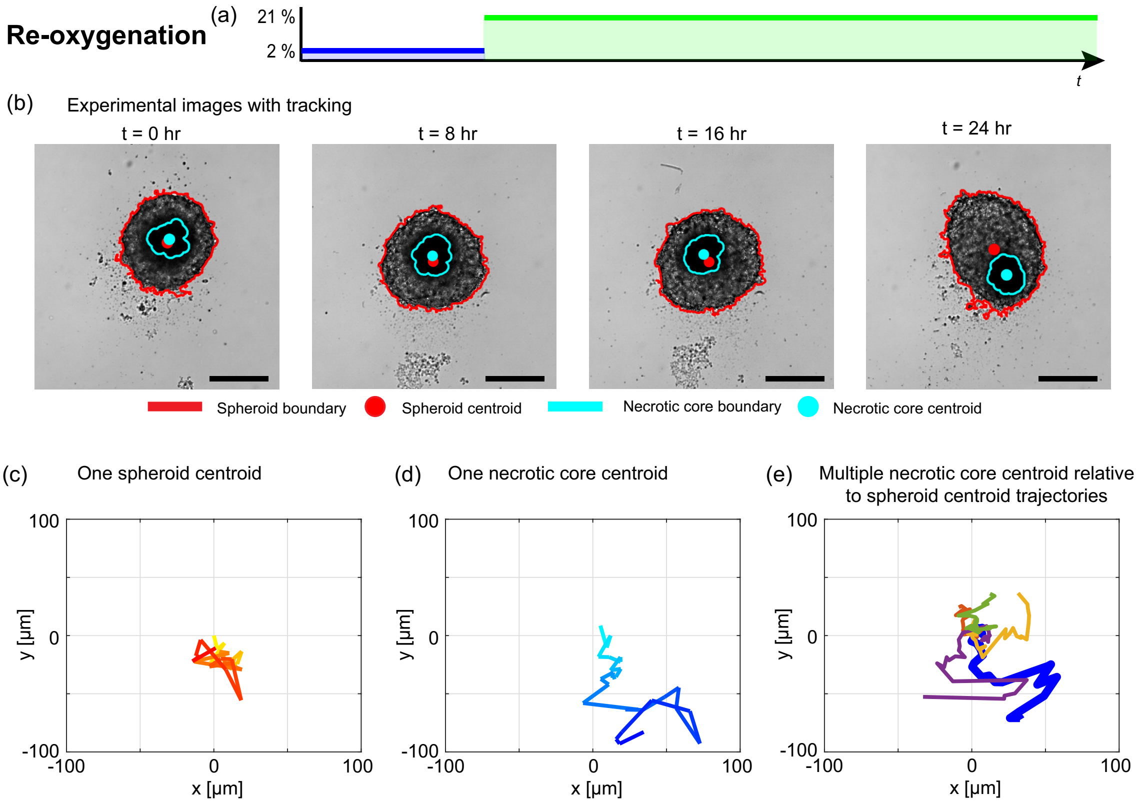


Figure S27: Direction of movement of necrotic core to edge of spheroid appears random in WM983b re-oxygenation experiments. (a) Schematic for re-oxygenation experiment, with $t_s = 2.5$ [days]. (b) Exemplar experimental brightfield images of a single spheroid with image processing to detect spheroid boundary and centroid (red) and necrotic core boundary and centroid (cyan). (c) Trajectory of centroid of spheroid for spheroid imaged in (b). (d) Trajectory of centroid of necrotic core for spheroid imaged in (b). (e) Five exemplar trajectories of the centroid of the necrotic core relative to the centroid of the spheroid. In (e) the thick blue trajectory corresponds to spheroid imaged in (b).

### E Additional results for WM793b cell line

Here, we present results for WM793b normoxia, hypoxia, and deoxygenation experiments. We also include additional results to supplement re-oxygenation results shown in Figure 5a-h.

Results in Figures S28a-d suggest that we interpret spheroid growth using hypothesis 2, in agreement with results in the main manuscript for WM983b spheroids. In Figures S28e-i, prediction intervals show that the mathematical models accurately describe WM793b normoxia, hypoxia, deoxygenation and re-oxygenation experiments.

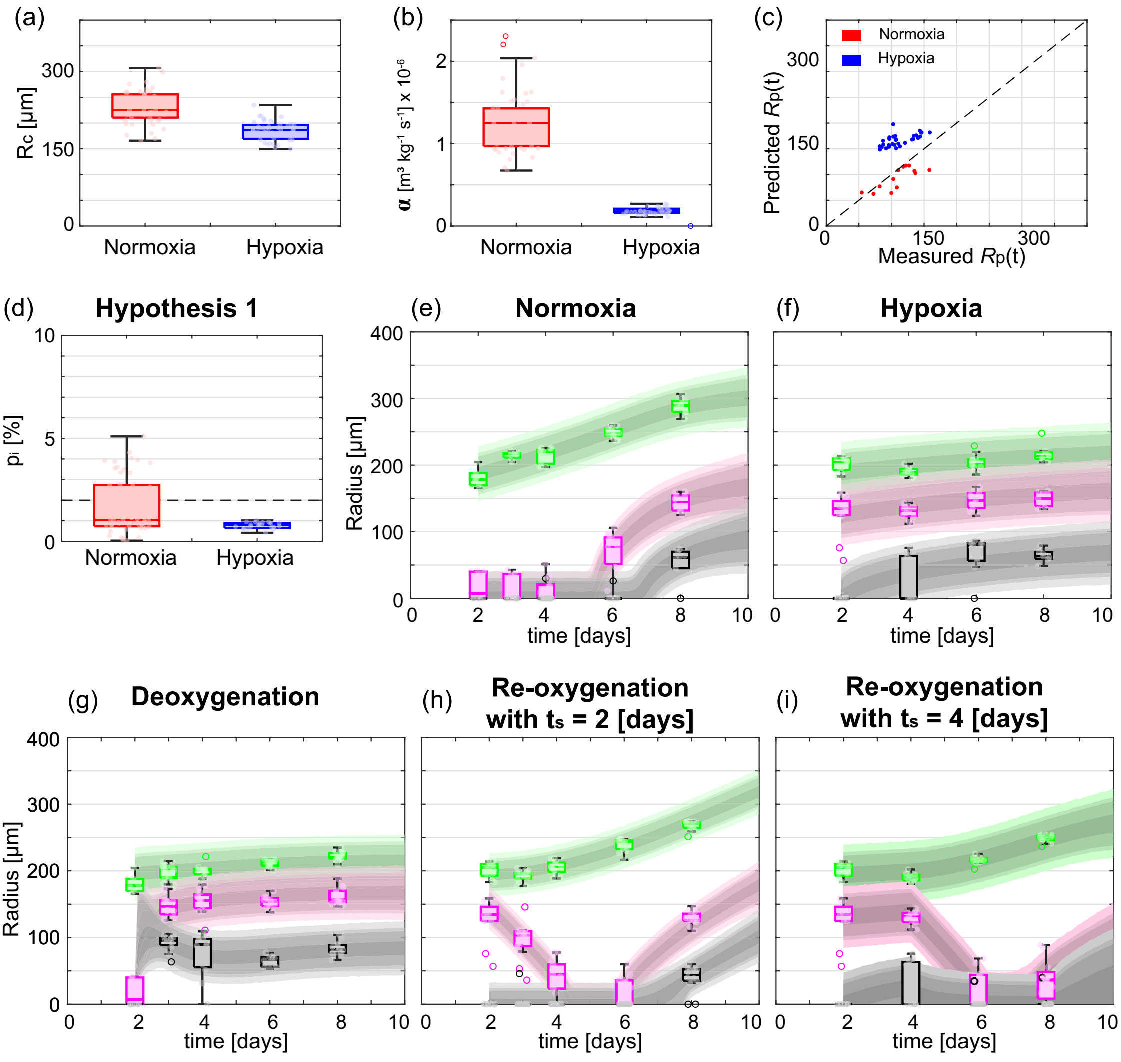


Figure S28: Additional results for WM793b spheroids. (a-d) Mechanisms governing tumour spheroid growth in normoxia and hypoxia. (a) Box chart for estimated outer radius when necrotic region forms, $R_c$ [µm]. (b) Box chart for estimated oxygen consumption rate, $\alpha$ [m$^3$ kg$^{-1}$ s$^{-1}$]. (c) Comparison of measured and predicted $R_p(t)$ when pimonidazole staining is present. Note this does not include images where pimonidazole staining is present but does not surround the necrotic core, for example Day 8 of Figure S7. (d) Box chart for estimated oxygen partial pressure defining inhibited region from hypothesis 1, $p_i$ [%]. (e-i) Experimental data and prediction intervals for (e) normoxia, (f) hypoxia, (g) deoxygenation, (h) re-oxygenation with $t_s = 2$ [days], and (i) re-oxygenation with $t_s = 4$ [days].

### F Additional results for WM164 cell line

Here, we present results for the WM164 normoxia, hypoxia, deoxygenation, and re-oxygenation experiments.

Results in Figures S29a-d suggest that we interpret spheroid growth using hypothesis 2, in agreement with results in the main manuscript for WM983b spheroids. In Figures S28e-i, prediction intervals show that the mathematical models accurately describe WM164 normoxia, hypoxia, deoxygenation and re-oxygenation experiments. Here prediction intervals are wider due to greater variability in WM164 spheroid measurements. Additional care should be exercised interpreting WM164 re-oxygenation results. Brightfield timelapse images show that mass from the necrotic core can move to the periphery and exit the spheroid (Figure S18, Movie S3).

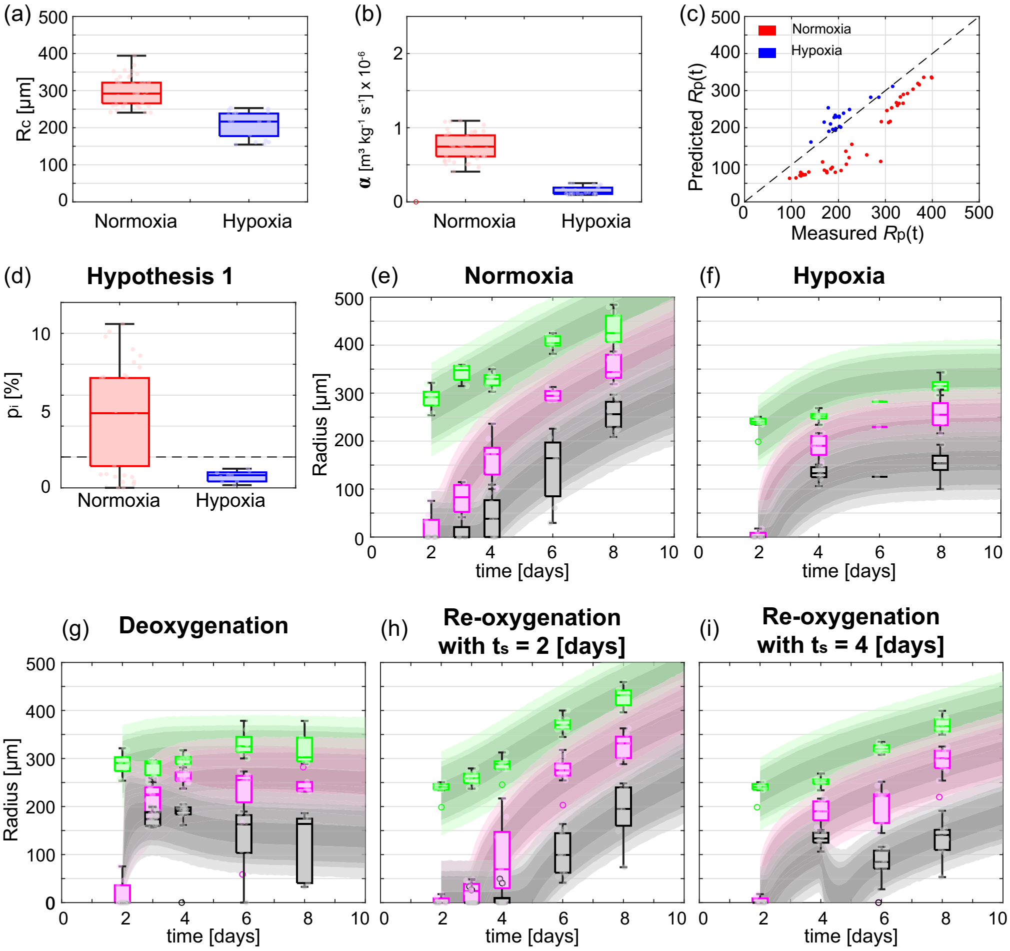


Figure S29: Additional results for WM164 spheroids. (a-d) Mechanisms governing tumour spheroid growth in normoxia and hypoxia. (a) Box chart for estimated outer radius when necrotic region forms, $R_c$ [µm]. (b) Box chart for estimated oxygen consumption rate, $\alpha$ [m$^3$ kg$^{-1}$ s$^{-1}$]. (c) Comparison of measured and predicted $R_p(t)$ when pimonidazole staining is present. (d) Box chart for estimated oxygen partial pressure defining inhibited region from hypothesis 1, $p_i$ [%]. (e-i) Experimental data and prediction intervals for (e) normoxia, (f) hypoxia, (g) deoxygenation, (h) re-oxygenation with $t_s = 2$ [days], and (i) re-oxygenation with $t_s = 4$ [days].

### G Summary statistics and MCMC diagnostics

Here, we present summary statistics of the MCMC chains and MCMC diagnostics. Results are shown for the mathematical models used to interpret normoxia (Table S2), hypoxia (Table S3), deoxygenation (Table S4), and re-oxygenation experiments (Tables S5 and S6).

To interpret normoxia and hypoxia experiments we use Greenspan's mathematical model. In each case and for every parameter $\hat{R}$ is very close to one indicating that the MCMC chains converge (Tables S2,S3). Furthermore, posterior densities are well-formed around a single central peak showing parameters are identifiable, for example Figure S24. To interpret deoxygenation and re-oxygenation experiments we increase the complexity of the mathematical model. In particular, the number of parameters increases from five parameters in Greenspan's model to fifteen and seventeen parameters in the deoxygenation and re-oxygenation mathematical models, respectively. As expected, when we increase the complexity of the model we encounter challenges of parameter identifiability and convergence of MCMC chains. While most values of $\hat{R}$ are below the convergence threshold a small number are above (Tables S4, S5, and S6). Additional radial measurements and different types of experimental measurements would be beneficial.

| Cell line | Parameter | Units | Mean | $\sigma$ | $Q_{25\%}$ | $Q_{50\%}$ | $Q_{75\%}$ | $\hat{R}$ |
|---|---|---|---|---|---|---|---|---|
| WM983b | $R_o(0)$ | µm | 204.58 | 3.83 | 202.01 | 204.61 | 207.17 | 1.0001 |
| | $R_c$ | µm | 264.69 | 4.39 | 261.74 | 264.69 | 267.66 | 1.0000 |
| | $\mathcal{R}$ | µm | 240.68 | 3.42 | 238.39 | 240.70 | 242.99 | 1.0000 |
| | $s$ | day$^{-1}$ | 0.30 | 0.02 | 0.28 | 0.30 | 0.31 | 1.0001 |
| | $\lambda$ | day$^{-1}$ | 1.05 | 0.34 | 0.82 | 1.02 | 1.25 | 1.0001 |
| WM793b | $R_o(0)$ | µm | 186.56 | 3.40 | 184.28 | 186.58 | 188.86 | 1.0000 |
| | $R_c$ | µm | 267.07 | 3.98 | 264.41 | 267.12 | 269.77 | 1.0001 |
| | $\mathcal{R}$ | µm | 242.06 | 2.92 | 240.09 | 242.05 | 244.01 | 1.0001 |
| | $s$ | day$^{-1}$ | 0.22 | 0.01 | 0.21 | 0.22 | 0.23 | 1.0000 |
| | $\lambda$ | day$^{-1}$ | 2.27 | 1.53 | 1.00 | 2.03 | 3.35 | 1.0000 |
| WM164 | $R_o(0)$ | µm | 265.60 | 7.78 | 260.40 | 265.65 | 270.84 | 1.0001 |
| | $R_c$ | µm | 326.83 | 7.23 | 322.02 | 326.92 | 331.76 | 1.0001 |
| | $\mathcal{R}$ | µm | 283.98 | 7.81 | 278.83 | 284.08 | 289.26 | 1.0001 |
| | $s$ | day$^{-1}$ | 0.37 | 0.03 | 0.35 | 0.37 | 0.39 | 1.0001 |
| | $\lambda$ | day$^{-1}$ | 0.12 | 0.09 | 0.05 | 0.10 | 0.17 | 1.0001 |

Table S2: Greenspan's model parameters and MCMC diagnostics for normoxia experiments. Summary statistics of the MCMC chains include: mean; standard deviation, $\sigma$; and, 25%, 50%, and 75% quartiles, $Q_{25\%}$, $Q_{50\%}$, and $Q_{75\%}$, respectively. To assess convergence of the MCMC chains we compute the potential scale reduction factor, $\hat{R}$, [9] where convergence corresponds to $\hat{R} < 1.1$.

| Cell line | Parameter | Units | Mean | $\sigma$ | $Q_{25\%}$ | $Q_{50\%}$ | $Q_{75\%}$ | $\hat{R}$ |
|---|---|---|---|---|---|---|---|---|
| WM983b | $R_o(0)$ | µm | 144.03 | 4.30 | 141.17 | 144.05 | 146.90 | 1.0001 |
| | $R_c$ | µm | 152.92 | 4.65 | 149.83 | 152.95 | 156.04 | 1.0000 |
| | $\mathcal{R}$ | µm | 141.09 | 4.29 | 138.25 | 141.13 | 143.97 | 1.0000 |
| | $s$ | day$^{-1}$ | 0.78 | 0.08 | 0.72 | 0.77 | 0.83 | 1.0001 |
| | $\lambda$ | day$^{-1}$ | 0.86 | 0.10 | 0.78 | 0.85 | 0.92 | 1.0001 |
| WM793b | $R_o(0)$ | µm | 188.81 | 3.09 | 186.75 | 188.82 | 190.89 | 1.0000 |
| | $R_c$ | µm | 189.56 | 3.11 | 187.48 | 189.56 | 191.65 | 1.0001 |
| | $\mathcal{R}$ | µm | 141.52 | 4.66 | 138.44 | 141.60 | 144.69 | 1.0001 |
| | $s$ | day$^{-1}$ | 0.12 | 0.02 | 0.11 | 0.12 | 0.14 | 1.0002 |
| | $\lambda$ | day$^{-1}$ | 0.54 | 0.40 | 0.23 | 0.46 | 0.76 | 1.0001 |
| WM164 | $R_o(0)$ | µm | 216.16 | 3.88 | 213.15 | 215.20 | 218.17 | 1.0001 |
| | $R_c$ | µm | 242.14 | 6.68 | 237.52 | 241.82 | 246.40 | 1.0001 |
| | $\mathcal{R}$ | µm | 217.25 | 4.49 | 213.93 | 216.45 | 219.75 | 1.0001 |
| | $s$ | day$^{-1}$ | 0.62 | 0.07 | 0.57 | 0.61 | 0.66 | 1.0002 |
| | $\lambda$ | day$^{-1}$ | 1.27 | 0.31 | 1.06 | 1.23 | 1.43 | 1.0003 |

Table S3: Greenspan's model parameters and MCMC diagnostics for hypoxia experiments. Summary statistics of the MCMC chains include: mean; standard deviation, $\sigma$; and, 25%, 50%, and 75% quartiles, $Q_{25\%}$, $Q_{50\%}$, and $Q_{75\%}$, respectively. To assess convergence of the MCMC chains we compute the potential scale reduction factor, $\hat{R}$, [9] where convergence corresponds to $\hat{R} < 1.1$.

| Cell line | Parameter | Units | Mean | $\sigma$ | $Q_{25\%}$ | $Q_{50\%}$ | $Q_{75\%}$ | $\hat{R}$ |
|---|---|---|---|---|---|---|---|---|
| WM983b | $\alpha_n$ | $m^3$ $kg^{-1}$ $s^{-1}$ $\times 10^{-7}$ | 13.87 | 0.89 | 13.27 | 13.88 | 14.47 | 1.0203 |
| | $\alpha_h$ | $m^3$ $kg^{-1}$ $s^{-1}$ $\times 10^{-7}$ | 3.12 | 0.18 | 2.99 | 3.10 | 3.22 | 1.0688 |
| | $\tau_\alpha$ | days | 0.14 | 0.10 | 0.06 | 0.13 | 0.23 | 1.0100 |
| | $\mathcal{R}_n$ | µm | 216.75 | 2.78 | 214.87 | 216.74 | 218.62 | 1.0103 |
| | $\mathcal{R}_h$ | µm | 130.42 | 3.82 | 128.21 | 130.74 | 132.98 | 1.2136 |
| | $\tau_{\mathcal{R}}$ | days | 0.85 | 0.12 | 0.79 | 0.87 | 0.94 | 1.0235 |
| | $s_n$ | $day^{-1}$ | 0.30 | 0.05 | 0.26 | 0.30 | 0.34 | 1.0015 |
| | $s_h$ | $day^{-1}$ | 0.84 | 0.17 | 0.72 | 0.84 | 0.97 | 1.0033 |
| | $\tau_s$ | days | 5.74 | 2.40 | 3.88 | 5.74 | 7.72 | 1.0726 |
| | $\lambda_n$ | $day^{-1}$ | 0.35 | 0.17 | 0.24 | 0.34 | 0.44 | 1.0803 |
| | $\lambda_h$ | $day^{-1}$ | 0.65 | 0.17 | 0.52 | 0.63 | 0.76 | 1.0097 |
| | $\tau_\lambda$ | days | 5.15 | 2.86 | 2.62 | 5.19 | 7.64 | 1.0098 |
| | $\hat{\lambda}$ | $day^{-1}$ | 5.91 | 2.48 | 3.87 | 6.03 | 8.05 | 1.0280 |
| | $\tau_{\hat{\lambda}}$ | days | 5.29 | 3.04 | 2.60 | 5.55 | 8.00 | 1.0331 |
| | $R_o(0)$ | µm | 217.12 | 2.77 | 215.25 | 217.10 | 218.99 | 1.0100 |
| WM793b | $\alpha_n$ | $m^3$ $kg^{-1}$ $s^{-1}$ $\times 10^{-7}$ | 19.56 | 1.24 | 18.74 | 19.56 | 20.39 | 1.0059 |
| | $\alpha_h$ | $m^3$ $kg^{-1}$ $s^{-1}$ $\times 10^{-7}$ | 2.19 | 0.21 | 2.04 | 2.15 | 2.28 | 1.1348 |
| | $\tau_\alpha$ | days | 0.34 | 0.10 | 0.27 | 0.32 | 0.41 | 1.1169 |
| | $\mathcal{R}_n$ | µm | 224.19 | 22.33 | 199.14 | 231.31 | 245.45 | 1.9616 |
| | $\mathcal{R}_h$ | µm | 127.90 | 3.58 | 125.46 | 127.90 | 130.40 | 1.0352 |
| | $\tau_{\mathcal{R}}$ | days | 0.16 | 0.08 | 0.09 | 0.15 | 0.22 | 1.0169 |
| | $s_n$ | $day^{-1}$ | 0.23 | 0.03 | 0.20 | 0.23 | 0.25 | 1.0149 |
| | $s_h$ | $day^{-1}$ | 0.11 | 0.05 | 0.08 | 0.11 | 0.14 | 1.0122 |
| | $\tau_s$ | days | 4.17 | 2.86 | 1.64 | 3.61 | 6.57 | 1.0550 |
| | $\lambda_n$ | $day^{-1}$ | 0.50 | 1.01 | 0.09 | 0.20 | 0.39 | 1.2686 |
| | $\lambda_h$ | $day^{-1}$ | 0.42 | 0.26 | 0.23 | 0.38 | 0.57 | 1.0214 |
| | $\tau_\lambda$ | days | 4.63 | 3.14 | 1.70 | 4.58 | 7.44 | 1.3675 |
| | $\hat{\lambda}$ | $day^{-1}$ | 1.26 | 2.28 | 0.19 | 0.27 | 0.61 | 1.1038 |
| | $\tau_{\hat{\lambda}}$ | days | 5.88 | 2.53 | 3.88 | 6.07 | 8.04 | 1.0354 |
| | $R_o(0)$ | µm | 194.57 | 3.56 | 192.12 | 194.47 | 197.01 | 1.0422 |
| WM164 | $\alpha_n$ | $m^3$ $kg^{-1}$ $s^{-1}$ $\times 10^{-7}$ | 7.60 | 0.82 | 7.05 | 7.61 | 8.15 | 1.0026 |
| | $\alpha_h$ | $m^3$ $kg^{-1}$ $s^{-1}$ $\times 10^{-7}$ | 1.47 | 0.30 | 1.26 | 1.42 | 1.63 | 1.0144 |
| | $\tau_\alpha$ | days | 0.83 | 0.24 | 0.68 | 0.85 | 1.00 | 1.0283 |
| | $\mathcal{R}_n$ | µm | 334.82 | 45.12 | 297.01 | 326.00 | 366.03 | 1.0036 |
| | $\mathcal{R}_h$ | µm | 149.50 | 12.73 | 141.70 | 150.45 | 158.31 | 1.0102 |
| | $\tau_{\mathcal{R}}$ | days | 0.35 | 0.22 | 0.17 | 0.34 | 0.49 | 1.0066 |
| | $s_n$ | $day^{-1}$ | 0.39 | 0.07 | 0.34 | 0.39 | 0.44 | 1.0019 |
| | $s_h$ | $day^{-1}$ | 0.60 | 0.17 | 0.47 | 0.60 | 0.73 | 1.0094 |
| | $\tau_s$ | days | 4.24 | 3.00 | 1.42 | 3.94 | 6.79 | 1.0073 |
| | $\lambda_n$ | $day^{-1}$ | 0.11 | 0.08 | 0.05 | 0.09 | 0.16 | 1.0041 |
| | $\lambda_h$ | $day^{-1}$ | 1.31 | 0.49 | 0.96 | 1.26 | 1.62 | 1.0025 |
| | $\tau_\lambda$ | days | 7.16 | 1.98 | 5.81 | 7.48 | 8.79 | 1.0034 |
| | $\hat{\lambda}$ | $day^{-1}$ | 1.86 | 2.52 | 0.34 | 0.50 | 2.38 | 1.0790 |
| | $\tau_{\hat{\lambda}}$ | days | 5.94 | 2.67 | 3.90 | 6.27 | 8.23 | 1.0031 |
| | $R_o(0)$ | µm | 281.63 | 7.99 | 276.23 | 281.35 | 286.68 | 1.0143 |

Table S4: Deoxygenation model parameters and MCMC diagnostics for deoxygenation experiments. Summary statistics of the MCMC chains include: mean; standard deviation, $\sigma$; and, 25%, 50%, and 75% quartiles, $Q_{25\%}$, $Q_{50\%}$, and $Q_{75\%}$, respectively. To assess convergence of the MCMC chains we compute the potential scale reduction factor, $\hat{R}$, [9] where convergence corresponds to $\hat{R} < 1.1$.

| Cell line | Parameter | Units | Mean | $\sigma$ | $Q_{25\%}$ | $Q_{50\%}$ | $Q_{75\%}$ | $\hat{R}$ |
|---|---|---|---|---|---|---|---|---|
| WM793b | $\alpha_n$ | m$^3$ kg$^{-1}$ s$^{-1}$ $\times 10^{-7}$ | 11.37 | 0.39 | 11.11 | 11.37 | 11.63 | 1.0059 |
| | $\alpha_h$ | m$^3$ kg$^{-1}$ s$^{-1}$ $\times 10^{-7}$ | 1.99 | 0.83 | 1.33 | 1.87 | 2.49 | 1.0209 |
| | $\tau_\alpha$ | days | 0.21 | 0.12 | 0.10 | 0.20 | 0.30 | 1.0018 |
| | $\mathcal{R}_n$ | µm | 258.07 | 4.04 | 255.20 | 258.06 | 260.76 | 1.1336 |
| | $\mathcal{R}_h$ | µm | 129.35 | 2.06 | 127.71 | 128.79 | 130.50 | 1.1404 |
| | $\tau_\mathcal{R}$ | days | 2.35 | 0.15 | 2.24 | 2.34 | 2.44 | 1.0966 |
| | $s_n$ | day$^{-1}$ | 0.29 | 0.01 | 0.28 | 0.29 | 0.29 | 1.0049 |
| | $s_h$ | day$^{-1}$ | 0.03 | 0.02 | 0.02 | 0.03 | 0.04 | 1.0036 |
| | $\tau_s$ | days | 2.83 | 0.33 | 2.61 | 2.81 | 3.03 | 1.0085 |
| | $\lambda_n$ | day$^{-1}$ | 2.69 | 2.19 | 0.88 | 2.18 | 3.99 | 1.0704 |
| | $\lambda_h$ | day$^{-1}$ | 0.97 | 0.62 | 0.46 | 0.90 | 1.42 | 1.0220 |
| | $\tau_\lambda$ | days | 3.57 | 2.72 | 1.23 | 3.02 | 5.51 | 1.1816 |
| | $\hat{\lambda}$ | day$^{-1}$ | 1.04 | 0.84 | 0.37 | 0.85 | 1.48 | 1.0552 |
| | $\tau_{\hat{\lambda}}$ | days | 6.56 | 1.91 | 5.06 | 6.56 | 8.13 | 1.0834 |
| | $\nu$ | - | 0.05 | 0.08 | 0.01 | 0.02 | 0.05 | 1.0226 |
| | $R_\mathrm{o}(t_\mathrm{s})$ | µm | 194.33 | 1.57 | 193.28 | 194.37 | 195.39 | 1.0345 |
| | $R_\mathrm{n}(t_\mathrm{s})$ | µm | 0.45 | 0.28 | 0.21 | 0.42 | 0.67 | 1.0131 |
| WM164 | $\alpha_n$ | m$^3$ kg$^{-1}$ s$^{-1}$ $\times 10^{-7}$ | 6.11 | 0.56 | 5.74 | 5.99 | 6.34 | 1.0520 |
| | $\alpha_h$ | m$^3$ kg$^{-1}$ s$^{-1}$ $\times 10^{-7}$ | 1.92 | 1.16 | 1.05 | 1.67 | 2.48 | 1.1873 |
| | $\tau_\alpha$ | days | 0.23 | 0.14 | 0.12 | 0.22 | 0.34 | 1.0035 |
| | $\mathcal{R}_n$ | µm | 322.27 | 78.90 | 273.33 | 325.23 | 372.46 | 1.0656 |
| | $\mathcal{R}_h$ | µm | 260.34 | 3.45 | 258.71 | 261.49 | 262.95 | 1.0230 |
| | $\tau_\mathcal{R}$ | days | 76.46 | 49.80 | 35.47 | 70.63 | 114.93 | 1.1486 |
| | $s_n$ | day$^{-1}$ | 0.47 | 0.03 | 0.45 | 0.48 | 0.50 | 1.0183 |
| | $s_h$ | day$^{-1}$ | 0.33 | 0.05 | 0.28 | 0.32 | 0.37 | 1.0128 |
| | $\tau_s$ | days | 2.74 | 2.07 | 1.40 | 2.01 | 3.29 | 1.0322 |
| | $\lambda_n$ | day$^{-1}$ | 0.24 | 0.15 | 0.11 | 0.22 | 0.35 | 1.0053 |
| | $\lambda_h$ | day$^{-1}$ | 1.31 | 0.69 | 0.76 | 1.28 | 1.83 | 1.0256 |
| | $\tau_\lambda$ | days | 4.83 | 2.95 | 2.19 | 4.72 | 7.43 | 1.0292 |
| | $\hat{\lambda}$ | day$^{-1}$ | 3.80 | 2.94 | 1.16 | 2.97 | 6.22 | 1.0474 |
| | $\tau_{\hat{\lambda}}$ | days | 5.52 | 2.85 | 3.19 | 5.63 | 8.04 | 1.0978 |
| | $\nu$ | - | 0.34 | 0.25 | 0.13 | 0.28 | 0.51 | 1.0276 |
| | $R_\mathrm{o}(t_\mathrm{s})$ | µm | 227.64 | 3.29 | 225.10 | 226.66 | 229.26 | 1.1002 |
| | $R_\mathrm{n}(t_\mathrm{s})$ | µm | 0.47 | 0.28 | 0.22 | 0.46 | 0.71 | 1.0027 |

Table S5: Re-oxygenation model parameters and MCMC diagnostics for re-oxygenation experiments with $t_\mathrm{s} = 2$ [days]. Summary statistics of the MCMC chains include: mean; standard deviation, $\sigma$; and, 25%, 50%, and 75% quartiles, $Q_{25\%}$, $Q_{50\%}$, and $Q_{75\%}$, respectively. To assess convergence of the MCMC chains we compute the potential scale reduction factor, $\hat{R}$, [9] where convergence corresponds to $\hat{R} < 1.1$.

| Cell line | Parameter | Units | Mean | $\sigma$ | $Q_{25\%}$ | $Q_{50\%}$ | $Q_{75\%}$ | $\hat{R}$ |
|---|---|---|---|---|---|---|---|---|
| WM793b | $\alpha_n$ | m$^3$ kg$^{-1}$ s$^{-1}$ $\times 10^{-7}$ | 11.27 | 0.96 | 10.51 | 11.26 | 11.98 | 1.0067 |
| | $\alpha_h$ | m$^3$ kg$^{-1}$ s$^{-1}$ $\times 10^{-7}$ | 1.59 | 0.05 | 1.56 | 1.59 | 1.63 | 1.0025 |
| | $\tau_\alpha$ | days | 0.36 | 0.22 | 0.18 | 0.34 | 0.52 | 1.0017 |
| | $\mathcal{R}_n$ | µm | 262.17 | 10.02 | 254.69 | 261.55 | 270.22 | 1.1506 |
| | $\mathcal{R}_h$ | µm | 142.79 | 4.72 | 139.56 | 142.90 | 146.07 | 1.0085 |
| | $\tau_{\mathcal{R}}$ | days | 2.11 | 0.31 | 1.89 | 2.09 | 2.32 | 1.1306 |
| | $s_n$ | day$^{-1}$ | 0.23 | 0.03 | 0.21 | 0.23 | 0.26 | 1.0317 |
| | $s_h$ | day$^{-1}$ | 0.07 | 0.02 | 0.05 | 0.07 | 0.08 | 1.0268 |
| | $\tau_s$ | days | 1.89 | 1.23 | 1.02 | 1.68 | 2.49 | 1.0287 |
| | $\lambda_n$ | day$^{-1}$ | 5.22 | 2.52 | 3.29 | 5.26 | 7.22 | 1.0127 |
| | $\lambda_h$ | day$^{-1}$ | 1.19 | 0.68 | 0.62 | 1.17 | 1.73 | 1.0025 |
| | $\tau_\lambda$ | days | 4.52 | 2.82 | 2.07 | 4.31 | 6.85 | 1.0183 |
| | $\hat{\lambda}$ | day$^{-1}$ | 2.75 | 2.47 | 1.01 | 1.75 | 3.64 | 1.0061 |
| | $\tau_{\hat{\lambda}}$ | days | 6.56 | 2.47 | 4.86 | 7.01 | 8.64 | 1.0190 |
| | $\nu$ | - | 0.48 | 0.28 | 0.24 | 0.48 | 0.72 | 1.0044 |
| | $R_o(0)$ | µm | 193.59 | 3.05 | 191.58 | 193.60 | 195.56 | 1.0028 |
| | $R_n(0)$ | µm | 0.46 | 0.28 | 0.22 | 0.45 | 0.69 | 1.0008 |
| WM164 | $\alpha_n$ | m$^3$ kg$^{-1}$ s$^{-1}$ $\times 10^{-7}$ | 7.12 | 0.28 | 6.93 | 7.11 | 7.30 | 1.0091 |
| | $\alpha_h$ | m$^3$ kg$^{-1}$ s$^{-1}$ $\times 10^{-7}$ | 1.04 | 0.04 | 1.01 | 1.04 | 1.07 | 1.0010 |
| | $\tau_\alpha$ | days | 0.29 | 0.18 | 0.15 | 0.27 | 0.42 | 1.0021 |
| | $\mathcal{R}_n$ | µm | 232.45 | 41.50 | 207.60 | 236.70 | 256.54 | 1.0212 |
| | $\mathcal{R}_h$ | µm | 226.81 | 2.80 | 224.85 | 226.36 | 228.31 | 1.0040 |
| | $\tau_{\mathcal{R}}$ | days | 23.99 | 15.19 | 11.50 | 22.70 | 35.18 | 1.0386 |
| | $s_n$ | day$^{-1}$ | 0.29 | 0.03 | 0.27 | 0.28 | 0.31 | 1.0025 |
| | $s_h$ | day$^{-1}$ | 0.46 | 0.07 | 0.40 | 0.45 | 0.50 | 1.0065 |
| | $\tau_s$ | days | 0.85 | 1.77 | 0.07 | 0.18 | 0.52 | 1.0166 |
| | $\lambda_n$ | day$^{-1}$ | 0.23 | 0.15 | 0.11 | 0.21 | 0.34 | 1.0008 |
| | $\lambda_h$ | day$^{-1}$ | 1.09 | 0.68 | 0.52 | 1.00 | 1.59 | 1.0014 |
| | $\tau_\lambda$ | days | 4.41 | 2.96 | 1.72 | 4.11 | 6.97 | 1.0023 |
| | $\hat{\lambda}$ | day$^{-1}$ | 5.26 | 2.63 | 3.04 | 5.18 | 7.48 | 1.0075 |
| | $\tau_{\hat{\lambda}}$ | days | 5.13 | 2.87 | 2.68 | 5.23 | 7.62 | 1.0027 |
| | $\nu$ | - | 0.22 | 0.22 | 0.06 | 0.15 | 0.32 | 1.0033 |
| | $R_o(0)$ | µm | 226.23 | 2.10 | 224.65 | 225.64 | 227.21 | 1.0087 |
| | $R_n(0)$ | µm | 0.47 | 0.28 | 0.22 | 0.45 | 0.70 | 1.0020 |

Table S6: Re-oxygenation model parameters and MCMC diagnostics for re-oxygenation experiments with $t_s = 4$ [days]. Summary statistics of the MCMC chains include: mean; standard deviation, $\sigma$; and, 25%, 50%, and 75% quartiles, $Q_{25\%}$, $Q_{50\%}$, and $Q_{75\%}$, respectively. To assess convergence of the MCMC chains we compute the potential scale reduction factor, $\hat{R}$, [9] where convergence corresponds to $\hat{R} < 1.1$.

### H Supplementary Movie Descriptions

#### H.1 Movie S1

Necrotic core movement and removal in WM983b spheroids in response to re-oxygenation. Time-lapse brightfield microscopy movie for three days following re-oxygenation at $t_s = 2.5$ [days]. The necrotic core of the spheroid is initially visible as a dark central region. As time progresses the necrotic core is located closer to the edge of the spheroid and the symmetric internal structure is lost. At later times the necrotic core appears to exit the spheroid as a single object.

#### H.2 Movie S2

Necrotic core movement in WM983b spheroids in response to re-oxygenation. Time-lapse brightfield microscopy movie for seven days following re-oxygenation at $t_s = 5.5$ [days]. The necrotic core of the spheroid is initially visible as a dark central region. At later times the necrotic core is close to the edge of the spheroid but does not exit as a single object. As the spheroid grows necrotic matter forms at the centre of the spheroid and appears to merge with the necrotic matter located closer to the periphery.

#### H.3 Movie S3

Loss of the necrotic core in WM164 re-oxygenation experiments. Time-lapse brightfield microscopy movie for three days following re-oxygenation at $t_s = 2.5$ [days]. The necrotic core of the spheroid is initially visible as a dark central region. As time progresses mass from the necrotic core at the centre of the spheroid moves towards the periphery and exits the spheroid. The spheroid then appears to resume growth.

### Supplementary References


[1] Browning, A. P. & Murphy, R. J. Image processing algorithm to identify structure of tumour spheroids with cell cycle labelling. *Zenodo* (2021). `https://doi.org/10.5281/zenodo.5121093`.

[2] Murphy, R. J., Browning, A. P., Gunasingh, G., Haass, N. K. & Simpson, M. J. Designing and interpreting 4D tumour spheroid experiments. *Communications Biology* **5**, 91 (2022).

[3] Browning, A. P. *et al.* Quantitative analysis of tumour spheroid structure. *eLife* **10**, e73020 (2021).

[4] Klowss, J. J. *et al.* A stochastic mathematical model of 4D tumour spheroids with real-time fluorescent cell cycle labelling. *Journal of the Royal Society Interface* **19**, 20210903 (2022).

[5] Greenspan, H. P. Models for the growth of a solid tumor by diffusion. *Studies in Applied Mathematics* **51**, 317–340 (1972).

[6] Bader, S. B., Dewhirst, M. W. & Hammond, E. M. Cyclic hypoxia: An update on its characteristics, methods to measure it and biological implications in cancer. *Cancers* **13**, 23 (2021).

[7] Grimes, D. R., Kelly, C., Bloch, K. & Partridge, M. A method for estimating the oxygen consumption rate in multicellular tumour spheroids. *Journal of the Royal Society Interface* **11**, 20131124 (2014).

[8] Gomes, A. *et al.* Oxygen partial pressure is a rate-limiting parameter for cell proliferation in 3D spheroids grown in physioxic culture condition. *PLoS One* **11**, e0161239 (2016).

[9] Gelman, A. *et al.* *Bayesian Data Analysis* (Chapman and Hall/CRC, New York, 2013), 3 edn.
